# The influence of biosecurity on the diversity and management of a 120-year old joint insect-fungus pest invasion

**DOI:** 10.1101/2025.04.12.647806

**Authors:** Firehiwot B. Eshetu, Irene Barnes, Helen F. Nahrung, Katrin N.E. Fitza, Bernard Slippers

## Abstract

The woodwasp, *Sirex noctilio*, and its mutualistic fungal symbiont, *Amylostereum areolatum*, are native to Eurasia and northern Africa. *Sirex noctilio* was first reported outside its native range in New Zealand in 1900, Tasmania in 1952 and mainland Australia in 1961. In this study, we consider the invasion history of these organisms across Australasia through population genetic analysis using mitochondrial sequence data and microsatellite markers and compared them with a previously published dataset from global collections. The study included contemporary (n=461) and historical (n=41) samples of *S. noctilio* dating back to 1952 and fungal (n=176) samples from across the range. No population structure was found in Australian and New Zealand populations of *S. noctilio* or the fungal symbiont *A. areolatum* reflecting both the natural (within the countries) and human-assisted (between the countries) spread of these symbionts. The *S. noctilio* populations in these countries had lower genetic diversity than other populations sampled globally. *Amylostereum areolatum* populations from Australia and New Zealand clustered separately from all other countries and were highly clonal. While the results suggested multiple early introductions in these two countries, it also reflected an efficient recent quarantine system that isolated these populations and reduced their complexity compared to other parts of the world. The findings also have relevance to the application of biological control for the pest complex.

## Introduction

The rise in international trade has increased the number of important invasive forest insects globally (Hurley et al. 2016; Nahrung and Carnegie 2020; Turner et al. 2021; Fenn-Moltu et al. 2022). The woodwasp *Sirex noctilio* Fabricius (Hymenoptera: Siricidae) and its symbiotic homobasidiomycetes fungus *Amylostereum areolatum* (Chaillet ex Fr.) Boidin (Russulales: Amylostereaceae) are native to Eurasia and northern Africa (Spradbery and Kirk 1978; Slippers et al. 2015). In these regions, *S. noctilio* is of minor economic importance and is not considered a pest of vigorous pine trees, with wasp attacks generally confined to highly stressed and already dying conifer trees (Wermelinger and Thomsen 2012). The situation is different in non-native environments, where extensive woodwasp outbreaks have occurred in large plantation-grown pine areas (Haugen 1990; Carnegie and Bashford 2012), and even mature and healthy trees have been severely attacked (Hurley et al. 2007; Haugen 1990).

In its non-native distribution in the Southern Hemisphere, it has been more than 124 years since the *Sirex-Amylostereum* complex established, with the earliest records from New Zealand (Miller and Clark 1935; Rawlings 1948). It was first detected in the southeastern part of the North Island (Wairarapa) in 1900 in a *Pinus radiata* plantation (Miller and Clark 1935; Bain et al. 2012), but no serious outbreaks occurred until the late 1940s, in drought-stressed and overstocked trees (Rawlings and Wilson 1949; Morgan and Stewart 1966). By the early 1950s, the wasp reached pine plantations on both Islands (Hurley et al. 2007; Bain et al. 2012), No serious outbreak occurred in these regions until the late 1940s, when *S. noctilio* invasion were identified in drought-stressed and overstocked trees (Rawlings and Wilson 1949; Morgan and Stewart 1966). No further serious outbreaks were reported in New Zealand and an integrated and effective management system is still practiced (Hurley et al. 2007; Bain et al. 2012).

In Australia, *S. noctilio* was first detected in Tasmania in 1952 (Gilbert and Miller 1952) and mainland Australia (Victoria) in 1961 (Collett and Elms 2009). Further spread within Australia followed pine-growing areas in South Australia, New South Wales, and Queensland between 1980 and 2009 (Carnegie et al. 2005; Collett and Elms 2009; Nahrung et al. 2015). In these areas, *S. noctilio* has been mainly a pest of *P. radiata* (the main species used in plantations), however, as the spread continues northward into Queensland it potentially threatens other pine species planted in the area (Nahrung et al. 2015). Australia experienced one of the most severe *S. noctilio* outbreaks in the Green Triangle Region (south-eastern South Australia and south-western Victoria) in 1987 which caused the death of over one million pine trees (Haugen 1990; Carnegie and Bashford 2012). By 2018, Sirex-related pine tree mortality and its management since the first arrival of the wasp have cost approximately AUD$35M (Cameron et al. 2018). Currently, the woodwasp population in Australia is well controlled using effective integrated management approaches (Carnegie and Bashford 2012), however, it is still a potential risk in the absence of these.

*Sirex noctilio* has shown the exceptional ability of a forest pest to spread and pose a risk to pine plantations globally (Ciesla 2003; Slippers et al. 2015; Ireland 2018). In South America, *S. noctilio* was detected in Uruguay (1980), Argentina (1985), Brazil (1988), and Chile (2001) (Corley et al. 2019). By 1994, *S. noctilio* was detected in the Cape province of South Africa and has since spread across the country (Tribe and Cillie 2004; Hurley et al. 2007; Mlonyeni et al. 2018). The expansion of *S. noctilio* reached the USA and Canada in 2004 and 2005, respectively (Ciesla 2003; Hoebeke et al. 2005; de Groot et al. 2006). Most recently, *S. noctilio* was detected in China in 2013 (Sun et al. 2016). The diversity of the host trees in North America and China - a mixture of introduced and native pines - differs from that of the Southern Hemisphere, where the commercial plantations are largely based on exotic *Pinus* spp. in uniform stands (Burgess and Wingfield 2001; Hurley et al. 2017). This diversity of tree species and native natural enemies often lowers the risk of *S. noctilio* related outbreaks (Yemshanov et al. 2009; Dodds et al. 2010; Ayres et al. 2014).

There has been significant interest in understanding the structure and diversity of *A. areolatum* and *S. noctilio* populations given their spread worldwide (Slippers et al. 2015; Sun et al. 2016). Before the enrichment of modern molecular tools, vegetative compatibility groups (VCGs) culture assays were performed to profile the genetic relatedness, structure and dispersal of *A. areolatum* populations (Thomsen and Koch 1999; Slippers et al. 2001), where it was important to understand the distribution of the clonal VCGs in northern Europe across large areas (Vasiliauskas et al.1998; Thomsen and Koch 1999; Vasiliauskas and Stenlid 1999). The clonality in these fungal populations is influenced by the prominence of its woodwasp-mediated vegetative dispersal over the dispersal of outcrossed basidiospores (Spradbery and Kirk 1978; Wermelinger and Thomsen 2012). A similar pattern of clonal VCG was noticed across introduced *A. areolatum* populations in South Africa and South America (Slippers et al. 2001). Slippers et al. (2001) also identified an additional and rare VCG linked to the fungus used in the biocontrol nematode mass-rearing program in Australia and South Africa. This rearing fungus is thought to have a separate European origin from *S. noctilio* field populations and later rearing fungal cultures (Nahrung 2017). Subsequent studies used VCG to supplement genetic diversity surveys and showed a similar pattern of clonality in the introduced fungus populations (Wooding et al. 2013; Fitza et al. 2016), although this approach lacks accuracy and barely distinguishes between closely related fungal clones (Mlonyeni et al. 2018).

Apart from VCGs, mitochondrial and nuclear sequence data, and microsatellite data have been used to understand the diversity of *A. areolatum* populations (Castrillo et al. 2015; Mlonyeni et al. 2018). Studies that applied the intergenic spacer (IGS) region of ribosomal RNA (nuclear IGS rRNA) identified two IGS sequence groups in the Southern Hemisphere and North America (Slippers et al. 2002; Nielsen et al. 2009; Bergeron et al. 2011), of which the dominant type overlaps between these regions. This aligns with the findings of Boissin et al. (2012), who identified two source populations for the global invasion of the wasp: the native European and an as-yet unsampled population. Two additional fungal IGS types have been recently reported from North America, showing the plausibility of multiple introductions from different sources (Castrillo et al. 2015). Mlonyeni et al. (2018) developed eleven microsatellite markers and investigated *A. areolatum* diversity in South Africa, where nine distinct multilocus genotypes were found spread across the country without any distinguishable population structure. Detailed studies on the population genetic diversity and structure from Australasia where the *Sirex*-*Amylostereum* invasion occurred first, are lacking.

Like *A. areolatum* populations, the *S. noctilio* global invasion has been traced using molecular tools, revealing complex patterns of spread (Boissin et al. 2012; Bittner et al. 2017). Boissin et al. (2012) assigned the Australasian *S. noctilio* population to a cluster that included populations from Europe, North America, South Africa, Uruguay, Argentina, and Chile, but the Australasian population had the lowest diversity of the invasive populations. More recent studies of the invasion history of the woodwasp and *A. areolatum* populations in North America identified complex and multiple introduction patterns (Castrillo et al. 2015; Bittner et al. 2017). These findings highlight why continuous monitoring of the *Sirex*-*Amylostereum* population is valuable.

Population genetics studies have created awareness of the complex introduction history of *S. noctilio*–*A. areolatum* based on independent investigations of each organism. The current study considers the insect-fungus joint invasion. We aimed to examine the impact of the invasion over 70 and 120 years in Australia and New Zealand, respectively - on the population genetic diversity, structure and history of these organisms in these Australasian countries. Specifically, we assessed whether the original incursion persists, or if subsequent introductions occurred. Lastly, the diversity of the woodwasp and fungal populations from Australasia was compared to those determined in previous global studies. Our collections allow us to consider the impact of quarantine systems to constrain multiple introductions, and how patterns of diversity could inform the current biological control management systems.

## Materials and methods

### Sampling of S. noctilio and A. areolatum

To represent all the invaded geographical regions, specimens of the woodwasp *Sirex noctilio* were obtained from five states in Australia (New South Wales, South Australia, Queensland, Tasmania, and Victoria) and New Zealand (the Northern and Southern Islands) (Fig. 1a, b). All insect specimens were preserved in absolute ethanol and kept frozen at −20°C until used for DNA extraction. To investigate the evolution and history of introductions of *S. noctilio* in Australia and New Zealand, our sampling also included dry specimens from the museum collection (University of Tasmania, 1952 to 2001) to the most recent collections (2006 to 2018; Supplementary Table S1). Specimens are deposited in the Forestry and Agricultural Biotechnology Institute (FABI, University of Pretoria, South Africa) at the insect specimen and DNA collection.

**Fig. 1.**
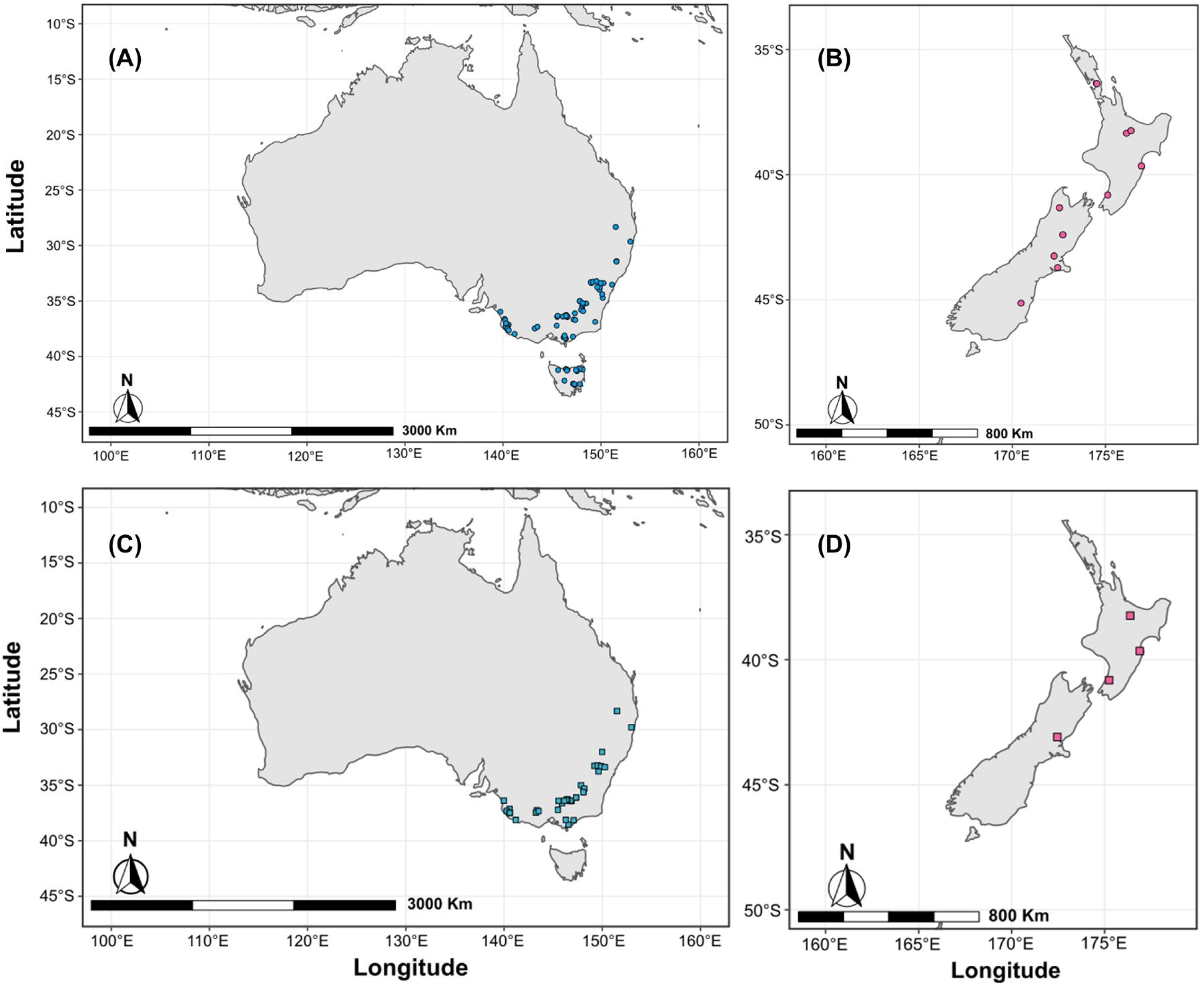
*Sirex noctilio* (A - B) and symbiotic fungus *A. areolatum* (C-D) collection sites. (A) Tasmania and four states (i.e., Victoria, New South Wales, South Australia, and Queensland) from mainland Australia, (B) South and North Island of New Zealand, (C) four states (i.e., Victoria, New South Wales, South Australia and Queensland) from mainland Australia, and (D) South and North Island of New Zealand.

Strains of *A. areolatum* were obtained from four states in Australia (New South Wales, Queensland, South Australia, and Victoria) and the Northern Island of New Zealand (Fig. 1c, d). Where there was limited access to live fungal cultures, *A. areolatum* was extracted from the mycangia of ethanol-preserved individual female *S. noctilio* (Supplementary Table S2). Strains of *A. areolatum* used to rear the Kamona strain of *D. siricidicola* (provided by EcoGrow, Australia) were included in this study. The global collection of the *A. areolatum* strains from the fungal culture collection (CMW) of FABI was included for comparison. The CMW strains include samples from Europe (Austria, Czech Republic, Denmark, Russia, and Spain), Australia, New Zealand, North America (Canada and USA) and South Africa (Supplementary Table S3). Cultures were grown on 1/2 Potato Extract Agar (PDA, 19.5 g l^-1^ potato dextrose extract, 19.5 g l^-1^ purified agar) until used for DNA extraction. All fungal cultures obtained in this study are maintained in the CMW culture collection of FABI.

### DNA extractions

#### The woodwasp *S. noctilio*

DNA was extracted from specimens preserved in absolute ethanol and pieces of dried specimens from museum collections. Ethanol-preserved specimens were rinsed thoroughly with sterilized distilled water and genomic DNA was either extracted from a portion of muscle taken from the thorax/abdomen junction or pieces taken from a single leg. In the case of dried museum specimens, a piece of the single leg was used. Genomic DNA from all specimen types was extracted using the NucleoSpin® Tissue XS (Macherey-Negel, Germany) a tissue DNA recovery kit following the manufacturer’s protocol. The concentration of the genomic DNA was quantified with a Nano-Drop ND-1000 UV/Vis Spectrometer (NanoDrop Technologies, Wilmington, DE 19810 USA) and adjusted to a final working concentration of 10 ng/µl for downstream PCR amplification.

#### The fungal symbiont *A. areolatum*

Genomic DNA was either extracted from the fresh mycelium of actively growing fungus on Petri dish plates containing PDA or from ethanol-preserved female *S. noctilio* mycangium. For the fresh mycelium growing in a Petri dish, mycelium collection and preparation were performed according to Slippers et al. (2001) and DNA was extracted using phenol-chloroform as described in Sambrook and Russell (2006). Genomic DNA precipitation, further cleaning and dilutions were performed as described by Fitza et al. (2016). The NucleoSpin® Tissue DNA extraction kit was used for genomic DNA extracted from ethanol-preserved female *S. noctilio* mycangia following the manufacturer’s instructions. The concentration of the genomic DNA harvested using both methods was quantified using the same method as above and the concentration was adjusted to a final working concentration of 30 ng/µl for downstream PCR amplification.

### Mitochondrial (mtCOI and mtSSU rRNA) amplifications and diversity analysis

#### Mitochondrial (mtCOI and mtSSU rRNA) amplifications and sequencing

A portion of the 5′ end of the general barcoding cytochrome c oxidase subunit I (COI) mitochondrial gene was amplified in all the *S. noctilio* specimens using the primers LiLCO1490: 50-ATT TGA TCT GGA ATT TTA GG30 (Dittrich-Schröder et al. 2018) and HCO2198 (C1N-2173): 50-TAA ACT TCA GGG TGA CCA AAA AAT CA-30 (Folmer et al. 1994). The PCR reaction mix was prepared in a 25 µl total volume using the MyTaq™ DNA polymerase protocol consisting of 0.5 µl of 1.5-unit MyTaq™ DNA polymerase (Bioline Ltd. UK), 5 µl of 10x MyTaq™ PCR buffer, 1µl of 0.5 M of each primer, 2 µl of template DNA (10 ng/µl) and 15.5 µl of sterilized PCR grade SABAX water. The thermocycling protocol included an initial denaturation of 3 min at 95 °C, 35 cycles of 45 s at 94 °C, 30 s at 56 °C and 1 min at 72 °C, followed by a final extension of 10 min at 72 °C. Gel electrophoresis was performed on 2 % (w/v) agarose using 3 µl of PCR product mixed with 2 µl GelRed^TM^ (Biotium, California) in a sodium-borate buffer system and visualized under ultraviolet light. PCR product purification, sequencing PCR, and precipitation were the same as those described by Fitza et al. (2019).

A portion of the 5′ end of the mitochondrial small subunit (mtSSU) ribosomal RNA gene was amplified in all the *A. areolatum* strains using the primers MS1 (forward: CAG CAG TCA AGA ATA TTA GTC AAT G) and MS2 (reverse: GCG GAT TAT CGA ATT AAA TAA C) (White 1990). The PCR reaction mixture and thermal cycler conditions were used as in Fitza et al. (2016) and the template DNA concentration was adjusted to 60 ng/µl. Gel electrophoresis, PCR products purification, sequencing PCR and precipitation were the same as those described by Eshetu et al. (2023).

Precipitated PCR products for both mtCOI gene of *S. noctilio* and mtSSU gene of *A. areolatum* were sent for sequencing at the DNA Sequencing Facility, Faculty of Natural and Agricultural Science, University of Pretoria, South Africa.

#### Haplotype analysis

Sequences generated both from the mtCOI gene region of *S. noctilio* and the mtSSU rRNA region of *A. areolatum* were edited and manually checked in Bioedit version 7.2.5 (Hall 1999) and aligned using the MAFFT free online alignment program version 7 (Katoh et al. 2017). A median-joining haplotype network was constructed to investigate the relationships among populations using NETWORK version 10.2.0.0. In the analyses, historical sequences were considered. For *S. noctilio*, mtCOI sequence data generated by Boissin et al. (2012), from specimens from Argentina, Australia, Chile, Europe, New Zealand, North America, South Africa, and Uruguay were included in the analyses. NCBI sequences were used for specimens from China (Sun et al. 2016).

For the symbiotic fungus, the mtSSU rRNA sequence used to generate the median-joining haplotype network in NETWORK included those generated for the strains from the CMW *A. areolatum* collection from both the native range in Europe (Austria, Czech Republic, Denmark, Spain, and Russia) and non-native regions (Australia, New Zealand and South Africa) (Supplementary Table S3).

### Amplification of microsatellite markers

To genotype the *S. noctilio* samples, 14 previously developed microsatellite markers (Santana et al. 2009) were used (Supplementary Table S4). PCR and the thermal cycler condition were conducted using QIAGEN Multiplex PCR master mix following the manufacturer’s protocol and as in Boissin et al. (2012). Genotyping of the fungal symbiont *A. areolatum* was performed using 11 previously developed microsatellite markers (Mlonyeni et al. 2018) (Supplementary Table S5). The PCR reaction mixture and thermal cycler conditions were the same as in Mlonyeni et al. (2018). The primer panel arrangements were the same as in Boissin et al. (2012) and Mlonyeni et al. (2018), for *S. noctilio* and *A. areolatum*, respectively. For fragment analyses, a 1:100 dilution of PCR amplicon with sterile SABAX water was made for all samples and pooled according to the panel arrangement above. LIZ500 (GeneScan™ 500 LIZ™ dye Size Standard, Thermo Fisher Scientific, Waltham, MA, USA) was used as the size standard marker. The GeneScan mix and denaturation conditions were as the same as in Eshetu et al. (2023). The samples (1 µl per lane) were run on the ABI PRISM^TM^ 3500xI DNA analyser to determine product size (DNA sequencing facility, University of Pretoria). The GeneScan data was analyzed using GeneMapper® v4.1 (Life Technologies, Foster City, CA) to score allele fragment sizes.

### Microsatellite analyses

#### Genetic diversity analyses

Various diversity indices were calculated for *S. noctilio* and *A. areolatum* using data generated in this study from Australia and New Zealand and the datasets from Boissin et al. (2012) and Mlonyeni et al. (2018). To accommodate for potential variation in alleles scored between experiments and different ABI machines used (ABI3100 in Boissin et al. (2012) and ABI PRISM^TM^ 3500xI in this study), DNA representing different haplotypes obtained for both wasp and fungi during these previous studies was re-screened in the current study to verify and standardize allele scoring. Where necessary, alleles were re-scored so that datasets could be combined. For *S. noctilio*, this was successfully done for 12 microsatellite markers. However, two loci (Sn507 and Sn231) did not show consistency in the conversion of their allelic size shift and were thus not included in the comparative analyses of global woodwasp populations. No shifts in allele sizes were observed for the *A. areolatum* samples and thus the data set of Mlonyeni et al. (2018) was integrated into the current study without modification.

Populations of the woodwasp and the fungus were defined based on the country of origin for further analysis. However, the European woodwasp collection was split based on the gene pool information as in Boissin et al. (2012): Europe-Pop1 (Czech Republic, Finland, France, Germany, Greece, Spain, and Russia) and Europe-Pop2 (Switzerland). The private allelic richness (P_a_) and allelic richness (A_R_) were calculated in Hp-rare version 1.1 Kalinowski (2005). The R (Team, 2013) package Poppr (Kamvar et al. 2014) was used to compute further genetic diversity indices, including the number of multilocus genotypes (MLG), numbers of expected multilocus genotypes based on rarefaction (eMLG) (Hurlbert 1971; Heck Jr. et al.1975), standard error (SE) based on eMLG rarefaction, MLG diversity (H) using Shannon-Wiener Index (Shannon 1948), MLG diversity (G) using Stoddart and Taylor’s index (Stoddart and Taylor 1988), Evenness (E5) (Grünwald et al. 2003) and Nei’s unbiased gene diversity (Hexp) (Nei 1978).

#### Population molecular variance, genetic differentiation and gene flow

To test the hypothesis of population differentiation among population and within a population, analysis of molecular variance (AMOVA) was conducted in R (Team, 2013) package Poppr (Kamvar et al. 2014) and populations were defined based on state (for within country) and country of origin (between country). Nei’s genetic distance and Nei’s unbiased genetic distance were used to estimate pairwise comparisons of genetic differentiation (DP_T_) and gene flow (Nm) using GenAlEx v 6.5 (Peakall and Smouse 2012). These estimates were conducted using both the *S. noctilio* and *A. areolatum* datasets.

#### Analysis of Minimum Spanning Network

To assess genetic relatedness amongst the observed multilocus genotypes (MLGs) for both the *S. noctilio* and *A. areolatum* populations, minimum spanning networks (MSN) were constructed using the Nei. dist function in R (Team 2013) package Poppr (Nei 1978; Kamvar et al. 2014). To determine the evolutionary history between observed MLGs of the fungus, and populations from Australia and New Zealand, the data set of Mlonyeni et al. (2018) and the isolates from this were compared. Two types of population clustering were considered to assess the relatedness of the *S. noctilio* populations: (i) geographic-based clustering on populations of Australia and New Zealand and the global collection from the data sets of Boissin et al. (2012), and (ii) collection time-based clustering on the datasets of the Australian, New Zealand and the museum specimens (>1952 – 2002) to investigate change over time in this population.

#### Population structure

A model-based Bayesian clustering algorithm was implemented in the STRUCTURE program v2.3.4 (Pritchard et al. 2000) to determine if there is a structure in populations of *S. noctilio* and *A. areolatum*. The parameters were set with the assumption of admixture (ancestry model), whereby individuals may have mixed ancestry and/or individuals have inherited a proportion of their genome from each of the K populations. The K values were tested from 1 to 10, each with 20 independent runs, 700 000 Markov Chain Monte Carlo (MCMC) iterations, and a burn-in of 100 000. The optimal cluster for K, that best fit the data was calculated in STRUCTURE using the Evanno method (Evanno et al. 2005), and the clusters were assessed and visualized in STRUCTURE HARVESTER (Earl and vonHoldt 2012) and CLUMPAK (http://clumpak.tau.ac.il/). Principal coordinate analysis (PCoA) was analyzed to compare the fungal population with a 999-permutation test using GenAlEx version 6.505 (Peakall and Smouse 2012) to further investigate the population sub-division without assuming Hardy-Weinberg equilibrium.

## Results

### Samples

For *S. noctilio*, a total of 461 contemporary specimens were obtained from five states in Australia (n = 94 from New South Wales; n = 53 from Queensland; n = 51 from South Australia; n = 69 from Tasmania; n = 133 from Victoria) and New Zealand (n = 45 from the North Island; n = 16 from the South Island; Supplementary Table S1). In addition, 41 *S. noctilio* historical (<2001) specimens were included from the museum collection in Australia (n = 6 from New South Wales; n = 2 from South Australia; n = 17 from Tasmania) and New Zealand (n = 5 from North Island; n = 11 from South Island; Supplementary Table S1).

A total of 143 *A. areolatum* isolates were obtained from the North Island of New Zealand (n = 25), and four states in Australia (n = 25 from New South Wales; n = 10 from Queensland; n = 10 from South Australia; n = 73 from Victoria; Supplementary Table S2). The three fungal isolates (CMW49355, CMW53289 and CMW53288) that have been used to mass-rear the biocontrol nematode Kamona strain of *D. siricidicola* provided by EcoGrow in Australia, were included in this study. Additionally, 30 CMW isolates of *A. areolatum* were included from introduced areas (n = 2 from Australia; n = 3 from Canada; n = 1 New Zealand; n = 1 South Africa; n = 3 from the USA) and native areas (n = 5 from Austria; n = 4 from Czech Republican; n= 2 from Denmark; n = 8 from Spain; n = 2 from Russia; Supplementary Table S2).

### Mitochondrial (mtCOI and mtSSU rRNA) amplifications and diversity analysis

#### Mitochondrial (mtCOI and mtSSU rRNA) amplification and sequencing

A 581 bp amplicon of the cytochrome oxidase I region (COI) of the mitochondrial DNA was sequenced for 64 (Australia, n = 53; New Zealand, n = 11) *S. noctilio* specimens.

A 530 bp amplicon of the mitochondrial small subunit (SSU) ribosomal RNA was sequenced for 107 (Australia, n = 87; New Zealand, n = 20) *A. areolatum* strains. The same 530 bp amplicon of mtSSU rRNA was sequenced for an additional 23 CMW *A. areolatum* global isolates from native (Austria, n = 5; Czech Republic, n = 3; Denmark, n=2; Russia, n = 2, Spain, n = 5) and non-native distributions (Australia, n =2; New Zealand, n=1; South Africa, n = 1; USA, n = 2; Supplementary Table S3).

### Haplotype network analysis

mtCOI sequences were used to construct a haplotype network of the woodwasp (Fig. 2). Overall, 17 haplotypes of *S. noctilio* were recovered from 243 sequences generated in this (n = 64) and previous studies available in GenBank (n = 179; Fig. 2). The reference sequences were obtained from previous studies and GenBank, representing the native (Switzerland, n = 28; other European countries (n = 8) and introduced range (Argentina, n = 16; Australia, n = 27; Chile, n = 22; China, n=16; North America, n=37; South Africa, n = 24; Uruguay, n = 1). Four haplotypes were present in Australia (H3, H4, H9 & H16) and two in New Zealand (H9 & H15) (Fig. 2). Haplotype H9 was the most common, represented by 150 samples from nine countries in the native and introduced ranges (Fig. 2). The Australian individuals represented 48.67 % of the total individuals in H9, while New Zealand individuals made up 6 %. Haplotype H9 represented 91.25 % and 81.82 % of the total numbers of individuals from Australia and New Zealand, respectively. Haplotype H16 represented 6.25 % of the total individuals in Australia and all individuals in this haplotype were sampled in the current study. Haplotypes H3 & H4 represented 1.25 % each of the total number of individuals in Australia and contained individuals only from previous studies. Haplotype H15 is present only in New Zealand and represents 18.18 % of the total number of individuals (Fig. 2). Except for the three haplotypes (H1, H8 & H9) that represent the native distribution, all other haplotypes represent individuals from only introduced regions.

**Fig. 2.**
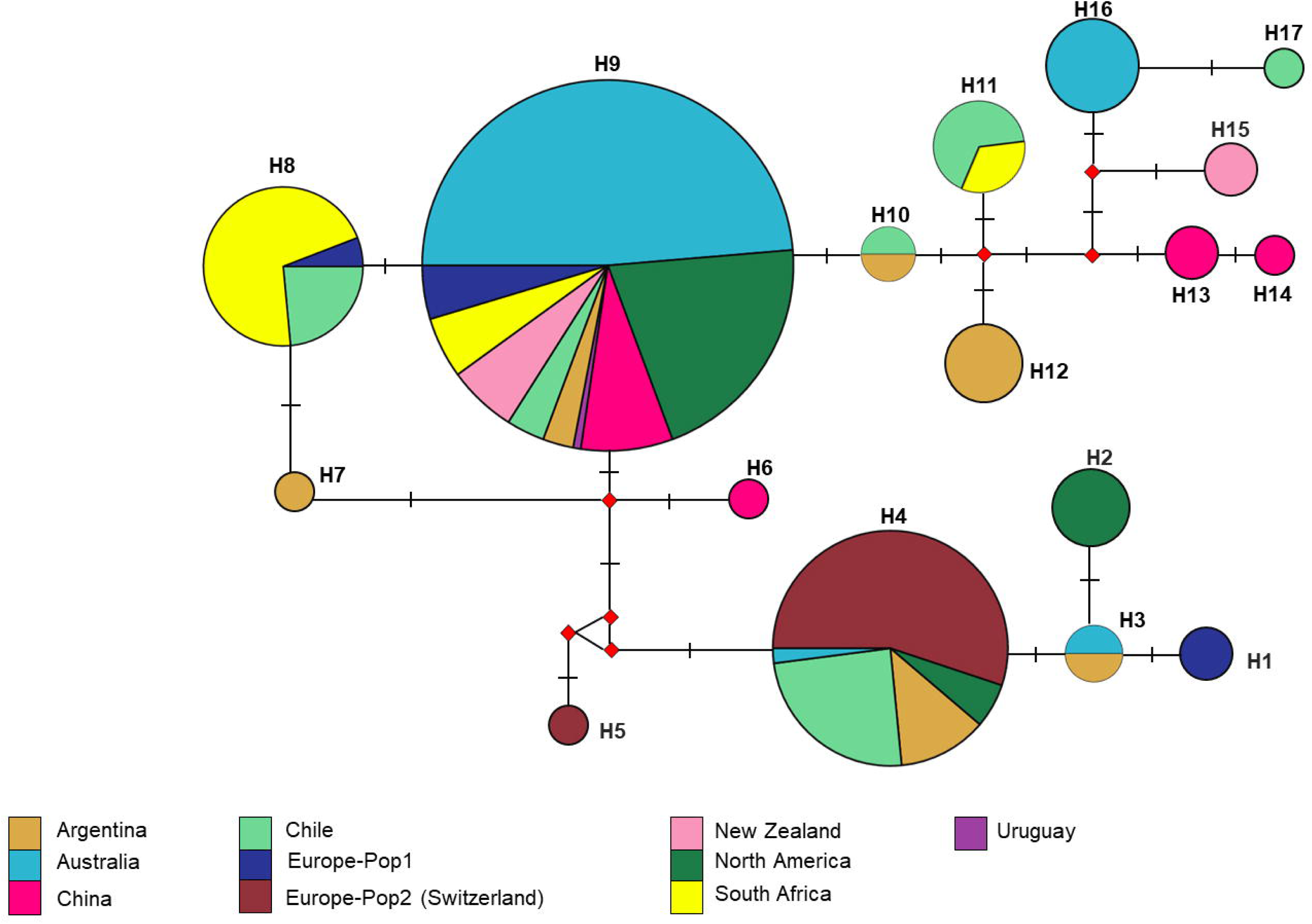
Mitochondrial cytochrome *c* oxidase subunit 1 (mtCOI) sequence-based median-joining haplotype network of *S. noctilio* populations. Each node represents specimens that share a similar mtCOI genetic profile, the size of the circles is proportional to the number of samples linked to the haplotype. and colors represent geographic origin.

mtSSU rRNA sequences were used to construct a haplotype network of *A. areolatum* populations (Fig. 3). Four haplotypes (H1-H4) of *A. areolatum* were recovered from 130 sequences generated from individuals in native and invasive regions (Fig. 3). H3 was dominant and was shared between populations in native and introduced regions. It represents 100 % of strains used in the mass-rearing program, in Australia and comprised 97.67 % and 100 % of the Australian and New Zealand isolates, respectively. This haplotype is also present in Austria, South Africa and North America, of which individuals from these countries all represent 2.6 % of the total individuals in H3. From the native distribution, only individuals from Austria present in both H1 and H3 (Fig. 3). H1 only represents populations from the native distribution except for individuals from Spain of which 100 % of the individuals are represented in H3 (Fig. 3).

**Fig. 3.**
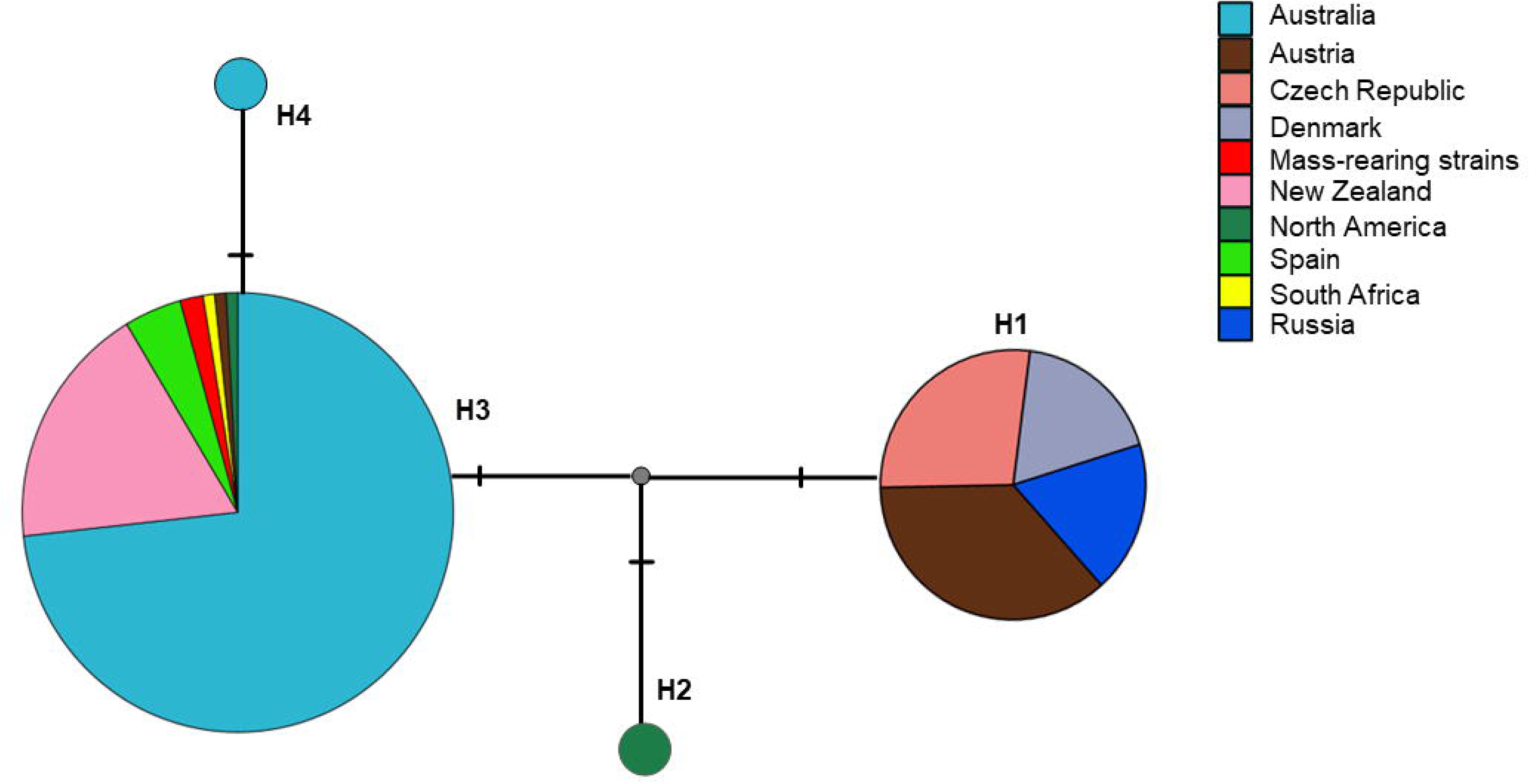
Mitochondrial small subunit (SSU) ribosomal RNA sequence-based median-joining haplotype network of *A. areolatum* populations. Each node represents specimens that share a similar mtSSU rRNA genetic profile and colours represent geographic origin.

### Microsatellite analyses

#### Genetic diversity analyses

For the *S. noctilio* sub-population in Australia, the number of observed alleles ranged from A_R_= 17 (South Australia) to A_R_= 21 (Victoria; Table 1). The allelic richness in Australia was the lowest (A_R_ = 22 and Pa = 1; Table 1) in all populations considered, with an average of 2.167 alleles per locus relative to other non-native populations from Argentina that had an average of 4.667 per locus (Supplementary Table S6). The allelic richness in the New Zealand population was higher than that in Australia (A_R_ = 30 and P_a_ = 2; Table 1), with an average of 2.667 alleles per locus (Supplementary Table S6). Overall, the highest allelic richness was observed in the native population (Europe-Pop1: A_R_ = 57 and P_a_ = 10), with an average of 4.75 alleles per locus.

**Table 1.**
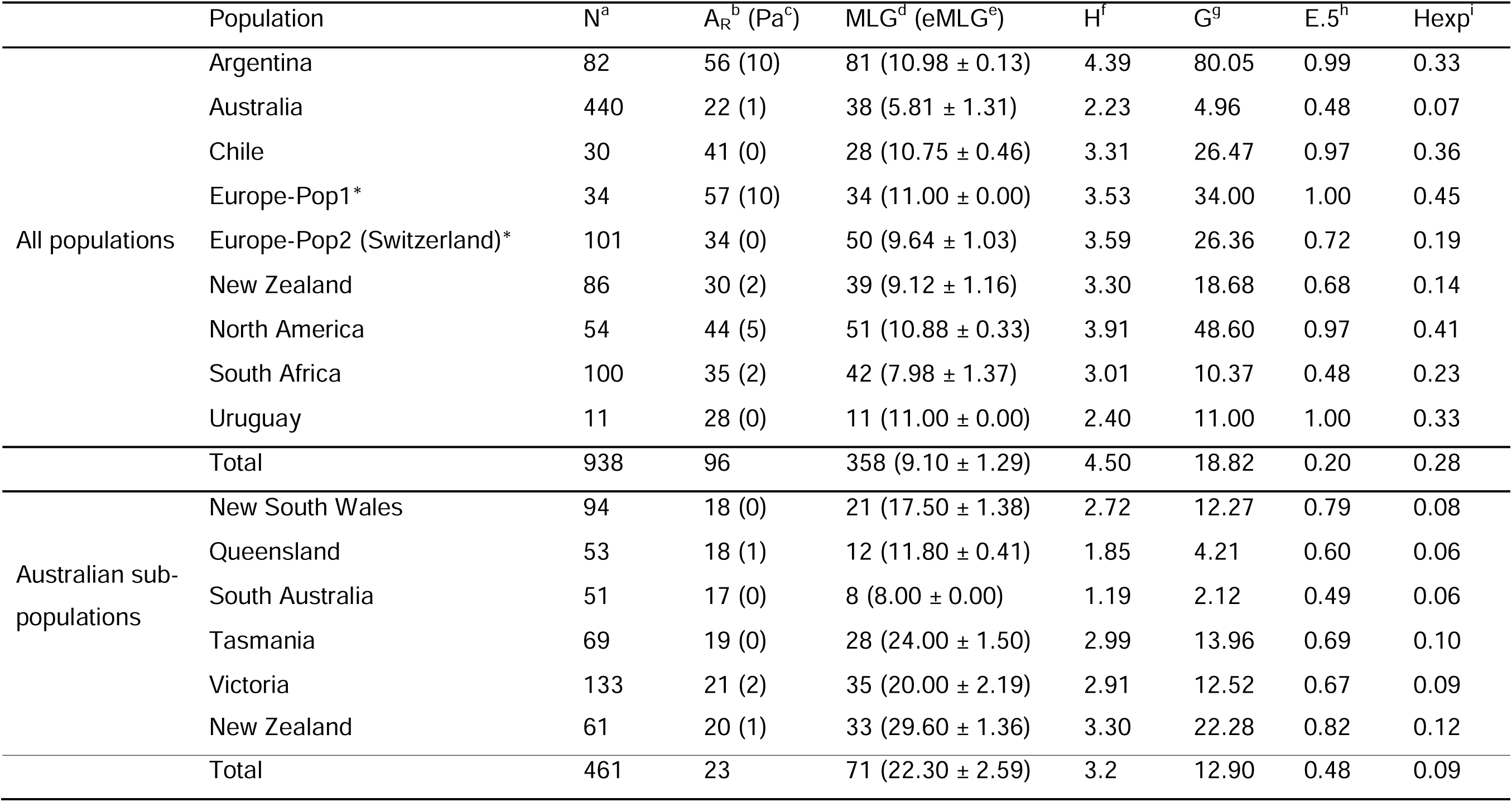
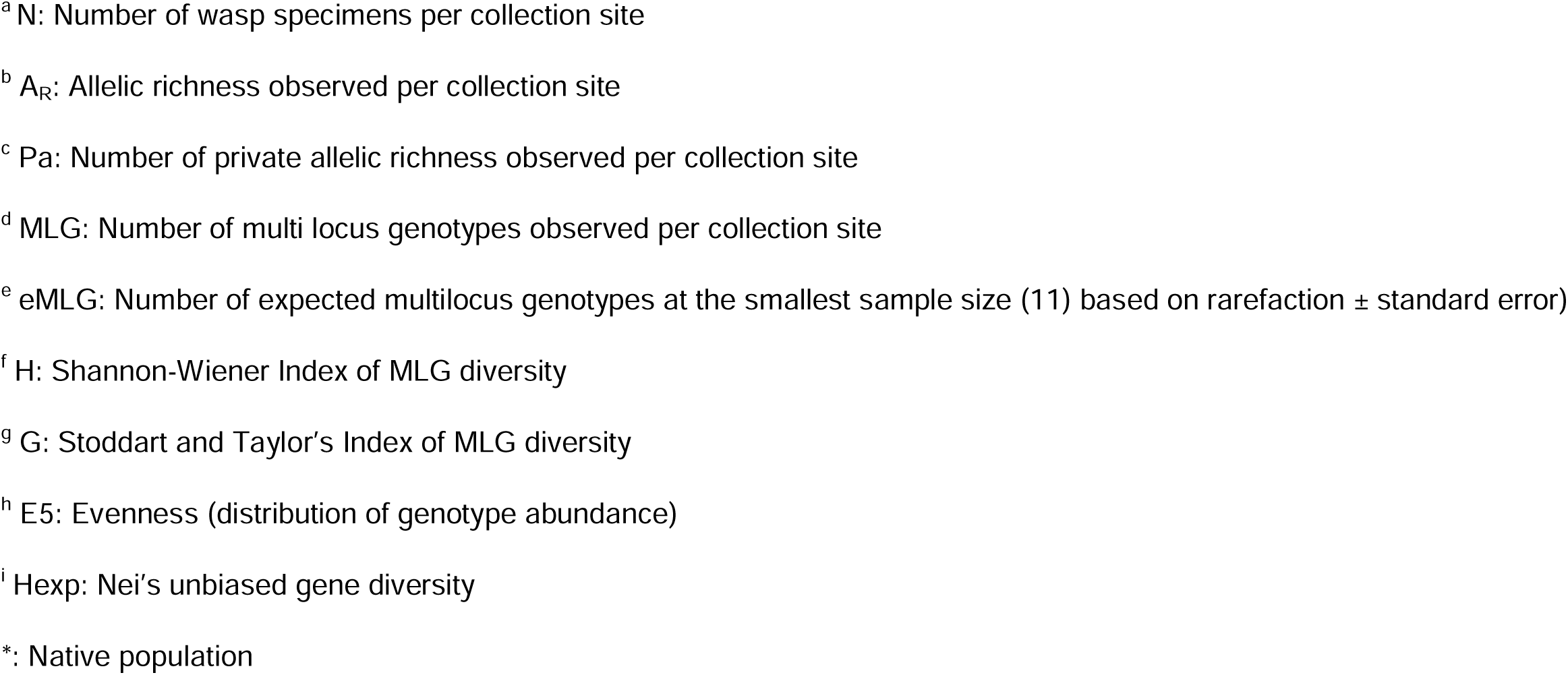
Summary of genetic diversity indices for *Sirex noctilio* populations from Boissin et al. (2012) and collections made in this study from Australia and New Zealand.

|  | Population | N<sup>a</sup> | A<sub>R</sub><sup>b</sup> (Pa<sup>c</sup>) | MLG<sup>d</sup> (eMLG<sup>e</sup>) | H<sup>f</sup> | G<sup>g</sup> | E.5<sup>h</sup> | Hexp<sup>i</sup> |
| --- | --- | --- | --- | --- | --- | --- | --- | --- |
| All populations | Argentina | 82 | 56 (10) | 81 (10.98 ± 0.13) | 4.39 | 80.05 | 0.99 | 0.33 |
|  | Australia | 440 | 22 (1) | 38 (5.81 ± 1.31) | 2.23 | 4.96 | 0.48 | 0.07 |
|  | Chile | 30 | 41 (0) | 28 (10.75 ± 0.46) | 3.31 | 26.47 | 0.97 | 0.36 |
|  | Europe-Pop1* | 34 | 57 (10) | 34 (11.00 ± 0.00) | 3.53 | 34.00 | 1.00 | 0.45 |
|  | Europe-Pop2 (Switzerland)* | 101 | 34 (0) | 50 (9.64 ± 1.03) | 3.59 | 26.36 | 0.72 | 0.19 |
|  | New Zealand | 86 | 30 (2) | 39 (9.12 ± 1.16) | 3.30 | 18.68 | 0.68 | 0.14 |
|  | North America | 54 | 44 (5) | 51 (10.88 ± 0.33) | 3.91 | 48.60 | 0.97 | 0.41 |
|  | South Africa | 100 | 35 (2) | 42 (7.98 ± 1.37) | 3.01 | 10.37 | 0.48 | 0.23 |
|  | Uruguay | 11 | 28 (0) | 11 (11.00 ± 0.00) | 2.40 | 11.00 | 1.00 | 0.33 |
|  | Total | 938 | 96 | 358 (9.10 ± 1.29) | 4.50 | 18.82 | 0.20 | 0.28 |
| Australian sub-populations | New South Wales | 94 | 18 (0) | 21 (17.50 ± 1.38) | 2.72 | 12.27 | 0.79 | 0.08 |
|  | Queensland | 53 | 18 (1) | 12 (11.80 ± 0.41) | 1.85 | 4.21 | 0.60 | 0.06 |
|  | South Australia | 51 | 17 (0) | 8 (8.00 ± 0.00) | 1.19 | 2.12 | 0.49 | 0.06 |
|  | Tasmania | 69 | 19 (0) | 28 (24.00 ± 1.50) | 2.99 | 13.96 | 0.69 | 0.10 |
|  | Victoria | 133 | 21 (2) | 35 (20.00 ± 2.19) | 2.91 | 12.52 | 0.67 | 0.09 |
|  | New Zealand | 61 | 20 (1) | 33 (29.60 ± 1.36) | 3.30 | 22.28 | 0.82 | 0.12 |
|  | Total | 461 | 23 | 71 (22.30 ± 2.59) | 3.2 | 12.90 | 0.48 | 0.09 |

<sup>a</sup> N: Number of wasp specimens per collection site
<sup>b</sup> $A_R$ : Allelic richness observed per collection site
<sup>c</sup> $P_a$ : Number of private allelic richness observed per collection site
<sup>d</sup> MLG: Number of multi locus genotypes observed per collection site
<sup>e</sup> eMLG: Number of expected multilocus genotypes at the smallest sample size (11) based on rarefaction ± standard error)
<sup>f</sup> H: Shannon-Wiener Index of MLG diversity
<sup>g</sup> G: Stoddart and Taylor's Index of MLG diversity
<sup>h</sup> E5: Evenness (distribution of genotype abundance)
<sup>i</sup> $H_{exp}$ : Nei's unbiased gene diversity
\*: Native population

A similar trend in allelic richness was observed for the *A. areolatum* populations where A_R_ was higher in New Zealand (A_R_ = 24 and Pa= 2) than in Australia (A_R_ = 22, Pa = 0; Table 2). Overall, the American population was shown to have the highest allelic richness (A_R_ = 30 and Pa = 5; Table 2) with an average of 3.180 alleles per loci (Supplementary Table S7).

**Table 2.**
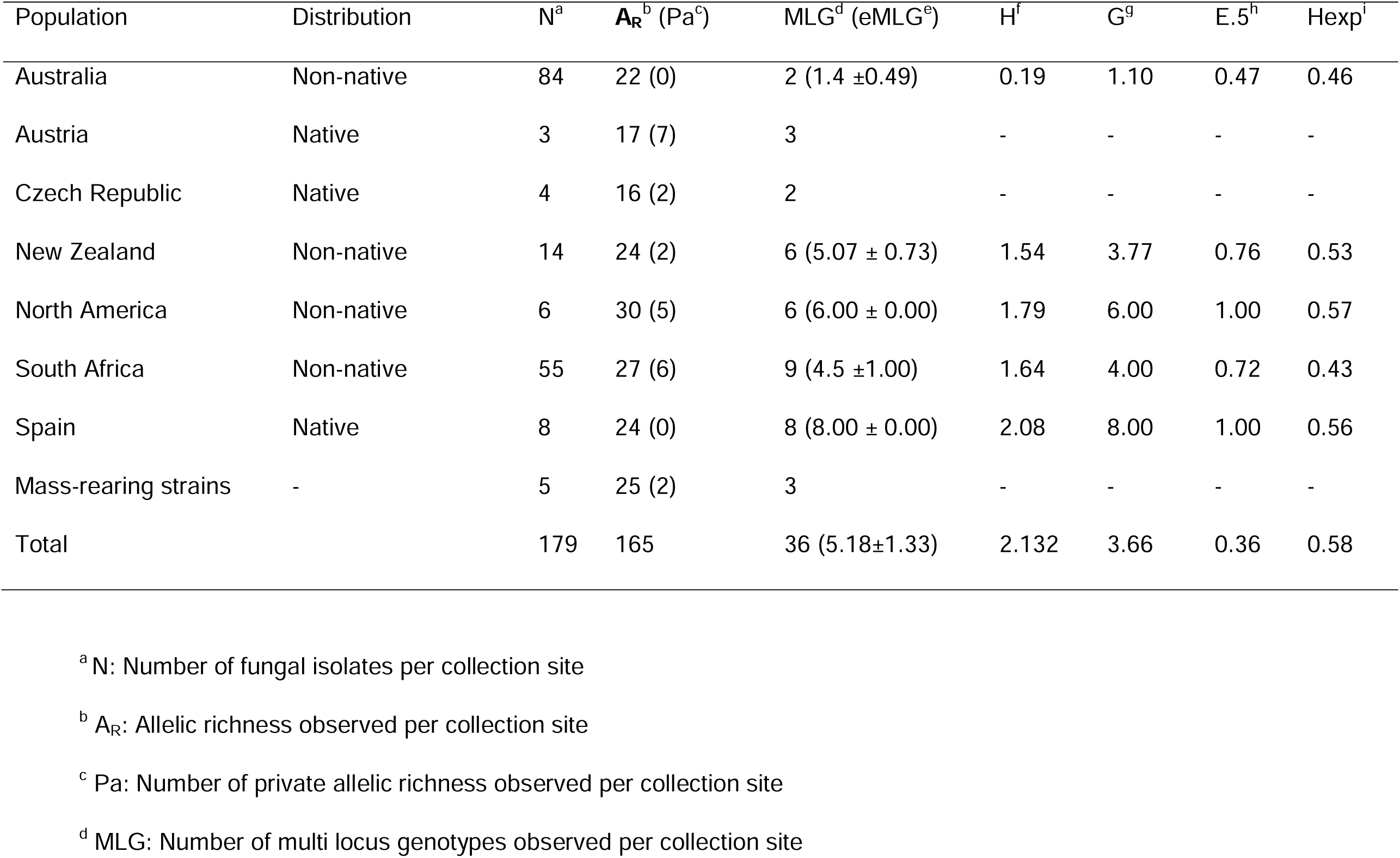
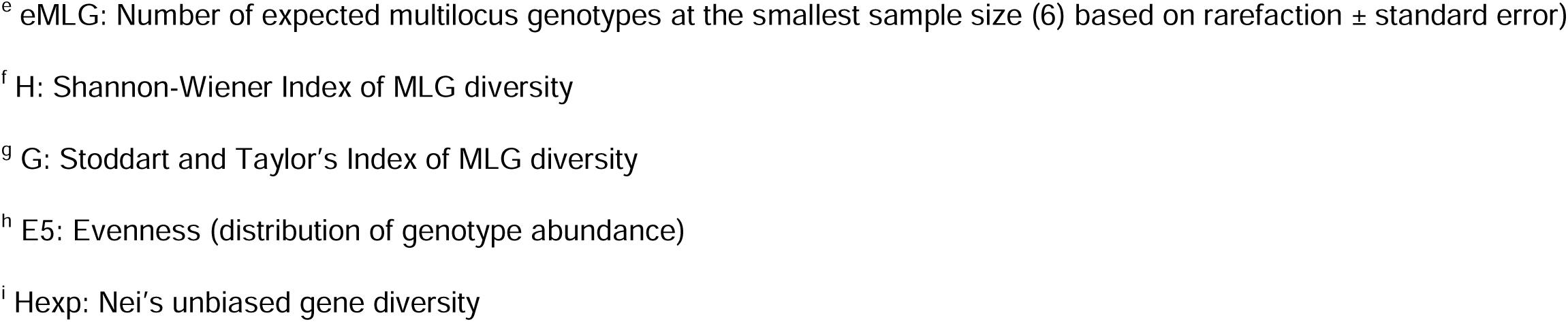
Summary of genetic diversity indices for *Amylostereum areolatum* from Mlonyeni et al. (2018), the global collection of isolates from the culture collection (CMW) and collections made in this study from Australia and New Zealand.

| Population | Distribution | N<sup>a</sup> | A<sub>R</sub><sup>b</sup> (Pa<sup>c</sup>) | MLG<sup>d</sup> (eMLG<sup>e</sup>) | H<sup>f</sup> | G<sup>g</sup> | E.5<sup>h</sup> | Hexp<sup>i</sup> |
| --- | --- | --- | --- | --- | --- | --- | --- | --- |
| Australia | Non-native | 84 | 22 (0) | 2 (1.4 ±0.49) | 0.19 | 1.10 | 0.47 | 0.46 |
| Austria | Native | 3 | 17 (7) | 3 | - | - | - | - |
| Czech Republic | Native | 4 | 16 (2) | 2 | - | - | - | - |
| New Zealand | Non-native | 14 | 24 (2) | 6 (5.07 ± 0.73) | 1.54 | 3.77 | 0.76 | 0.53 |
| North America | Non-native | 6 | 30 (5) | 6 (6.00 ± 0.00) | 1.79 | 6.00 | 1.00 | 0.57 |
| South Africa | Non-native | 55 | 27 (6) | 9 (4.5 ±1.00) | 1.64 | 4.00 | 0.72 | 0.43 |
| Spain | Native | 8 | 24 (0) | 8 (8.00 ± 0.00) | 2.08 | 8.00 | 1.00 | 0.56 |
| Mass-rearing strains | - | 5 | 25 (2) | 3 | - | - | - | - |
| Total |  | 179 | 165 | 36 (5.18±1.33) | 2.132 | 3.66 | 0.36 | 0.58 |

<sup>a</sup> N: Number of fungal isolates per collection site
<sup>b</sup> A<sub>R</sub>: Allelic richness observed per collection site
<sup>c</sup> Pa: Number of private allelic richness observed per collection site
<sup>d</sup> MLG: Number of multi locus genotypes observed per collection site
<sup>e</sup> eMLG: Number of expected multilocus genotypes at the smallest sample size (6) based on rarefaction ± standard error)
<sup>f</sup> H: Shannon-Wiener Index of MLG diversity
<sup>g</sup> G: Stoddart and Taylor's Index of MLG diversity
<sup>h</sup> E5: Evenness (distribution of genotype abundance)
<sup>i</sup> Hexp: Nei's unbiased gene diversity

A total of 71 MLGs were obtained from *S. noctilio* sub-populations in Australia and New Zealand (Table 1). The highest number of MLGs in Australia was observed from Victoria (n = 35) followed by Tasmania (n = 28) and New South Wales (n = 21). For the comparisons of gene diversity (Hexp) between the Australasian populations, with those from native and other introduced regions, the highest diversity was observed for European-Pop1 (Hexp = 0.45) followed by North America (Hexp = 0.41) (Table 1). The *S. noctilio* populations from Australia and New Zealand had the lowest gene diversity (Hexp = 0.07 and Hexp = 0.14, respectively). The population from Australia also had the lowest genotype diversity (H = 2.23, G = 4.96 and eMLG = 5.81; Table 1). The highest genotypic diversity was observed in introduced populations: Argentina (H = 4.39, G = 80.05, and eMLG = 10.98) followed by North America (H =3.91, G = 48.60, and eMLG = 10.88; Table 1). The native European population had the highest genotype abundance (E5 = 1.00), followed by Argentina from the invasive distribution (Table 1).

For *A. areolatum*, a total of 36 MLGs were obtained from all populations analyzed in this study (Table 2). Gene diversity (Hexp) values from the native and invasive populations ranged from 0.30 to 0.57 (Table 2). The highest gene diversity was observed for the population from North America followed by that from Spain and New Zealand, while the population from the Czech Republic had the lowest gene diversity (Table 2). Similarly, the highest genotypic diversity was observed for the native population from Spain, followed by diversity in the introduced populations from North America and South Africa (Table 2). Genotypic diversity and estimates of genotype abundance were moderate for populations from New Zealand and Australia compared to the high genotype abundances for Austria, North America, and Spain (Table 2).

#### Population molecular variance, genetic differentiation and gene flow

AMOVA analyses showed significant genetic differentiation (p ≤ 0.001) in the *S. noctilio* population for both the Australian sub-population and amongst populations in the native and introduced areas (Table 3). For the Australian sub-population, most of the genetic variation was due to variation within the population (54.98 %) compared to the low variation amongst the populations (8.51 %; Table 3). For the Australian and New Zealand *S. noctilio* populations compared with the global collection most of the genetic variation was observed amongst populations (55.40 %), compared to the lower variation within the population (17.30 %; Table 3). Likewise, for the *A. areolatum* populations most of the genetic variation was observed amongst populations (77.25 %) compared to variation within-population (22.75 %) variation (Table 4).

**Table 3.** Analysis of molecular variance (AMOVA) of the *Sirex noctilio* collection estimated between populations, within populations between samples, and within population.

|  | Source of variation | Degree of freedom | Sum of square | Mean square | Estimates of variance | Percentage (%) of the total variance | P-value |
| --- | --- | --- | --- | --- | --- | --- | --- |
| All populations | Between population | 8 | 2187.40 | 273.43 | 2.08 | 55.40 | 0.001 |
|  | Between samples within population | 774 | 1795.03 | 2.32 | 0.65 | 17.30 | 0.001 |
|  | Within samples | 783 | 801.00 | 1.02 | 1.02 | 27.30 | 0.001 |
|  | Total | 1565 | 4783.43 | 3.06 | 3.75 | 100.00 |  |
| Australian and New Zealand sub-populations | Between population | 5 | 90.20 | 18.041 | 0.11 | 8.51 | 0.001 |
|  | Between samples within population | 455 | 751.44 | 1.65 | 0.47 | 36.51 | 0.001 |
|  | Within samples | 461 | 327.00 | 0.71 | 0.71 | 54.98 | 0.001 |
|  | Total | 921 | 1168.64 | 1.27 | 1.29 | 100.00 |  |

**Table 4.** Analysis of molecular variance (AMOVA) of *Amylostereum areolatum* collections estimated between and within populations.

|  | Degree of<br>freedom | Sum of<br>square | Mean<br>square | Estimates of<br>variance | Percentage (%) of<br>the total variance | p-value |
| --- | --- | --- | --- | --- | --- | --- |
| Between population | 7 | 244.180 | 34.883 | 1.988 | 77.25 | 0.001 |
| Within populations | 171 | 100.125 | 0.586 | 0.586 | 22.75 | 0.001 |
| Total | 179 | 344.305 | 1.934 | 2.574 | 100.00 |  |

Analysis of pair-wise population differentiation based on genetic differentiation (DP_T_) and gene flow (Nm) in Australia showed that the highest genetic differentiation (DP_T_ = 0.326; Nm = 0. 517) occurred between *S. noctilio* populations in Queensland and South Australia (Supplementary Table S8). The lowest genetic differentiation (DP_T_ = 0.018; Nm = 13.388) was between the populations from New South Wales and Victoria (Supplementary Table S8). Comparing New Zealand with states in Australia, the lowest genetic differentiation (DP_T_ = 0.035) and highest gene flow (Nm = 6. 866) was revealed with the population in Tasmania followed by Victoria (DP_T_ = 0.056 vs Nm = 4.185; Supplementary Table S8). Genetic differentiation was lower between Australian and New Zealand populations (DP_T_ = 0.087 vs Nm = 2.626) as compared to other *S. noctilio* populations included from elsewhere in the world (Table 5). The highest differentiation and lowest gene flow were between Australian and Swiss populations (DP_T_ = 0.700 vs Nm = 0.107) followed by New Zealand and Swiss populations (DP_T_ = 0.486 vs Nm = 0.265; Table 5). The lowest differentiation and highest gene flow between native and invasive populations stand between populations from Europe and South America (Table 5).

**Table 5.** Summary of population differentiation between *Sirex noctilio* populations from Australia and New Zealand, in comparison with elsewhere in the world (Boissin et al. 2012). Genetic differentiation (lJP_T_) and gene flow (Nm) between pairs of *Sirex noctilio* populations are represented below and above the diagonal, respectively. Note F_ST_ (lJP_T_) 0 means complete sharing of genetic material and F_ST_ (lJP_T_) 1 means no sharing.

|  | Australia | New Zealand | Europe-Pop1 | Europe-Pop2 (Switzerland) | North America | South Africa | Argentina | Chile | Uruguay |
| --- | --- | --- | --- | --- | --- | --- | --- | --- | --- |
| Australia | - | 2.626 | 0.213 | 0.107 | 0.148 | 0.161 | 0.215 | 0.169 | 0.176 |
| New Zealand | 0.087 | - | 0.816 | 0.265 | 0.443 | 0.420 | 0.619 | 0.586 | 0.614 |
| Europe-Pop1 | 0.540 | 0.234 | - | 0.452 | 1.565 | 0.985 | 2.986 | 2.172 | 2.077 |
| Europe-Pop2 (Switzerland) | 0.700 | 0.486 | 0.356 | - | 0.328 | 0.376 | 0.305 | 1.435 | 0.308 |
| North America | 0.628 | 0.361 | 0.138 | 0.432 | - | 0.620 | 0.770 | 0.917 | 1.243 |
| South Africa | 0.608 | 0.373 | 0.202 | 0.400 | 0.287 | - | 0.445 | 1.057 | 0.591 |
| Argentina | 0.538 | 0.288 | 0.077 | 0.451 | 0.245 | 0.360 | - | 0.800 | 1.260 |
| Chile | 0.597 | 0.299 | 0.103 | 0.148 | 0.214 | 0.191 | 0.238 | - | 0.985 |
| Uruguay | 0.587 | 0.289 | 0.107 | 0.448 | 0.167 | 0.297 | 0.166 | 0.202 | - |

The analyses of pair-wise population genetic differentiation of the symbiotic fungus *A. areolatum* population also revealed the lowest genetic differentiation and highest gene flow between populations of Australia and New Zealand (Table 6). Comparing the Australian and New Zealand populations to that of the native population, the population from Spain showed the highest gene flow and lowest genetic differentiation (Table 6).

**Table 6.**
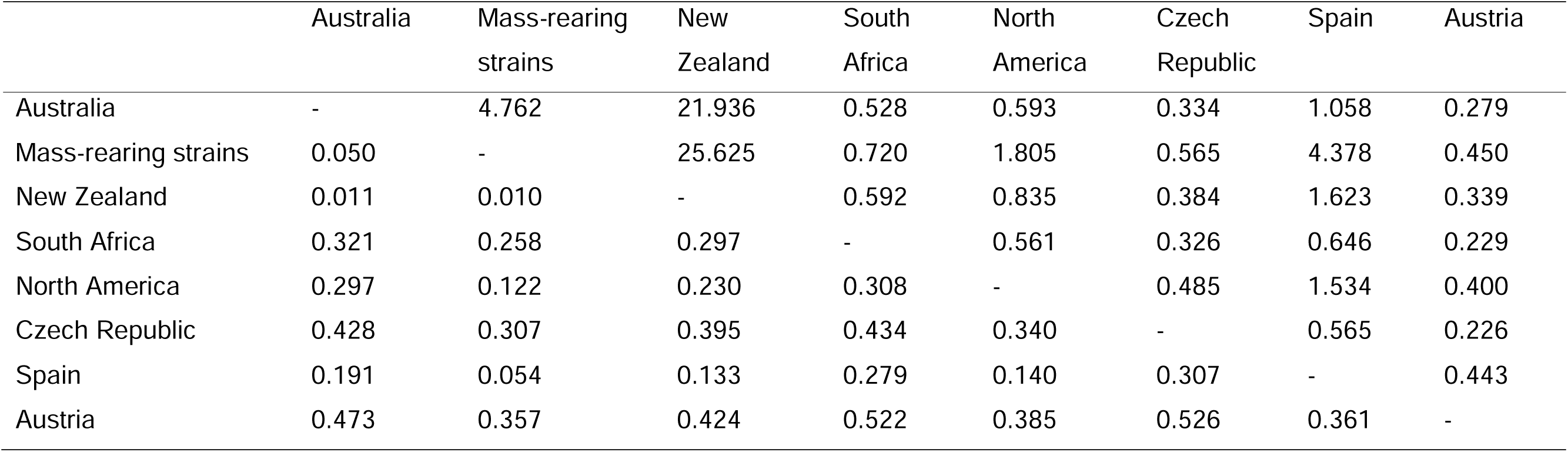
Summary of population differentiation between *Amylostereum areolatum* populations from Australia and New Zealand, in comparison with elsewhere in the world (Mlonyeni et al. 2018). Genetic differentiation (lJP_T_) and gene flow (Nm) between pairs of *Amylostereum areolatum* populations are represented below and above the diagonal, respectively. Note F_ST_ (lJP_T_) 0 means complete sharing of genetic material and F_ST_ (lJP_T_) 1 means no sharing.

|  | Australia | Mass-rearing strains | New Zealand | South Africa | North America | Czech Republic | Spain | Austria |
| --- | --- | --- | --- | --- | --- | --- | --- | --- |
| Australia | - | 4.762 | 21.936 | 0.528 | 0.593 | 0.334 | 1.058 | 0.279 |
| Mass-rearing strains | 0.050 | - | 25.625 | 0.720 | 1.805 | 0.565 | 4.378 | 0.450 |
| New Zealand | 0.011 | 0.010 | - | 0.592 | 0.835 | 0.384 | 1.623 | 0.339 |
| South Africa | 0.321 | 0.258 | 0.297 | - | 0.561 | 0.326 | 0.646 | 0.229 |
| North America | 0.297 | 0.122 | 0.230 | 0.308 | - | 0.485 | 1.534 | 0.400 |
| Czech Republic | 0.428 | 0.307 | 0.395 | 0.434 | 0.340 | - | 0.565 | 0.226 |
| Spain | 0.191 | 0.054 | 0.133 | 0.279 | 0.140 | 0.307 | - | 0.443 |
| Austria | 0.473 | 0.357 | 0.424 | 0.522 | 0.385 | 0.526 | 0.361 | - |

#### Analysis of Minimum Spanning Network

Minimum spanning network (MSN) analyses showed the unique diversity of *S. noctilio* in Australia and New Zealand (Fig. 4a, b). However, no clear genetic or geographic-based cluster was identified between *S. noctilio* populations in Australia and New Zealand (Fig. 4b). The historical samples of *S. noctilio* were important in showing the change in the genetic diversity over time (Fig. 4b). A single MLG from Tasmania collected in 1962 seemed to be the founder of many of the historical MLGs that are now extinct (Fig. 4b). There are also a few of these MLGs from historical samples that persist today with more diversity in the current population (Fig. 4b). MSN analyses revealed at least two distinctly identifiable clusters for the global *S. noctilio* populations, where most of the Australian and New Zealand individuals fall into one of the clusters (Fig. 4a).

**Fig. 4.**
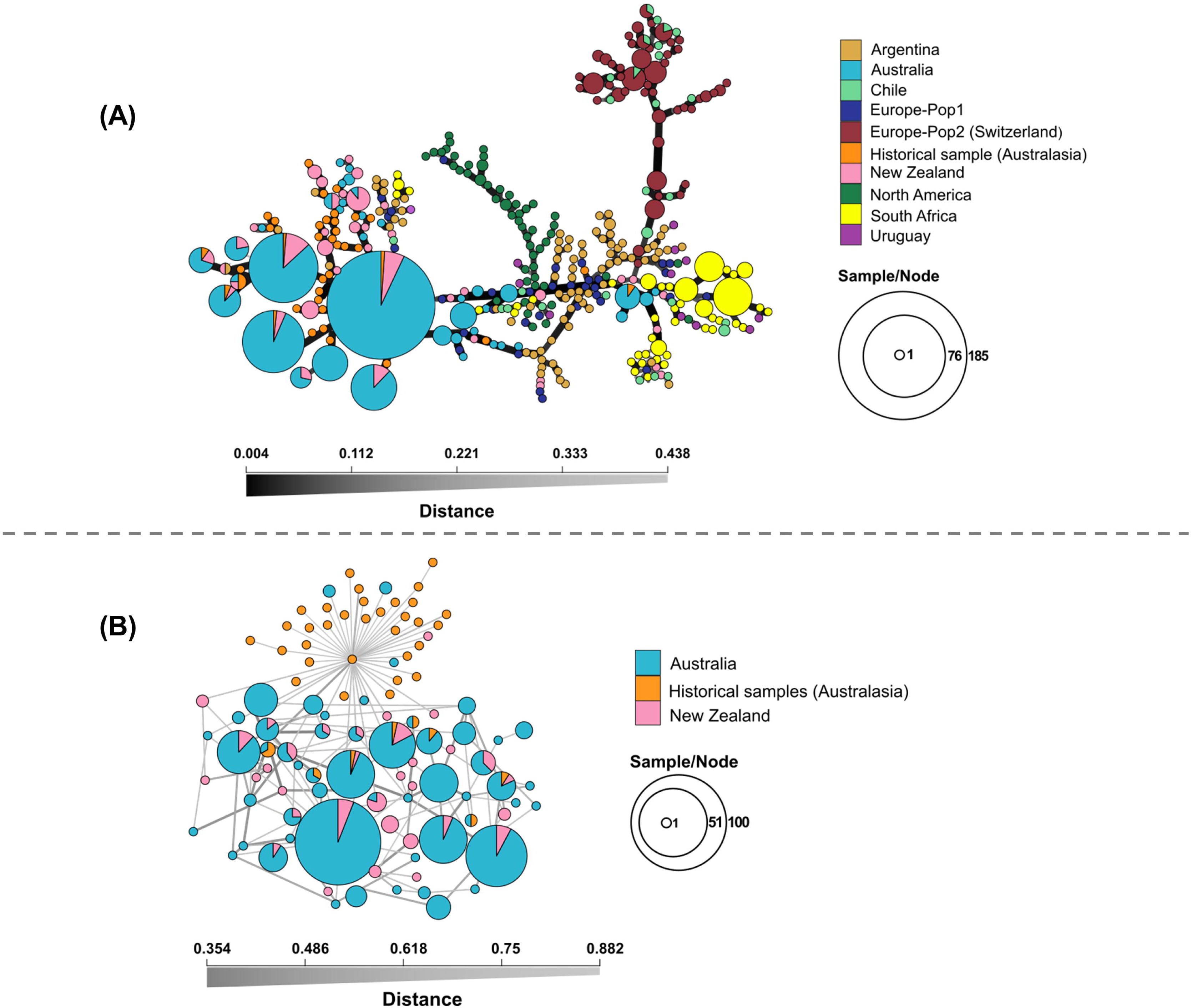
Minimum spanning network (MSN) of *S. noctilio* population constructed using Nei’s distances. (A) MSN analyses for 979 specimens showing three distinct clusters of the global population, and (B) of 502 specimens from populations in Australia and New Zealand. The sizes of the nodes are proportional to the number of specimens representing the MLG and the thickness of the lines represents the Nei genetic distance between two nodes (thicker lines denote smaller genetic distance). Colours represent different geographic origins.

MSN analyses showed three distinct global clusters of the *A. areolatum* populations and these clusters were separated by large genetic distance (Fig. 5). The clusters include populations from Australia, Mass-rearing and New Zealand (cluster I), South Africa (cluster II), and North America, Czech Republic, Spain and Austria (cluster III) (Fig. 5). MSN analyses did not show a defined cluster between the population in Australia and New Zealand and there was evidence of shared history at two MLGs (Fig. 5).

**Fig. 5.**
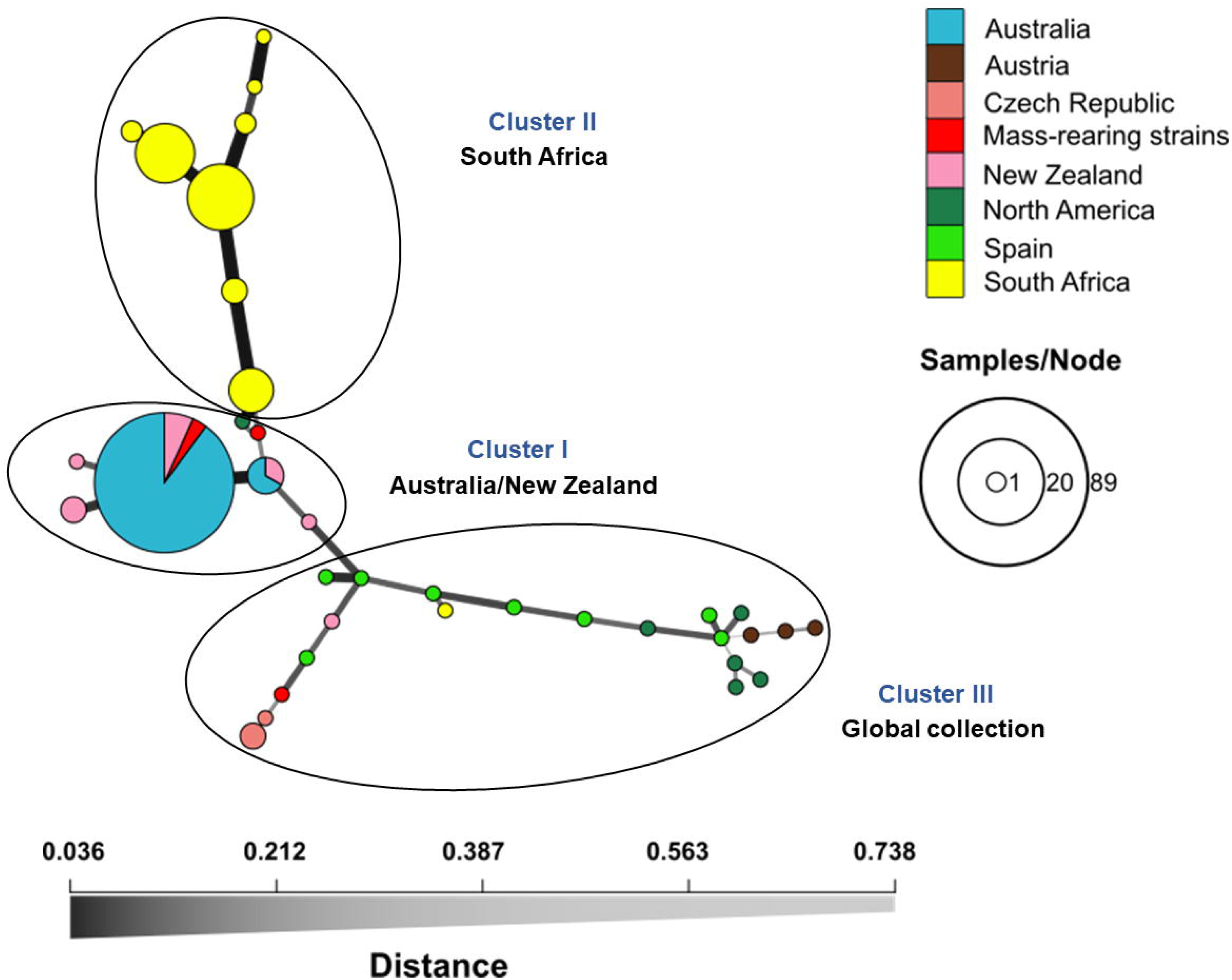
Minimum spanning network (MSN) of *A. areolatum* population constructed using Nei’s distances showing three distinct clusters for 179 isolates collected from Australia and New Zealand and CMW isolates for the global population. The sizes of the nodes are proportional to the number of specimens representing the MLG and the thickness of the lines represents the Nei genetic distance between two nodes (thicker) lines denote smaller genetic distance). Colours represent different geographic origins.

#### Population structure

Bayesian clustering-based population structure of *S. noctilio* was conducted using STRUCTURE analyses for 502 specimens from Australian and New Zealand populations and for 979 specimens that include the global collection (Fig. 6a, b). STRUCTURE analyses suggested three widespread global gene pools (Fig. 6a). This aligns with the distance-based population structure PCoA, which also revealed three global clusters (Supplementary Fig. S3). PCoA showed two coordinates describing 68.85 % of the total variation in the global *S. noctilio* collections (Supplementary Fig. S3). The ΔK statistics (Evanno et al. 2005) for the optimal K search, however, supported two populations globally (Supplementary Fig. S1a). For the Australian and New Zealand *S. noctilio* populations both the STRUCTURE and assessment of the ΔK statistics (Evanno et al. 2005) for the optimal K value search suggest the establishment of two populations (K= 2) (Supplementary Fig. S1b).

**Fig. 6.**
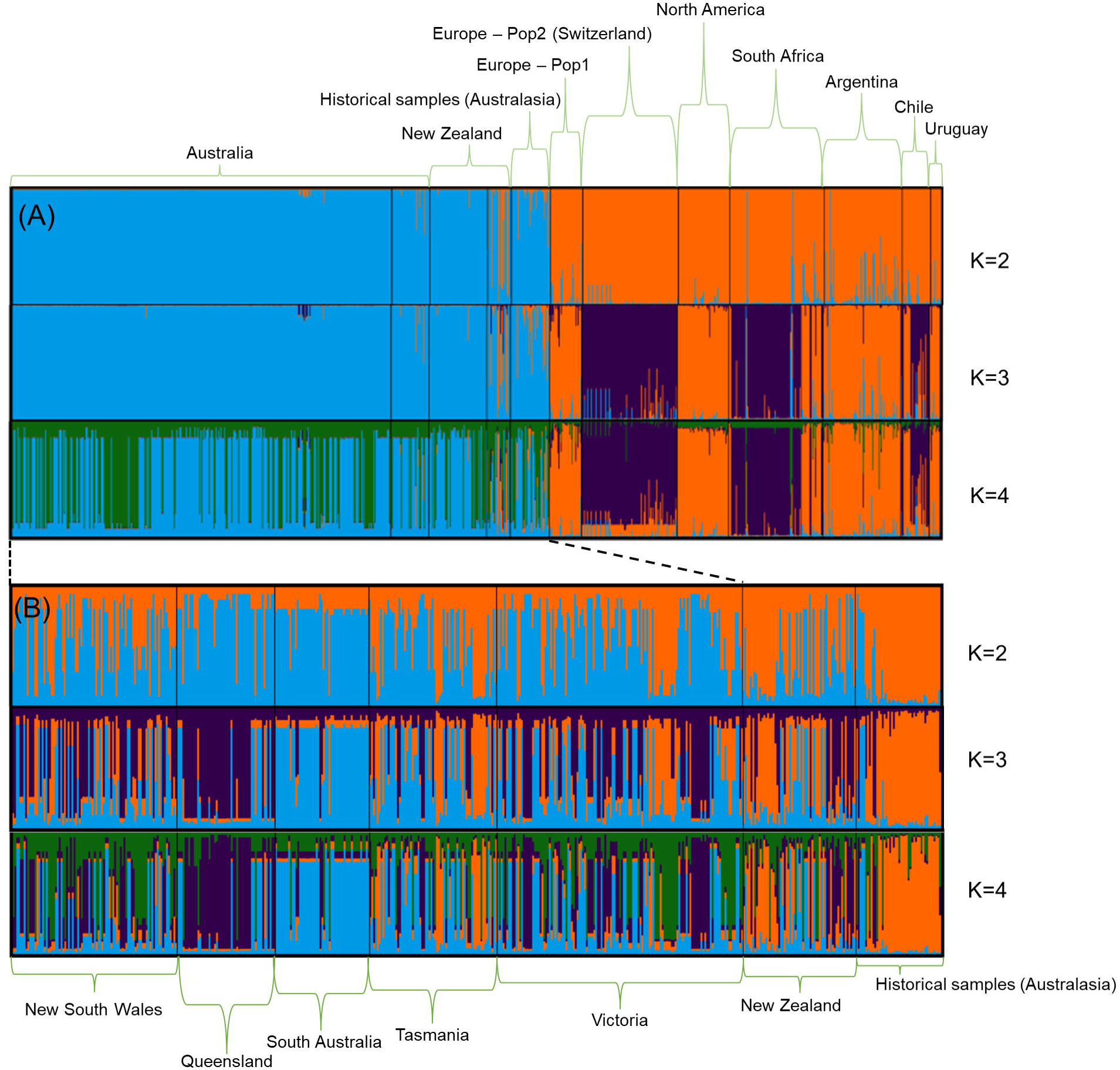
Bayesian clustering of *S. noctilio* populations using STRUCTURE. Clustering inferred (A) at Δk = 2, Δk = 3 and Δk = 4 using data from 979 specimens from across native and introduced regions and (B) at Δk = 1, Δk = 2 and Δk = 3 using data from the 502 *S. noctilio* specimens in Australian, New Zealand and historical samples obtained from these two countries. Different clusters are divided into K colours and the vertical bar represents the individual strain that lies within each cluster.

The STRUCTURE and assessment of the ΔK statistics (Evanno et al., 2005) for the best K search of 179 *A. areolatum* exhibited three global populations (Fig. 7; Supplementary Fig. S2). The PCoA analysis was congruent with STRUCTURE and the ΔK statistics, supporting the widespread of three global populations (Supplementary Fig. S2). The PCoA showed that two principal coordinates described 62.94 % of the total observed variation (Supplementary Fig. S4). Like that of the distance-based MSN analyses, PCoA revealed three population clusters.

**Fig. 7.**
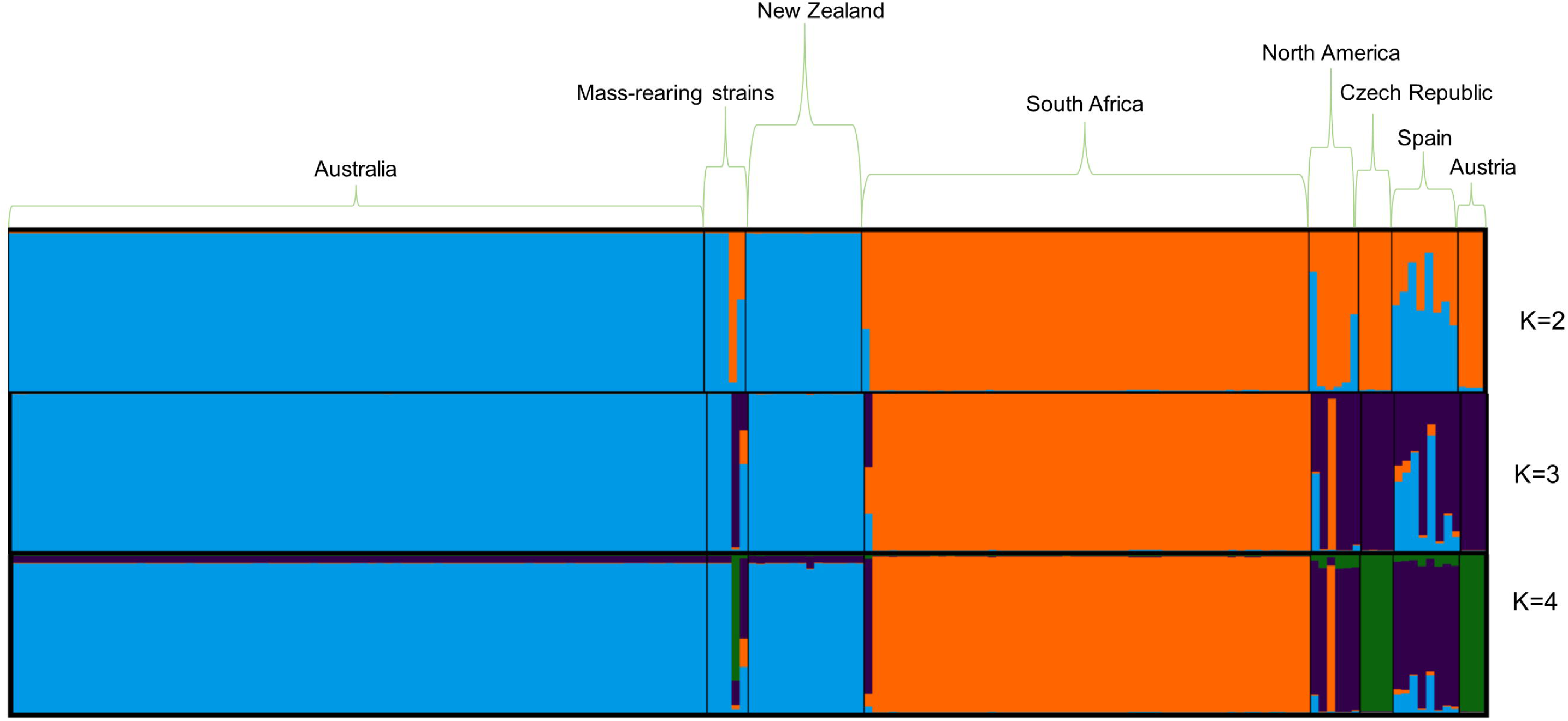
Bayesian clustering of *A. areolatum* populations using STRUCTURE. Clustering inferred at Δk = 2, Δk = 3 and Δk = 4 using data from 179 specimens from across native and introduced regions. Different clusters are divided into K colors and the vertical bar represents the individual strain that lies within each cluster.

## Discussion

This study provides unprecedented insight into the genetic diversity, invasion history and population structure of populations of *S. noctilio* and its symbiotic fungus *A. areolatum* in Australia and New Zealand. We found lower genetic diversity in *S. noctilio* and *A. areolatum* populations in these regions compared to other invaded regions, and a lack of population structure between Australia and New Zealand. The limited diversity in the woodwasp and fungus in Australasia likely reflects the impact of strict biosecurity measures that reduced the number of introduction events over time, compared with other parts of the world that experienced repeated introductions. Additionally, the clonality that persists in the fungal populations is an advantage for management programs using the parasitic nematode *D. siricidicola*, which feeds and is reared on the fungus.

The low diversity of *S. noctilio* in Australia has been noted previously (Boissin et al. 2012; Bittner et al. 2017) but for a limited number of samples from one location. *Sirex noctilio* has been in Australia for more than seven decades, so we expected that the genetic diversity could have accumulated from repeated introduction than in more recently invaded regions. However, our investigation negates this assumption. The low diversity of the woodwasp population in Australia must have been driven by combined impact of a genetic bottleneck and/or genetic drift, influenced by an effective quarantine system.

There was no population genetic structure among *S. noctilio* populations in Australia, as shown by all analyses, including allelic diversity and frequencies and the low number of private alleles amongst the populations from different states in Australia. Most of the diversity in Australia was within, rather than between, populations in different states, reflecting a high flow of genetic diversity across the areas. These results are not surprising considering that the spread of *S. noctilio* in Australia was estimated at 30 - 40 km/year which would alter natural spread across the distribution in a few decades with ample opportunity for reciprocal gene flow between (Carnegie et al. 2005). The pathways of spread of *S. noctilio* within Australia are predominantly linked to natural dispersal and partly to human-assisted transportation of infected timber and wood products (Carnegie et al. 2006). Abiotic and biotic factors such as temperature, host characteristics and density could directly and/or indirectly determine *S. noctilio* spread rates (Lantschner et al. 2014; Lantschner and Corley 2015). In mainland Australia, the commercial pine plantations are situated contagiously between different pine-growing states, and the range expansion occurred as predicted across these relatively contiguous plantation regions: Victoria going south toward South Australia, and northward to New South Wales and Queensland (Carnegie et al. 2006; Nahrung et al. 2015). However, the spread between Tasmania and mainland Australia more likely followed the human-assisted transportation of infected timber and wood materials. High gene flow between *S. noctilio* populations in Tasmania and mainland Australia, and between different states in mainland Australia aligns with the historic range expansion records of *S. noctilio* in Australia (Carnegie et al. 2005; Nahrung et al. 2015) and with the distribution of a naturally-dispersed lineage of *D. siricidicola* (Eshetu et al. 2023).

The genetic diversity of *S. noctilio* in New Zealand was higher than in the Australian woodwasp populations (45.35 % of the total sampled population (39 /86) represents a unique MLG in New Zealand vs only 8. 64 % which is 38 MLG of the total 440 samples collected from Australia). This diversity in New Zealand could either reflect a diverse gene pool of original incursion or could be due to consecutive multiple introductions before the implementation of proper biosecurity regulations. This is likely because *S. noctilio* management in New Zealand only received attention following the population outbreak on highly stressed pine trees almost 50 years after its first report (Miller and Clark 1935; Rawlings and Wilson 1949; Morgan and Stewart 1966).

The woodwasp populations had low genetic differentiation between Australia and New Zealand. The lack of population structure could be due to the founding populations originating from the same source or, more likely, that New Zealand served as the bridgehead population for introduction to Australia (Boissin et al. 2012; Slippers et al. 2015; Bertelsmeier and Keller 2018). There has been a high level of gene flow between New Zealand and Tasmania (oldest records of the woodwasp-fungus invasion in the region) followed by New Zealand and Victoria (the first record in mainland Australia) and this aligns with the records of historical introductions (Gilbert and Miller 1952; Collett and Elms 2009). The high level of gene flow between New Zealand and Australia was despite the geographic barrier between these two countries. The gene flow observed is most likely not the reflection of recent introductions, but it could rather reflect introductions that occurred before the implementation of strict phytosanitary measures in both countries. Likewise, the presence of Kamona strain nematodes in New Zealand (Fitza et al. 2019), despite never being deliberately released there, presumably resulted from the movement of infected female woodwasp/s prior to strengthened biosecurity.

We were in a unique position to compare current *S. noctilio* populations (2006-2018) to those from museum collections (1952-2001), illustrating the link between these populations. There was genetic overlap between the current and historical collections (26 % of the samples contained alleles that overlap with the current sample), but there was also a distinct shift evident in diversity. The oldest historical samples contained many unique alleles and MLGs that were not detected in the current populations. This shift is possible due to drift in these populations, and this most likely happened in periods soon after introduction when the populations would be expected to be small. In the decades since its introduction to Australia in the 1950s, the populations in this region have been quite large across the region and less likely to be susceptible to such drift. There is also unique diversity in the current populations that are not represented amongst the historical populations. This might be due to subsequent introductions, a mutation in the markers or under-sampling in the historical samples. For example, Victoria (where *S. noctilio* was reported first in mainland Australia in 1961 and was one of the outbreak sites in the GTR) was not represented in our historical sample collection. These results illustrate how diversity can shift over time in invasive populations. This unique collection of historical samples demonstrates the value of preserving specimens at the earliest stages of invasion.

The fungal populations from Australia and New Zealand group separately from all other populations elsewhere in the world that were included in this study. Two dominant genotypes are shared between Australia and New Zealand. While the connection and shared history between the fungal populations in Australia and Zealand likely aligned with the pattern of the woodwasp introductions, the persistence of only limited clones of the fungus could have arisen in several ways. First, like for the woodwasp, it is most likely the result of invasion from New Zealand to Australia (Boissin et al. 2012; Slippers et al. 2015). Secondly, multiple introductions of diverse fungal clones to these countries could have occurred, with selection favoring the persistence of only a few. Furthermore, it has been shown that the adaptation potential of *A. areolatum* clones could vary in different pine taxa (Wang et al. 2019), and *Amylostereum* is known to be a weak contender in the proximity of endophytic and coexisting sapstain fungi, vectored by bark beetles (Hurley et al. 2012; Yousuf et al. 2014; Wang et al. 2019). Such biotic factors are likely to compromise the chance of less adapted clones to persist between different wasp generations. Thirdly, low diversity is influenced by the transmission of asexually produced fungal arthrospores vertically between generations of the wasp (Thomsen and Koch 1999).

*Amylostereum areolatum* samples from Europe were distinct from Australia, New Zealand, and South Africa, with the exception of one shared mtSSU sequence haplotype being shared among fungal isolates from native (Spain and Austria) and non-native areas (Australia, New Zealand, South Africa and North America), supporting the hypothesis of Europe as the origin of global invasive woodwasp-fungal populations (Boissin et al. 2012). One of the oldest rearing isolates (CMW4644) used in mass-rearing of the biological control nematode, *D. siricidicola*, clustered with the European populations. Slippers et al. (2001) have also shown that the fungus initially used in biological control in Australia has a different IGS genotype than *A. areolatum* in the Southern Hemisphere. This strain was used in the mass-rearing program from 1995 and must have been introduced to Australia from Europe (Nahrung 2017). However, the current mass-rearing strain in Australia (Ecogrow: CMW53289) sourced from Queensland in 2015 (Nahrung 2017), shared the same MLG with the most field-dominant fungal clone (cluster I) in Australia. The second mass-rearing MLG (Aussie: Ecogrow, CMW40871) in this same cluster has been imported from Australia to South Africa for a rearing program (Mlonyeni et al. 2018). This is advantageous for the nematode rearing program in Australia because the interaction of different nematode-fungus genotypes has shown extreme variability of nematode reproduction (Hurley et al. 2012; Caetano et al. 2016; Morris et al. 2012, 2023), impacting the mass-rearing program and nematode performance against diverse *S. noctilio* populations in the field. Selecting a nematode from the diversity in Australia (Eshetu et al. 2023) that reproduces well on the dominant *A. areolatum* strain in Australia and introducing this into the rearing of the biological control agent presents an opportunity to potentially improve the program.

There is asymmetrical genetic diversity between the populations of the woodwasp *S. noctilio* and symbiotic fungus *A. areolatum* in Australasia, despite their shared introduction and establishment history into the region. This is in line with the previous studies that have investigated a similar pattern of misalignment in genetic diversity and population structure between the woodwasp-fungus population based on independent studies of each organism (Castrillo et al. 2015; Slippers et al. 2015; Bittner et al. 2017). However, the South African fungal population was relatively genetically diverse (Mlonyeni et al. 2018), and this aligns with the introduction history of the woodwasp population (Boissin et al. 2012). Australian and New Zealand *S. noctilio* populations have had the most frequently occurring genotypes that could be adapted to the existing biotic and abiotic factors. These genotypes therefore may become a risk in the absence of proper management, poorly managed and stressed trees, and if they escape to other pine-growing regions. The information in this study has the relevance of informing *S. noctilio* management in Australasia and beyond where the insect-fungus complex has become established and in predicted areas in the future. Moreover, results in this study should be useful for motivation for national and international phytosanitary measures.

## Acknowledgment

We are grateful to the Australian National Sirex Co-ordinate Committee (NSCC) and the Tree Protection Co-operative Programme (TPCP) at the Forestry and Agricultural Biotechnology Institute (FABI), University of Pretoria, South Africa for financial support, as well as for equipment and the use of facilities. We thank Stephen Elms, Matthew McCabe and Kate Brock (HVP Plantations), Craig Wilson (EcoGrow Australia), and NSCC partners Angus Carnegie (NSW), Nita Ramsden (Tasmania), and Kim Thomas and Gary Pearson (South Australia), for the sample collections. Geoff Allen kindly provided the historical samples from the University of Tasmania’s collection. We thank Scion, the New Zealand Forest Research Institute, for providing both woodwasp and fungus samples from New Zealand.

## Authors contributions

**FBE**: Conceptualization, Investigation, Data curation, Formal analysis and Writing-original draft. **IB**: Conceptualization and Data curation and Writing-reviewing and editing. **HNF**: Project administration, Funding acquisition, Resource, Conceptualization, Data curation and Writing-reviewing and editing. **KNEF**: Data curation and Writing-reviewing and editing. **BS**: Principal investigator, Project administration, Funding acquisition, Resource, Conceptualization, Data curation and Writing-reviewing and editing. All authors contributed to the writing of the manuscript and approved for publication.

## Data availability statement

A representative mitochondrial haplotype of the mtCOI sequence of the woodwasp *S. noctilio* and mitochondrial small subunit (mtSSU) ribosomal RNA gene sequence of the symbiotic fungus *A. areolatum* will be submitted to NCBI. All the SSR data have made available as Supplementary material data.

## Supplementary figure captions

Supplementary Fig. S1. Assessment of the best ΔK statistics Evanno et al (2005) for the best K search. (A) best a ΔK when comparing Australia and New Zealand *S. noctilio* with global population, and (B) best Δ K when comparing Australia and New Zealand *S. noctilio* population.

Supplementary Fig. S2. Assessment of the best ΔK statistics Evanno et al (2005) for the best K search when comparing Australia and New Zealand *A. areolatum* with the global population.

Supplementary Fig. S3. Principal coordinate analysis plot (PCoA) of 979 *S. noctilio* specimens from across native and introduced regions around the world, constructed using Nei’s genetic distance (Nei 1978). The percentage at coordinates 1 and 2 indicates the extent of observed genetic variation in the simulated dataset. Colours represent the geographic origin of strains. AGT, Argentina; AUS, Australia; CHL, Chile; EUR, Europe-Pop1; CHE, Europe-Pop2 (Switzerland); HS, Historical samples (Australasia); NZL, New Zealand; NA, North America; RSA, South Africa; and URY, Uruguay.

Supplementary Fig. S4. Principal coordinate analysis plot (PCoA) of 179 *A. areolatum* isolates, constructed using Nei’s genetic distance (Nei 1978). The percentage at coordinates 1 and 2 indicates the extent of observed genetic variation in the simulated dataset. Colours represent the geographic origin of strains. AUS, Australia; MRC, Mass-rearing strains; NZL, New Zealand; RSA, South Africa; NorthA, North America; CZ, Czech Republic; ESP, Spain; and AT, Austria

