## Supplementary Table S1 for "The influence of biosecurity on the diversity and management of a 120-year old joint insect-fungus pest invasion"

Supplementary Table S1. *Sirex noctilio* collections from Australia and New Zealand are characterized in this study. Numbers in brackets under the sampling year column indicate the number of samples collected in each year and no brackets mean samples were collected in the same year.

| State | District | Block name | Sampling year | No. of samples |
| --- | --- | --- | --- | --- |
| Australia |  |  |  |  |
| New South Wales | NSW | Goldsmiths Block | 2015 | 5 |
|  |  | Nowendoc SF | 2015 | 4 |
|  |  | Bago SF | 2016 | 3 |
|  |  | Bucclcuch SF | 2016 | 3 |
|  |  | Canobolos SF | 2016 | 4 |
|  |  | Glenwood SF | 2015 | 3 |
|  |  | Mannus SF | 2015 | 4 |
|  |  | Wingello SF | 2016 | 1 |
|  |  | Nangus JV | 2016 | 1 |
|  |  | Mt. Lambie Block | 2016 | 2 |
|  |  | Green Hills SF | 1986 (1), 2016 (10) | 11 |
|  |  | DD, Maragle SF | 2016 | 1 |
|  |  | Gurnang SF | 2016 | 1 |
|  |  | Coolangubra SF | 2016 | 2 |
|  |  | Kinross SF | 2015 | 4 |
|  |  | Vulcan SF | 2016 | 1 |
|  |  | Sunny Corner | 2016 (1), 2018 (7) | 8 |
|  |  | Blowering | 2016 | 7 |
|  |  | Dog Rocks | 2016 (3), unknown (4) | 7 |
|  |  | Skinners Road | 2016 | 1 |
|  |  | Nalbaugh | 2016 | 1 |
|  |  | Hampton | unknown | 1 |
|  |  | Tumut | 2000 (3), 2011(16) | 19 |
|  |  | Hume | 2006 | 2 |
|  |  | Penrose State Forest | 1989 | 1 |
|  |  | Sydney | 1987 | 2 |
|  |  | Mount Macquarie SF | 1993 | 1 |
| Queensland |  | Passchendaele | 2009 (1), 2011 (40), 2012 (10), 2014 (2) | 53 |
| South Australia |  | Noolook-Saltwell | 2017 | 12 |
|  |  | Comaum-Taylors | 2017 | 16 |
|  |  | Comaum | unknown | 4 |
|  |  | Cave Range | 2017 | 2 |
|  |  | Penola-Border | unknown | 2 |
|  |  | Comaum-Penders | unknown | 1 |
|  |  | Sandwood | unknown | 2 |
|  |  | Mt Burr | unknown | 3 |
|  |  | Myora | unknown | 1 |
|  |  | Mt. Gambier | 1988 | 2 |
|  |  | Unkown | 2014 | 8 |
| Tasmania | Murchison-NW | Oonah | 2014 (2), 2015 (2) | 4 |
|  | Bass-NE | Long Hill | 2014 | 5 |
|  | Bass-Central North | Badger Hills | 2013 (5), 2014 (5), 2015 (12), 2017 (2) | 24 |
|  | Bass-NE | Branches Creek | 2016 (2) | 2 |
|  | Bass-NE | Saddleback | 2014 (4), 2016 (3), 2017 (2), unknown (1) | 10 |
|  | Tas | SA Panel 846 | 2017 | 1 |
|  | Tas | SA Panel 848 | 2017 | 1 |
|  | Bass-NE | Lisle | 2014 (2), 2015 (3), 2016 (1), 2017 (7) | 13 |
|  | Bass-NE | Tower Hill | 2014 (2), 2015 (2), 2016 (4) | 8 |
|  | Bass-NE | Evercreech | 2017 | 1 |
|  | Tas | University of Tasmania | 1973 | 2 |
|  | Tas | Selbourne Tasmania | 2007 | 1 |
|  | Tas | Hobart | 1973 | 2 |
|  | Tas | Lucaston | 1999 | 1 |
|  | Tas | Austins Ferry | 2001 | 1 |
|  | Tas | Cambridge | 1963 (3), 1965 (1) | 4 |
|  | Tas | Ex insectary (incubated) | 1962 (2), 1963 (1) | 3 |
|  | Tas | Unkown | 1952 (2), 1953 (1) | 3 |
| Victoria | Shelley | Koetong | 2014 (2), 2015 (4), 2016 (12), 2017 (4), 2018 (7) | 29 |
|  | Shelley | Jinjellic | 2015 (1), 2016 (1), 2017 (2) | 5 |
|  | Benalla | Warrenbayne | 2014 (2), 2015 (4), 2016 (6), 2017 (2) | 14 |
|  | Myrtleford | Porepunkah | 2014 (1), 2015 (2) | 3 |
|  | Shelley | Railway (Shelley) | 2014 (1), 2015 (1), 2016 (1) | 3 |
|  | Myrtleford | Magpie | 2015 (1), 2016 (3) | 4 |
|  | Myrtleford | Stanley | 2016 (2), 2018 (2) | 4 |
|  | Myrtleford | Merriang | 2015 (5), 2016 (6), 2017 (2), 2018 (10) | 23 |
|  | Myrtleford | Bright | 2015 (1), 2016 (1) | 2 |
|  | Benalla | Holland | 2017 (2) | 2 |
|  | Myrtleford | Ovens | 2017 | 1 |
|  | Rennick | Kentbruck | 2017 | 1 |
|  | Benalla | Blue Range | 2015 (1), 2017 (1), 2018 (3) | 5 |
|  | Gippsland | Bradvale | 2016 | 1 |
|  | Gippsland | Livingston | 2016 | 1 |
|  | Myrtleford | German Ck | 2015 | 1 |
|  | Gippsland | Albert | 2017 | 1 |
|  | SW Ballarat | Spargo | 2016 | 1 |
|  | Gippsland | Longford | 2016 (2), 2018 (14) | 16 |
|  | Gippsland | Mack | 2016 | 1 |
|  | Gippsland | Boola | 2018 | 1 |
|  | Benalla | Upper Ryan | 2018 (8) | 8 |
|  | SW Rennick | Mt Richmond | unknown | 1 |
|  | SW Rennick | Rennick | 2018 | 1 |
|  | SW Rennick | Mocamboro | 2018 | 1 |
|  | Myrtleford | Havilah | 2018 | 1 |
|  | Unkown | Unkown | unknown | 2 |
| New Zealand |  |  |  |  |
| North Island |  | Kaingaroa Forest | 2017 | 15 |
|  |  | Mohaka Forest | 2017 | 22 |
|  |  | Santoft Forest | 2017 | 8 |
|  |  | Tohorakuri block, Taupo | 1959 | 1 |
|  |  | Golden Downs S. F | 1974 | 3 |
|  |  | Matangi | 1986 | 1 |
| Southern Island |  | Eyrewell | 2015 | 1 |
|  |  | Bottle Lake | 2015 | 5 |
|  |  | Amberley | 2015 | 10 |
|  |  | Herbert S. F | 1974 (2), 1975 (1) | 3 |
|  |  | Unkown | 1957 (3), 1961 (2), 1964 (2), 1966 (1) | 8 |
| Total |  |  |  | 502 |
