## Supplementary Table S2 for "The influence of biosecurity on the diversity and management of a 120-year old joint insect-fungus pest invasion"

Supplementary Table S2. *Amylostereum areolatum* collections from Australia and New Zealand are characterized in this study. Numbers in brackets under the sampling year column indicate the samples collected yearly.

| State | District | Block name | Sampling year | No. of samples |
| --- | --- | --- | --- | --- |
| Australia |  |  |  |  |
| New South Wales | NSW | Dog Rocks | 2016 (2), unknown (2) | 4 |
|  |  | Skinners Road | 2016 | 2 |
|  |  | Green Hills | 2016 (1), unknown (4) | 5 |
|  |  | Sunny corner | Unknown | 6 |
|  |  | Dark corner | Unknown | 1 |
|  |  | Mt. Lambie Block | 2017 | 1 |
|  |  | Blowering | Unknown | 2 |
|  |  | Bago | Unknown | 1 |
|  |  | Buccleuh | Unknown | 1 |
|  |  | Kinross | 2016 | 1 |
|  |  | Hampton | Unknown | 1 |
| Queensland |  | Passchendaele | Unknown | 10 |
| South Australia |  | Noolook-Saltwell | 2017 | 1 |
|  |  | Comaum | Unknown | 2 |
|  |  | Sandwood | Unknown | 3 |
|  |  | Mt. Burr | Unknown | 3 |
|  |  | Myora | Unknown | 1 |
| Victoria | Shelley | Railway (Shelley) | 2014, 2015 | 2 |
|  | Myrtleford | Bruarong | 2014 | 1 |
|  | Benalla | Warrenbayne | 2014 (4), 2015 (2) | 6 |
|  | Myrtleford | Stanley | 2014, 2017 | 2 |
|  | Benalla | Stanley | 2018 | 2 |
|  | Ovens | Stanley | 2014 | 1 |
|  | Gippsland | Jack | 2014 | 1 |
|  | Myrtleford | Merriang | 2015 (3), 2017 (2), 2018 (9) | 14 |
|  | Ballarat | Mt. Lonarch | 2017 | 2 |
|  | Benalla | Blue Range | 2015 (1), 2018 (3) | 4 |
|  | Gippsland | Bradvale | 2016 | 1 |
|  | Myrtleford | Porepunkah | 2015 | 1 |
|  | Myrtleford | Magpie | 2016 | 1 |
|  | Shelley | Koetong | 2015 (2), 2016 (1), 2017 (3), 2018 (7) | 13 |
|  | SW Ballarat | Spargo | 2016 | 1 |
|  | Gippsland | Longford | 2018 | 8 |
|  | Gippsland | Boola | 2018 | 1 |
|  | Benalla | Upper Ryan | 2018 | 8 |
|  | SW Rennick | Mt. Richmond | 2018 | 1 |
|  | SW Rennick | Rennick | 2018 | 1 |
|  | SW Rennick | Mocamboro | 2018 | 1 |
| New Zealand | New Zealand Region | Forest |  |  |
|  | Bay of Plentry | Kaingaroa | 1962 (2), 1967 (1), 2017 (10) | 13 |
|  | Hawke’s Bay | Mohaka | 2017 | 4 |
|  | ManawantuWanganui | Santoft | 2017 | 7 |
|  | Rotorua | Whakarewarewa Forest | 1962 | 1 |
|  | Okere Falls | Rotoehu Forest | 1962 | 1 |
| Commercial strain | Collection | Status |  |  |
| EcoGrowSubbed2013 | *Ecogrow | Sourced from Queensland |  | 1 |
| EcogrowRM1 | *Ecogrow | Sourced from Queensland |  | 1 |
| Kamona Fungus | *Ecogrow | - |  | 1 |
| Total |  |  |  | 146 |

*Ecogrow: refers to *Amylostereum areolatum* strain used for commercial nematode mass-production in Australia.
