## Supplementary Table S3 for "The influence of biosecurity on the diversity and management of a 120-year old joint insect-fungus pest invasion"

Supplementary Table S3. Culture collection (CMW) isolates of *A. areolatum* characterized in this study and used as a reference for the global collection.

| *CMW number | Continent | Country | Region | Collection year | Host/Vector | *Gene regions |
| --- | --- | --- | --- | --- | --- | --- |
| CMW4644 | Oceania | Australia | Unknown | 1995 | Isolates from nematode cultures from CSIRO | Mt (SSU rRNA); Nuc (SSR) |
| CMW6863 | Oceania | Australia | Unknown | 2001 | *Sirex noctilio* | Mt (SSU rRNA) |
| CMW37117 | Oceania | New Zealand | Kaingaroa Forest (Rotorua) | 2011 | *Sirex noctilio* | Mt (SSU rRNA); Nuc (SSR) |
| CMW8902 | Asia | Russia | Unknown | 1994 | *Picea abies* | Mt (SSU rRNA) |
| CMW8905 | Asia | Russia | Unknown | 1994 | *Picea abies* | Mt (SSU rRNA) |
| CMW13659 | Africa | South Africa | KwaZulu-Natal | 2003 | *Sirex noctilio* | Mt (SSU rRNA); Nuc (SSR) |
| CMW15098 | Europe | Denmark | Randbol | 1993 | Picea abies | Mt (SSU rRNA) |
| CMW15112 | Europe | Denmark | Rold Skov | 1983 | *Picea abies* | Mt (SSU rRNA) |
| CMW16841 | Europe | Austria | Knittenfeld | 2004 | Unknown | Mt (SSU rRNA) |
| CMW16850 | Europe | Austria | Knittenfeld | 2004 | Unknown | Mt (SSU rRNA); Nuc (SSR) |
| CMW16851 | Europe | Austria | Knittenfeld | 2004 | Unknown | Mt (SSU rRNA) |
| CMW16854 | Europe | Austria | Knittenfeld | 2004 | Unknown | Mt (SSU rRNA); Nuc (SSR) |
| CMW16859 | Europe | Austria | Knittenfeld | 2004 | Unknown | Mt (SSU rRNA); Nuc (SSR) |
| CMW27368 | Europe | Czech Republic | South Moravia | 2007 | *Sirex juvencus* | Mt (SSU rRNA); Nuc (SSR) |
| CMW27369 | Europe | Czech Republic | South Moravia | 2007 | *Sirex juvencus* | Nuc (SSR) |
| CMW27371 | Europe | Czech Republic | South Moravia | 2007 | *Sirex juvencus* | Mt (SSU rRNA); Nuc (SSR) |
| CMW27374 | Europe | Czech Republic | South Moravia | 2007 | *Sirex juvencus* | Mt (SSU rRNA); Nuc (SSR) |
| CMW40566 | Europe | Spain | Lugo (Galicia) | 2013 | *Sirex noctilio* | Mt (SSU rRNA); Nuc (SSR) |
| CMW40567 | Europe | Spain | Lugo (Galicia) | 2013 | *Sirex noctilio* | Mt (SSU rRNA); Nuc (SSR) |
| CMW40568 | Europe | Spain | Lugo (Galicia) | 2013 | *Sirex noctilio* | Mt (SSU rRNA); Nuc (SSR) |
| CMW40570 | Europe | Spain | Lugo (Galicia) | 2013 | *Sirex noctilio* | Mt (SSU rRNA); Nuc (SSR) |
| CMW42632 | Europe | Spain | Galicia (Begonte) | 2014 | *Sirex noctilio* | Nuc (SSR) |
| CMW46039 | Europe | Spain | Lugo (Galicia) | 2015 | *Sirex noctilio* | Nuc (SSR) |
| CMW42630 | Europe | Spain | Galicia (Cova Da Serpe) | 2014 | *Sirex noctilio* | Mt (SSU rRNA); Nuc (SSR) |
| CMW46037 | Europe | Spain | Lugo (Galicia) | 2015 | *Sirex noctilio* | Nuc (11SSR) |
| CMW42617 | North America | USA | Pennsylvania | 2014 | *Sirex noctilio* | Nuc (11SSR) |
| CMW42624 | North America | USA | Vermont | 2014 | *Sirex noctilio* | Mt (SSU rRNA); Nuc (SSR) |
| CMW42626 | North America | USA | Vermont | 2014 | *Sirex noctilio* | Mt (SSU rRNA); Nuc (SSR) |
| CMW37118 | North America | Canada | Unknown | Unknown | Unknown | Nuc (SSR) |
| CMW37119 | North America | Canada | Ontario | 2006 | *Sirex noctilio* | Nuc (SSR) |

*CMW= culture collection of the Forestry and Agricultural Biotechnology Institute (FABI), University of Pretoria

*****Gene regions= refers to the molecular markers used to analyze these CMW *A. areolatum* isolates
