## Supplementary Table S4 for "The influence of biosecurity on the diversity and management of a 120-year old joint insect-fungus pest invasion"

Supplementary Table S4. Alleles scored at 14 loci for strains of *S. noctilio*. ‘-1’ indicates alleles that could not be scored and microsatellite markers in this study were designed by (Santana et al. 2009). The forward primers of these loci were fluorescently labeled using FAM, NED, PET and VIC, dyes for filter set G5 (Applied biosystem). The fragments were analyzed on an ABI PRISM TM 3500xI analyser (Applied Biosystems) at the DNA sequencing facility, University of Pretoria. This collection included samples from Australia and New Newland (Samples IDs: **New South Wales**, N1 -NO4I; **Queensland**, Q2 – Q19B; **South Australia**, S1 – S8J; **Tasmania**, T1 – T79; **Victoria**, V1 – V497; **New Zealand**, NZS1 – Z16; **Historical samples Australia**, NO2B – TH9 and **Historical samples New Zealand**, NZ1H1 – NZLH16).

| Sample ID | Sn90 | | Sn104 | | Sn185 | | Sn177 | | SnB2 | | S97 | | Sn641 | | Sn525 | | SnB4 | | S92 | | Sn576 | | S93 | | Sn507 | | Sn231 | |
| --- | --- | --- | --- | --- | --- | --- | --- | --- | --- | --- | --- | --- | --- | --- | --- | --- | --- | --- | --- | --- | --- | --- | --- | --- | --- | --- | --- | --- |
| N1 | 101 | 101 | 177 | 177 | 203 | 203 | 157 | 157 | 187 | 187 | 196 | 196 | 229 | 229 | 188 | 188 | 173 | 173 | 194 | 194 | 212 | 212 | 207 | 207 | 223 | 223 | 188 | 188 |
| N2 | 101 | 101 | 177 | 177 | 203 | 203 | 157 | 157 | 187 | 187 | 196 | 196 | 229 | 229 | 188 | 188 | 173 | 173 | 194 | 194 | 212 | 212 | 207 | 207 | 223 | 223 | 188 | 207 |
| N3 | 101 | 101 | 177 | 177 | 203 | 203 | 157 | 157 | 187 | 187 | 196 | 196 | 229 | 229 | 188 | 188 | 173 | 173 | 194 | 194 | 212 | 212 | 195 | 219 | 223 | 223 | 207 | 207 |
| N4 | 101 | 101 | 177 | 177 | 203 | 203 | 157 | 157 | 187 | 187 | 196 | 196 | 229 | 229 | 188 | 188 | 173 | 173 | 194 | 194 | 212 | 212 | 195 | 219 | 223 | 223 | 207 | 207 |
| N8 | 101 | 101 | 177 | 177 | 203 | 203 | 157 | 157 | 187 | 187 | 196 | 196 | 229 | 229 | 188 | 188 | 173 | 173 | 194 | 194 | 212 | 212 | 195 | 207 | 223 | 223 | 207 | 207 |
| N12 | 101 | 101 | 177 | 177 | 203 | 203 | 157 | 157 | 187 | 187 | 196 | 196 | 229 | 229 | 188 | 188 | 173 | 173 | 194 | 194 | 212 | 212 | 195 | 207 | 223 | 223 | 207 | 207 |
| N16 | 101 | 101 | 177 | 177 | 203 | 203 | 157 | 157 | 187 | 187 | 196 | 196 | 229 | 229 | 188 | 188 | 173 | 173 | 194 | 194 | 212 | 212 | 219 | 219 | 223 | 223 | 207 | 207 |
| N19 | 101 | 101 | 177 | 177 | 203 | 203 | 157 | 157 | 187 | 187 | 196 | 196 | 229 | 229 | 188 | 188 | 173 | 173 | 194 | 194 | 212 | 212 | 207 | 207 | 223 | 223 | 207 | 207 |
| N23 | 101 | 101 | 177 | 177 | 203 | 203 | 157 | 157 | 187 | 187 | 196 | 196 | 229 | 229 | 188 | 188 | 173 | 173 | 194 | 194 | 212 | 212 | 195 | 195 | 223 | 223 | 188 | 188 |
| N24 | 101 | 101 | 177 | 177 | 203 | 203 | 157 | 157 | 187 | 187 | 196 | 196 | 229 | 229 | 188 | 188 | 173 | 173 | 194 | 194 | 212 | 212 | 195 | 207 | 223 | 223 | 207 | 207 |
| N28 | 101 | 101 | 177 | 177 | 203 | 203 | 157 | 157 | 187 | 187 | 196 | 196 | 229 | 229 | 188 | 188 | 173 | 173 | 194 | 194 | 212 | 212 | 195 | 207 | 223 | 223 | 207 | 207 |
| N49 | 101 | 101 | 177 | 177 | 203 | 203 | 157 | 157 | 187 | 187 | 196 | 196 | 229 | 229 | 188 | 188 | 173 | 173 | 194 | 194 | 212 | 212 | 195 | 207 | 223 | 223 | 207 | 207 |
| N50 | 101 | 101 | 177 | 177 | 203 | 203 | 157 | 157 | 187 | 187 | 196 | 196 | 229 | 229 | 188 | 188 | 173 | 173 | 194 | 194 | 212 | 212 | 195 | 195 | 223 | 223 | 188 | 188 |
| N51 | 101 | 101 | 177 | 177 | 203 | 203 | 157 | 157 | 187 | 187 | 196 | 196 | 229 | 229 | 188 | 188 | 173 | 173 | 194 | 194 | 212 | 212 | 219 | 219 | 223 | 223 | 188 | 188 |
| N52 | 101 | 101 | 177 | 177 | 203 | 203 | 157 | 157 | 187 | 187 | 196 | 196 | 229 | 229 | 188 | 188 | 173 | 173 | 194 | 194 | 212 | 212 | 195 | 195 | 223 | 223 | 207 | 207 |
| N53 | 101 | 101 | 177 | 177 | 203 | 203 | 157 | 157 | 187 | 187 | 196 | 196 | 229 | 229 | 188 | 188 | 173 | 173 | 194 | 194 | 212 | 212 | 195 | 207 | 223 | 223 | 207 | 207 |
| N54 | 101 | 101 | 177 | 177 | 203 | 203 | 157 | 157 | 187 | 187 | 196 | 196 | 229 | 229 | 188 | 188 | 173 | 173 | 194 | 194 | 212 | 212 | 195 | 195 | 223 | 223 | 207 | 207 |
| N55 | 101 | 101 | 177 | 177 | 203 | 203 | 157 | 157 | 187 | 187 | 196 | 196 | 229 | 229 | 188 | 188 | 173 | 173 | 194 | 194 | 212 | 212 | 195 | 207 | 223 | 223 | 188 | 188 |
| N56 | 101 | 101 | 177 | 177 | 203 | 203 | 157 | 157 | 187 | 187 | 196 | 196 | 229 | 229 | 188 | 188 | 173 | 173 | 194 | 194 | 212 | 212 | 195 | 195 | 223 | 223 | 207 | 207 |
| N57 | 101 | 101 | 177 | 177 | 203 | 203 | 157 | 157 | 187 | 187 | 196 | 196 | 229 | 229 | 188 | 188 | 173 | 173 | 194 | 194 | 212 | 212 | 207 | 207 | 223 | 223 | 188 | 188 |
| N58 | 101 | 101 | 177 | 177 | 203 | 203 | 157 | 157 | 187 | 187 | 196 | 196 | 229 | 229 | 188 | 188 | 173 | 173 | 194 | 194 | 212 | 212 | 195 | 207 | 223 | 223 | 188 | 188 |
| N59 | 101 | 101 | 177 | 177 | 203 | 203 | 157 | 157 | 187 | 187 | 196 | 196 | 229 | 229 | 188 | 188 | 173 | 173 | 194 | 194 | 212 | 212 | 195 | 195 | 223 | 223 | 188 | 188 |
| N60 | 101 | 101 | 177 | 177 | 203 | 203 | 157 | 157 | 187 | 187 | 196 | 196 | 229 | 229 | 188 | 188 | 173 | 173 | 194 | 194 | 212 | 212 | 195 | 195 | 223 | 223 | 188 | 188 |
| N61 | 101 | 101 | 177 | 177 | 203 | 203 | 157 | 157 | 187 | 187 | 196 | 196 | 229 | 229 | 188 | 188 | 173 | 173 | 194 | 194 | 212 | 212 | 195 | 195 | 223 | 223 | -1 | -1 |
| N62 | 101 | 101 | 177 | 177 | 203 | 203 | 157 | 157 | 187 | 187 | 196 | 196 | 229 | 229 | 188 | 188 | 173 | 173 | 194 | 194 | 212 | 212 | 195 | 195 | 223 | 223 | 207 | 207 |
| N63 | 101 | 101 | 177 | 177 | 203 | 203 | 157 | 157 | 187 | 187 | 196 | 196 | 229 | 229 | 188 | 188 | 173 | 173 | 194 | 194 | 212 | 212 | 207 | 207 | 223 | 223 | 207 | 207 |
| N64 | 101 | 101 | 177 | 177 | 203 | 203 | 157 | 157 | 187 | 187 | 196 | 196 | 229 | 229 | 188 | 188 | 173 | 173 | 194 | 194 | 212 | 212 | 195 | 195 | 223 | 223 | 207 | 207 |
| N65 | 101 | 101 | 177 | 177 | 203 | 203 | 157 | 157 | 187 | 187 | 196 | 196 | 229 | 229 | 188 | 188 | 173 | 173 | 194 | 194 | 212 | 212 | 207 | 207 | 223 | 223 | 188 | 188 |
| N66 | 101 | 101 | 177 | 177 | 203 | 203 | 157 | 157 | 187 | 187 | 196 | 196 | 229 | 229 | 188 | 188 | 173 | 173 | 194 | 194 | 212 | 212 | 195 | 195 | 223 | 223 | 188 | 188 |
| N67 | 101 | 101 | 177 | 177 | 203 | 203 | 157 | 157 | 187 | 187 | 196 | 196 | 229 | 229 | 188 | 188 | 173 | 173 | 194 | 194 | 212 | 212 | 207 | 207 | 223 | 223 | -1 | -1 |
| N68 | 101 | 101 | 177 | 177 | 203 | 203 | 157 | 157 | 187 | 187 | 196 | 196 | 229 | 229 | 188 | 188 | 173 | 173 | 194 | 194 | 212 | 212 | 195 | 195 | 223 | 223 | -1 | -1 |
| N69 | 101 | 101 | 177 | 177 | 203 | 203 | 157 | 157 | 187 | 187 | 196 | 196 | 229 | 229 | 188 | 188 | 173 | 173 | 194 | 194 | 212 | 212 | 207 | 207 | 223 | 223 | 207 | 207 |
| N70 | 101 | 101 | 177 | 177 | 203 | 203 | 157 | 157 | 187 | 187 | 196 | 196 | 229 | 229 | 188 | 188 | 173 | 173 | 194 | 194 | 212 | 212 | 195 | 195 | 223 | 223 | -1 | -1 |
| N71 | 101 | 101 | 177 | 177 | 203 | 203 | 157 | 157 | 187 | 187 | 196 | 196 | 229 | 229 | 188 | 188 | 173 | 173 | 194 | 194 | 212 | 212 | 207 | 207 | 223 | 223 | 207 | 207 |
| N72 | 101 | 101 | 177 | 177 | 203 | 203 | 157 | 157 | 187 | 187 | 196 | 196 | 229 | 229 | 188 | 188 | 173 | 173 | 194 | 194 | 212 | 212 | 207 | 207 | 223 | 223 | 188 | 188 |
| N73 | 101 | 101 | 177 | 177 | 203 | 203 | 157 | 157 | 187 | 187 | 196 | 196 | 229 | 229 | 188 | 188 | 173 | 173 | 194 | 194 | 212 | 212 | 195 | 219 | 223 | 223 | 188 | 207 |
| N74 | 101 | 101 | 177 | 177 | 203 | 203 | 157 | 157 | 187 | 187 | 196 | 196 | 229 | 229 | 188 | 188 | 173 | 173 | 194 | 194 | 212 | 212 | 207 | 207 | 223 | 223 | 188 | 207 |
| N75 | 101 | 101 | 177 | 177 | 203 | 203 | 157 | 157 | 187 | 187 | 196 | 196 | 229 | 229 | 188 | 188 | 173 | 173 | 194 | 194 | 212 | 212 | 195 | 207 | 223 | 223 | 188 | 207 |
| N76 | 101 | 101 | 177 | 177 | 203 | 203 | 157 | 157 | 187 | 187 | 196 | 196 | 229 | 229 | 188 | 188 | 173 | 173 | 194 | 194 | 212 | 212 | 195 | 207 | 223 | 223 | 207 | 207 |
| N77 | 101 | 101 | 177 | 177 | 203 | 203 | 157 | 157 | 187 | 187 | 196 | 196 | 229 | 229 | 188 | 188 | 173 | 173 | 194 | 194 | 212 | 212 | 195 | 207 | 223 | 223 | 207 | 207 |
| N78 | 101 | 101 | 177 | 177 | 203 | 203 | 157 | 157 | 187 | 187 | 196 | 196 | 229 | 229 | 188 | 188 | 173 | 173 | 194 | 194 | 212 | 212 | 195 | 207 | 223 | 223 | 207 | 207 |
| N79 | 101 | 101 | 177 | 177 | 203 | 203 | 157 | 157 | 187 | 187 | 196 | 196 | 229 | 229 | 188 | 188 | 173 | 173 | 194 | 194 | 212 | 212 | 195 | 195 | 223 | 223 | 207 | 207 |
| N80 | 101 | 101 | 177 | 177 | 203 | 203 | 157 | 157 | 187 | 187 | 196 | 196 | 229 | 229 | 188 | 188 | 173 | 173 | 194 | 194 | 212 | 212 | 195 | 207 | 223 | 223 | 207 | 207 |
| N81 | 101 | 101 | 177 | 177 | 203 | 203 | 157 | 157 | 187 | 187 | 196 | 196 | 229 | 229 | 188 | 188 | 173 | 173 | 194 | 194 | 212 | 212 | 195 | 207 | 223 | 223 | 188 | 207 |
| N86 | 101 | 101 | 177 | 177 | 203 | 203 | 157 | 157 | 187 | 187 | 196 | 196 | 229 | 229 | 188 | 188 | 173 | 173 | 194 | 194 | 212 | 212 | 195 | 195 | 223 | 223 | 207 | 207 |
| N87 | 101 | 101 | 177 | 177 | 203 | 203 | 157 | 157 | 187 | 187 | 196 | 196 | 229 | 229 | 188 | 188 | 173 | 173 | 194 | 194 | 212 | 212 | 195 | 195 | 223 | 223 | 188 | 188 |
| N88 | 101 | 101 | 177 | 177 | 203 | 203 | 157 | 157 | 187 | 187 | 196 | 196 | 229 | 229 | 188 | 188 | 173 | 173 | 194 | 194 | 212 | 212 | 195 | 195 | 223 | 223 | 188 | 207 |
| N90 | 101 | 101 | 177 | 177 | 203 | 203 | 157 | 157 | 187 | 187 | 196 | 196 | 229 | 229 | 188 | 188 | 173 | 173 | 194 | 194 | 212 | 212 | 195 | 195 | 223 | 223 | 207 | 207 |
| N92 | 101 | 101 | 177 | 177 | 203 | 203 | 157 | 157 | 187 | 187 | 196 | 196 | 229 | 229 | 188 | 188 | 173 | 173 | 194 | 194 | 212 | 212 | 195 | 195 | 223 | 223 | 188 | 188 |
| N94 | 101 | 101 | 177 | 177 | 203 | 203 | 157 | 157 | 187 | 187 | 196 | 196 | 229 | 229 | 188 | 188 | 173 | 173 | 194 | 194 | 212 | 212 | 195 | 207 | 223 | 223 | 207 | 207 |
| N95 | 101 | 101 | 177 | 177 | 203 | 203 | 157 | 157 | 187 | 187 | 196 | 196 | 229 | 229 | 188 | 188 | 173 | 173 | 194 | 194 | 212 | 212 | 195 | 219 | 223 | 223 | 207 | 207 |
| N96 | 101 | 101 | 177 | 177 | 203 | 203 | 157 | 157 | 187 | 187 | 196 | 196 | 229 | 229 | 188 | 188 | 173 | 173 | 194 | 194 | 212 | 212 | 195 | 219 | 223 | 223 | 207 | 207 |
| N97 | 101 | 101 | 177 | 177 | 203 | 203 | 157 | 157 | 187 | 187 | 196 | 196 | 229 | 229 | 188 | 188 | 173 | 173 | 194 | 194 | 212 | 212 | 195 | 219 | 223 | 223 | 188 | 207 |
| N98 | 101 | 101 | 177 | 177 | 203 | 203 | 157 | 157 | 187 | 187 | 196 | 196 | 229 | 229 | 188 | 188 | 173 | 173 | 194 | 194 | 212 | 212 | 195 | 207 | 223 | 223 | 188 | 207 |
| N99 | 101 | 101 | 177 | 177 | 203 | 203 | 157 | 157 | 187 | 187 | 196 | 196 | 229 | 229 | 188 | 188 | 173 | 173 | 194 | 194 | 212 | 212 | 195 | 219 | 223 | 223 | 188 | 207 |
| N100 | 101 | 101 | 177 | 177 | 203 | 203 | 157 | 157 | 187 | 187 | 196 | 196 | 229 | 229 | 188 | 188 | 173 | 173 | 194 | 194 | 212 | 212 | 207 | 219 | 223 | 223 | 207 | 207 |
| N101 | 101 | 101 | 177 | 177 | 203 | 203 | 157 | 157 | 187 | 187 | 196 | 196 | 229 | 229 | 188 | 188 | 173 | 173 | 194 | 194 | 212 | 212 | 195 | 219 | 223 | 223 | 188 | 188 |
| N102 | 101 | 101 | 177 | 177 | 203 | 203 | 157 | 157 | 187 | 187 | 196 | 196 | 229 | 229 | 188 | 188 | 173 | 173 | 194 | 194 | 212 | 212 | 195 | 195 | 223 | 223 | 188 | 207 |
| N103 | 101 | 101 | 177 | 177 | 203 | 203 | 157 | 157 | 187 | 187 | 196 | 196 | 229 | 229 | 188 | 188 | 173 | 173 | 194 | 194 | 212 | 212 | 207 | 207 | 223 | 223 | -1 | -1 |
| N104 | 101 | 101 | 177 | 177 | 203 | 203 | 157 | 157 | 187 | 187 | 196 | 196 | 229 | 229 | 188 | 188 | 173 | 173 | 194 | 194 | 212 | 212 | 207 | 207 | 223 | 223 | 188 | 207 |
| N105 | 101 | 101 | 177 | 177 | 203 | 203 | 157 | 157 | 187 | 187 | 196 | 196 | 229 | 229 | 188 | 188 | 173 | 173 | 194 | 194 | 212 | 212 | 207 | 207 | 223 | 223 | 207 | 207 |
| N106 | 101 | 101 | 177 | 177 | 203 | 203 | 157 | 157 | 187 | 187 | 196 | 196 | 229 | 229 | 188 | 188 | 173 | 173 | 194 | 194 | 212 | 212 | 195 | 207 | 223 | 223 | 188 | 207 |
| N107 | 101 | 101 | 177 | 177 | 203 | 203 | 157 | 157 | 187 | 187 | 196 | 196 | 229 | 229 | 188 | 188 | 173 | 173 | 194 | 194 | 212 | 212 | 195 | 195 | 223 | 223 | 207 | 207 |
| N108 | 101 | 101 | 177 | 177 | 203 | 203 | 157 | 157 | 187 | 187 | 196 | 196 | 229 | 229 | 188 | 188 | 173 | 173 | 194 | 194 | 212 | 212 | 195 | 195 | 223 | 223 | 207 | 207 |
| N109 | 101 | 101 | 177 | 177 | 203 | 203 | 157 | 157 | 187 | 187 | 196 | 196 | 229 | 229 | 188 | 188 | 173 | 173 | 194 | 194 | 212 | 212 | 195 | 207 | 223 | 223 | 207 | 207 |
| N110 | 101 | 101 | 177 | 177 | 203 | 203 | 157 | 157 | 187 | 187 | 196 | 196 | 229 | 229 | 188 | 188 | 173 | 173 | 194 | 194 | 212 | 212 | 207 | 219 | 223 | 223 | 188 | 207 |
| N111 | 101 | 101 | 177 | 177 | 203 | 203 | 157 | 157 | 187 | 187 | 196 | 196 | 229 | 229 | 188 | 188 | 173 | 173 | 194 | 194 | 212 | 212 | 195 | 207 | 223 | 223 | 188 | 188 |
| N112 | 101 | 101 | 177 | 177 | 203 | 203 | 157 | 157 | 187 | 187 | 196 | 196 | 229 | 229 | 188 | 188 | 173 | 173 | 194 | 194 | 212 | 212 | 195 | 207 | 223 | 223 | 188 | 207 |
| N113 | 101 | 101 | 177 | 177 | 203 | 203 | 157 | 157 | 187 | 187 | 196 | 196 | 229 | 229 | 188 | 188 | 173 | 173 | 194 | 194 | 212 | 212 | 195 | 195 | 223 | 223 | 188 | 207 |
| N114 | 101 | 101 | 177 | 177 | 203 | 203 | 157 | 157 | 187 | 187 | 196 | 196 | 229 | 229 | 188 | 188 | 173 | 173 | 194 | 194 | 212 | 212 | 195 | 195 | 223 | 223 | 188 | 207 |
| N115 | 101 | 101 | 177 | 177 | 203 | 203 | 157 | 157 | 187 | 187 | 196 | 196 | 229 | 229 | 188 | 188 | 173 | 173 | 194 | 194 | 212 | 212 | 195 | 207 | 223 | 223 | 188 | 207 |
| N116 | 101 | 101 | 177 | 177 | 203 | 203 | 157 | 157 | 187 | 187 | 196 | 196 | 229 | 229 | 188 | 188 | -1 | -1 | -1 | -1 | 212 | 212 | 195 | 207 | 223 | 223 | 188 | 207 |
| N117 | 101 | 101 | 177 | 177 | 203 | 203 | 157 | 157 | 187 | 187 | 196 | 196 | 229 | 229 | 188 | 188 | 173 | 173 | 194 | 194 | 212 | 212 | 195 | 195 | 223 | 223 | 188 | 207 |
| NO1B | 101 | 101 | 177 | 177 | 203 | 203 | 157 | 157 | 187 | 187 | 196 | 196 | 229 | 229 | 188 | 188 | 173 | 173 | 194 | 194 | 212 | 212 | 195 | 195 | 223 | 223 | 207 | 207 |
| NO1C | 101 | 101 | 177 | 177 | 203 | 203 | 157 | 157 | 187 | 187 | 196 | 196 | 229 | 229 | 188 | 188 | 173 | 173 | 194 | 194 | 212 | 212 | 195 | 195 | 223 | 223 | 188 | 188 |
| NO1D | 101 | 101 | 177 | 177 | 203 | 203 | 157 | 157 | 187 | 187 | 196 | 196 | 229 | 229 | 188 | 188 | 173 | 173 | 194 | 194 | 212 | 212 | 195 | 195 | 223 | 223 | 207 | 207 |
| NO1E | 101 | 101 | 177 | 177 | 203 | 203 | 157 | 157 | 187 | 187 | 196 | 196 | 229 | 229 | 188 | 188 | 173 | 173 | 194 | 194 | 212 | 212 | 207 | 207 | 223 | 223 | 207 | 207 |
| NO1F | 101 | 101 | 177 | 177 | 203 | 203 | 157 | 157 | 187 | 187 | 194 | 194 | 229 | 229 | 188 | 188 | 173 | 173 | 194 | 194 | 212 | 212 | 195 | 195 | 223 | 223 | 207 | 207 |
| NO1G | 101 | 101 | 177 | 177 | 203 | 203 | 157 | 157 | 187 | 187 | 196 | 196 | 229 | 229 | 188 | 188 | 173 | 173 | 194 | 194 | 212 | 212 | 219 | 219 | 223 | 223 | 207 | 207 |
| NO1H | 101 | 101 | 177 | 177 | 203 | 203 | 157 | 157 | 187 | 187 | 196 | 196 | 229 | 229 | 188 | 188 | 173 | 173 | 194 | 194 | 212 | 212 | 219 | 219 | 223 | 223 | 188 | 188 |
| NO1I | 101 | 101 | 177 | 177 | 203 | 203 | 157 | 157 | 187 | 187 | 196 | 196 | 229 | 229 | 188 | 188 | 173 | 173 | 194 | 194 | 212 | 212 | 195 | 195 | 223 | 223 | 207 | 207 |
| NO4B | 101 | 101 | 177 | 177 | 203 | 203 | 157 | 157 | 187 | 187 | 196 | 196 | 229 | 229 | 188 | 188 | 173 | 173 | 194 | 194 | 212 | 212 | 207 | 207 | 223 | 223 | 188 | 207 |
| NO4C | 101 | 101 | 177 | 177 | 203 | 203 | 157 | 157 | 187 | 187 | 196 | 196 | 229 | 229 | 188 | 188 | 173 | 173 | 194 | 194 | 212 | 212 | 195 | 207 | 223 | 223 | 207 | 207 |
| NO4D | 101 | 101 | 177 | 177 | 203 | 203 | 157 | 157 | 187 | 187 | 196 | 196 | 229 | 229 | 188 | 188 | 173 | 173 | 194 | 194 | 212 | 212 | 195 | 219 | 223 | 223 | 207 | 207 |
| NO4E | 101 | 101 | 177 | 177 | 203 | 203 | 157 | 157 | 187 | 187 | 194 | 196 | 229 | 229 | 188 | 188 | 173 | 173 | 194 | 194 | 212 | 212 | 195 | 195 | 223 | 223 | 207 | 207 |
| NO4F | 101 | 101 | 177 | 177 | 203 | 203 | 157 | 157 | 187 | 187 | 196 | 196 | 229 | 229 | 188 | 188 | 173 | 173 | 194 | 194 | 212 | 212 | 195 | 195 | 223 | 223 | 188 | 207 |
| NO4G | 101 | 101 | 177 | 177 | 203 | 203 | 157 | 157 | 187 | 187 | 196 | 196 | 229 | 229 | 188 | 188 | 173 | 173 | 194 | 194 | 212 | 212 | 195 | 207 | 223 | 223 | 188 | 207 |
| NO4H | 101 | 101 | 177 | 177 | 203 | 203 | 157 | 157 | 187 | 187 | 196 | 196 | 229 | 229 | 188 | 188 | 173 | 173 | 194 | 194 | 212 | 212 | 207 | 219 | 223 | 223 | 188 | 207 |
| NO4I | 101 | 101 | 177 | 177 | 203 | 203 | 157 | 157 | 187 | 187 | 196 | 196 | 229 | 229 | 188 | 188 | 173 | 173 | 194 | 194 | 212 | 212 | 195 | 207 | 223 | 223 | 188 | 207 |
| Q2 | 101 | 101 | 177 | 177 | 203 | 203 | 157 | 157 | 187 | 187 | 196 | 196 | 229 | 229 | 188 | 188 | 173 | 173 | 194 | 194 | 212 | 212 | 195 | 195 | 223 | 223 | 207 | 207 |
| Q3 | 101 | 101 | 177 | 177 | 203 | 203 | 157 | 157 | 187 | 187 | 196 | 196 | 229 | 229 | 188 | 188 | 173 | 173 | 194 | 194 | 212 | 212 | 195 | 195 | 223 | 223 | 207 | 207 |
| Q4 | 101 | 101 | 177 | 177 | 203 | 203 | 157 | 157 | 187 | 187 | 196 | 196 | 229 | 229 | 188 | 188 | 173 | 173 | 194 | 194 | 212 | 212 | 195 | 195 | 223 | 223 | 207 | 207 |
| Q5 | 101 | 101 | 177 | 177 | 203 | 203 | 157 | 157 | 187 | 187 | 196 | 196 | 229 | 229 | 188 | 188 | 173 | 173 | 194 | 194 | 212 | 212 | 195 | 207 | 223 | 223 | 188 | 188 |
| Q6 | 101 | 101 | 177 | 177 | 203 | 203 | 157 | 157 | 187 | 187 | 196 | 196 | 229 | 229 | 188 | 188 | 173 | 173 | 194 | 194 | 212 | 212 | 207 | 207 | 223 | 223 | 207 | 207 |
| Q7 | 101 | 101 | 177 | 177 | 203 | 203 | 157 | 157 | 187 | 187 | 196 | 196 | 229 | 229 | 188 | 188 | 173 | 173 | 194 | 194 | 212 | 212 | 207 | 207 | 223 | 223 | 207 | 207 |
| Q8 | 101 | 101 | 177 | 177 | 203 | 203 | 157 | 157 | 187 | 187 | 196 | 196 | 229 | 229 | 188 | 188 | 173 | 173 | 194 | 194 | 212 | 212 | 207 | 207 | 223 | 223 | 188 | 207 |
| Q9 | 101 | 101 | 177 | 177 | 203 | 203 | 157 | 157 | 187 | 187 | 196 | 196 | 229 | 229 | 188 | 188 | 173 | 173 | 194 | 194 | 212 | 212 | 207 | 207 | 223 | 223 | 207 | 207 |
| Q10 | 101 | 101 | 177 | 177 | 203 | 203 | 157 | 157 | 187 | 187 | 196 | 196 | 229 | 229 | 188 | 188 | 173 | 173 | 194 | 194 | 212 | 212 | 207 | 207 | 223 | 223 | 207 | 207 |
| Q11 | 101 | 101 | 177 | 177 | 203 | 203 | 157 | 157 | 187 | 187 | 196 | 196 | 229 | 229 | 188 | 188 | 173 | 173 | 194 | 194 | 212 | 212 | 207 | 207 | 223 | 223 | 188 | 207 |
| Q12 | 101 | 101 | 177 | 177 | 203 | 203 | 157 | 157 | 187 | 187 | 196 | 196 | 229 | 229 | 188 | 188 | 173 | 173 | 194 | 194 | 212 | 212 | 207 | 207 | 223 | 223 | 188 | 207 |
| Q13 | 101 | 101 | 177 | 177 | 203 | 203 | 157 | 157 | 187 | 187 | 196 | 196 | 229 | 229 | 188 | 188 | 173 | 173 | 194 | 194 | 212 | 212 | 195 | 195 | 223 | 223 | 188 | 207 |
| Q14 | 101 | 101 | 177 | 177 | 203 | 203 | 157 | 157 | 187 | 187 | 196 | 196 | 229 | 229 | 188 | 188 | 173 | 173 | 194 | 194 | 212 | 212 | 195 | 207 | 223 | 223 | 207 | 207 |
| Q15 | 101 | 101 | 177 | 177 | 203 | 203 | 157 | 157 | 187 | 187 | 196 | 196 | 229 | 229 | 188 | 188 | 173 | 173 | 194 | 194 | 212 | 212 | 207 | 207 | 223 | 223 | 207 | 207 |
| Q16 | 101 | 101 | 177 | 177 | 203 | 203 | 157 | 157 | 187 | 187 | 196 | 196 | 229 | 229 | 188 | 188 | 173 | 173 | 194 | 194 | 212 | 212 | 207 | 207 | 223 | 223 | 207 | 207 |
| Q17 | 101 | 101 | 177 | 177 | 203 | 203 | 157 | 157 | 187 | 187 | 196 | 196 | 229 | 229 | 188 | 188 | 173 | 173 | 194 | 194 | 212 | 212 | 207 | 207 | 223 | 223 | 207 | 207 |
| Q18 | 101 | 101 | 177 | 177 | 203 | 203 | 157 | 157 | 187 | 187 | 196 | 196 | 229 | 229 | 188 | 188 | 173 | 173 | 194 | 194 | 212 | 212 | 207 | 207 | 223 | 223 | 207 | 207 |
| Q19 | 101 | 101 | 177 | 177 | 203 | 203 | 157 | 157 | 187 | 187 | 196 | 196 | 229 | 229 | 188 | 188 | 173 | 173 | 194 | 194 | 212 | 212 | 195 | 207 | 223 | 223 | 207 | 207 |
| Q4B | 101 | 101 | 177 | 177 | 203 | 203 | 157 | 157 | 187 | 187 | 194 | 196 | 229 | 229 | 188 | 188 | 173 | 173 | 194 | 194 | 212 | 212 | 207 | 207 | 223 | 223 | 207 | 207 |
| Q4C | 101 | 101 | 177 | 177 | 203 | 203 | 157 | 157 | 187 | 187 | 194 | 196 | 229 | 229 | 188 | 188 | 173 | 173 | 194 | 194 | 212 | 212 | 207 | 207 | 223 | 223 | 207 | 207 |
| Q4D | 101 | 101 | 177 | 177 | 203 | 203 | 157 | 157 | 187 | 187 | 194 | 196 | 229 | 229 | 188 | 188 | 173 | 173 | 194 | 194 | 212 | 212 | 207 | 207 | 223 | 223 | 188 | 207 |
| Q4E | 101 | 101 | 177 | 177 | 203 | 203 | 157 | 157 | 187 | 187 | 194 | 196 | 229 | 229 | 188 | 188 | 173 | 173 | 194 | 194 | 212 | 212 | 195 | 207 | 223 | 223 | 207 | 207 |
| Q4F | 101 | 101 | 177 | 177 | 203 | 203 | 157 | 157 | 187 | 187 | 194 | 196 | 229 | 229 | 188 | 188 | 173 | 173 | 194 | 194 | 212 | 212 | 207 | 207 | 223 | 223 | 207 | 207 |
| Q4G | 101 | 101 | 177 | 177 | 203 | 203 | -1 | -1 | 187 | 187 | 194 | 196 | 229 | 229 | 188 | 188 | 173 | 173 | 194 | 194 | 212 | 212 | 207 | 207 | 223 | 223 | 207 | 207 |
| Q4H | 101 | 101 | 177 | 177 | 203 | 203 | -1 | -1 | 187 | 187 | 194 | 196 | 229 | 229 | 188 | 188 | 173 | 173 | 194 | 194 | 212 | 212 | 207 | 207 | 223 | 223 | 207 | 207 |
| Q4I | 101 | 101 | 177 | 177 | 203 | 203 | 157 | 157 | 187 | 187 | 194 | 196 | 229 | 229 | 188 | 188 | 173 | 173 | 194 | 194 | 212 | 212 | 207 | 207 | 223 | 223 | 207 | 207 |
| Q4J | 101 | 101 | 177 | 177 | 203 | 203 | 157 | 157 | 187 | 187 | 194 | 196 | 229 | 229 | 188 | 188 | 173 | 173 | 194 | 194 | 212 | 212 | 207 | 207 | 223 | 223 | 207 | 207 |
| Q5B | 101 | 101 | 177 | 177 | 203 | 203 | 157 | 157 | 187 | 187 | 196 | 196 | 229 | 229 | 188 | 188 | 173 | 173 | 194 | 194 | 212 | 212 | 207 | 207 | 223 | 223 | 207 | 207 |
| Q5C | 101 | 101 | 177 | 177 | 203 | 203 | 157 | 157 | 187 | 187 | 194 | 196 | 229 | 229 | 188 | 188 | 173 | 173 | 194 | 194 | 212 | 212 | 207 | 207 | 223 | 223 | 207 | 207 |
| Q5D | 101 | 101 | 177 | 177 | 203 | 203 | 157 | 157 | 187 | 187 | 196 | 196 | 229 | 229 | 188 | 188 | 173 | 173 | 194 | 194 | 212 | 212 | 195 | 195 | 223 | 223 | 207 | 207 |
| Q5E | 101 | 101 | 177 | 177 | 203 | 203 | 157 | 157 | 187 | 187 | 196 | 196 | 229 | 229 | 188 | 188 | 173 | 173 | 194 | 194 | 212 | 212 | 207 | 207 | 223 | 223 | 207 | 207 |
| Q5F | 101 | 101 | 177 | 177 | 203 | 203 | 157 | 157 | 187 | 187 | 196 | 196 | 229 | 229 | 188 | 188 | 173 | 173 | 194 | 194 | 212 | 212 | 207 | 207 | 223 | 223 | 207 | 207 |
| Q5G | 101 | 101 | 177 | 177 | 203 | 203 | 157 | 157 | 187 | 187 | 196 | 196 | 229 | 229 | 188 | 188 | 173 | 173 | 194 | 194 | 212 | 212 | 195 | 207 | 223 | 223 | 188 | 188 |
| Q6B | 101 | 101 | 177 | 177 | 203 | 203 | 157 | 157 | 187 | 187 | 196 | 196 | 229 | 229 | 188 | 188 | 173 | 173 | 194 | 194 | 212 | 212 | 207 | 207 | 223 | 223 | 207 | 207 |
| Q6C | 101 | 101 | 177 | 177 | 203 | 203 | 157 | 157 | 187 | 187 | 196 | 196 | 229 | 229 | 188 | 188 | 173 | 173 | 194 | 194 | 212 | 212 | 207 | 207 | 223 | 223 | 207 | 207 |
| Q7B | 101 | 101 | 177 | 177 | 203 | 203 | 157 | 157 | 187 | 187 | 196 | 196 | 229 | 229 | 188 | 188 | 173 | 173 | 194 | 194 | 212 | 212 | 207 | 207 | 223 | 223 | 207 | 207 |
| Q7C | 101 | 101 | 177 | 177 | 203 | 203 | 157 | 157 | 187 | 187 | 196 | 196 | 229 | 229 | 188 | 188 | 173 | 173 | 194 | 194 | 212 | 212 | 207 | 207 | 223 | 223 | 207 | 207 |
| Q7D | 101 | 101 | 177 | 177 | 203 | 203 | 157 | 157 | 187 | 187 | 196 | 196 | 229 | 229 | 188 | 188 | 173 | 173 | 194 | 194 | 212 | 212 | 207 | 207 | 223 | 223 | 207 | 207 |
| Q7E | 101 | 101 | 177 | 177 | 203 | 203 | 157 | 157 | 187 | 187 | 196 | 196 | 229 | 229 | 188 | 188 | 173 | 173 | 194 | 194 | 212 | 212 | 207 | 207 | 223 | 223 | 207 | 207 |
| Q7F | 101 | 101 | 177 | 177 | 203 | 203 | 157 | 157 | 187 | 187 | 196 | 196 | 229 | 229 | 188 | 188 | 173 | 173 | 194 | 194 | 212 | 212 | 207 | 207 | 223 | 223 | 207 | 207 |
| Q10B | 101 | 101 | 177 | 177 | 203 | 203 | 157 | 157 | 187 | 187 | 196 | 196 | 229 | 229 | 188 | 188 | 173 | 173 | 194 | 194 | 212 | 212 | 195 | 195 | 223 | 223 | 207 | 207 |
| Q10C | 101 | 101 | 177 | 177 | 203 | 203 | 157 | 157 | 187 | 187 | 196 | 196 | 229 | 229 | 188 | 188 | 173 | 173 | 194 | 194 | 212 | 212 | 195 | 195 | 223 | 223 | 207 | 207 |
| Q10D | 101 | 101 | 177 | 177 | 203 | 203 | 157 | 157 | 187 | 187 | 196 | 196 | 229 | 229 | 188 | 188 | 173 | 173 | 194 | 194 | 212 | 212 | 195 | 195 | 223 | 223 | 188 | 188 |
| Q10E | 101 | 101 | 177 | 177 | 203 | 203 | 157 | 157 | 187 | 187 | 196 | 196 | 229 | 229 | 188 | 188 | 173 | 173 | 194 | 194 | 212 | 212 | 195 | 195 | 223 | 223 | 207 | 207 |
| Q10F | 101 | 101 | 177 | 177 | 203 | 203 | 157 | 157 | 187 | 187 | 196 | 196 | 229 | 229 | 188 | 188 | 173 | 173 | 194 | 194 | 212 | 212 | 195 | 195 | 223 | 223 | 207 | 207 |
| Q10G | 101 | 101 | 177 | 177 | 203 | 203 | 157 | 157 | 187 | 187 | 196 | 196 | 229 | 229 | 188 | 188 | 173 | 173 | 194 | 194 | 212 | 212 | 195 | 195 | 223 | 223 | 207 | 207 |
| Q12B | 101 | 101 | 177 | 177 | 203 | 203 | 157 | 157 | 187 | 187 | 196 | 196 | 229 | 229 | 188 | 188 | 173 | 173 | 194 | 194 | 212 | 212 | 207 | 207 | 223 | 223 | 207 | 207 |
| Q16B | 101 | 101 | 177 | 177 | 203 | 203 | 157 | 157 | 187 | 187 | 196 | 196 | 229 | 229 | 188 | 188 | 173 | 173 | 194 | 194 | 212 | 212 | 193 | 193 | 223 | 223 | 207 | 207 |
| Q16C | 101 | 101 | 177 | 177 | 203 | 203 | 157 | 157 | 187 | 187 | 196 | 196 | 229 | 229 | 188 | 188 | 173 | 173 | 194 | 194 | 212 | 212 | 195 | 195 | 223 | 223 | 207 | 207 |
| Q18B | 101 | 101 | 177 | 177 | 203 | 203 | 157 | 157 | 187 | 187 | 196 | 196 | 229 | 229 | 188 | 188 | 173 | 173 | 194 | 194 | 212 | 212 | 207 | 207 | 223 | 223 | 207 | 207 |
| Q18C | 101 | 101 | 177 | 177 | 203 | 203 | 157 | 157 | 187 | 187 | 196 | 196 | 229 | 229 | 188 | 188 | 173 | 173 | 194 | 194 | 212 | 212 | 195 | 195 | 223 | 223 | 207 | 207 |
| Q18D | 101 | 101 | 177 | 177 | 203 | 203 | 157 | 157 | 187 | 187 | 196 | 196 | 229 | 229 | 188 | 188 | 173 | 173 | 194 | 194 | 212 | 212 | 207 | 207 | 223 | 223 | 207 | 207 |
| Q19B | 101 | 101 | 177 | 177 | 203 | 203 | 157 | 157 | 187 | 187 | 196 | 196 | 229 | 229 | 188 | 188 | 173 | 173 | 194 | 194 | 212 | 212 | 207 | 207 | 223 | 223 | 207 | 207 |
| S1 | 101 | 101 | 177 | 177 | 203 | 203 | 157 | 157 | 187 | 187 | 196 | 196 | 229 | 229 | 188 | 188 | 173 | 173 | 194 | 194 | 212 | 212 | 195 | 219 | 223 | 223 | 207 | 207 |
| S2 | 101 | 101 | 177 | 177 | 203 | 203 | 157 | 157 | 187 | 187 | 196 | 196 | 229 | 229 | 188 | 188 | 173 | 173 | 194 | 194 | 212 | 212 | 195 | 195 | 223 | 223 | 207 | 207 |
| S3 | 101 | 101 | 177 | 177 | 203 | 203 | 157 | 157 | 187 | 187 | 196 | 196 | 229 | 229 | 188 | 188 | 173 | 173 | 194 | 194 | 212 | 212 | 195 | 195 | 223 | 223 | 207 | 207 |
| S4 | 101 | 101 | 177 | 177 | 203 | 203 | 157 | 157 | 187 | 187 | 196 | 196 | 229 | 229 | 188 | 188 | 173 | 173 | 194 | 194 | 212 | 212 | 195 | 195 | 223 | 223 | 207 | 207 |
| S5 | 101 | 101 | 177 | 177 | 203 | 203 | 157 | 157 | 187 | 187 | 196 | 196 | 229 | 229 | 188 | 188 | 173 | 173 | 194 | 194 | 212 | 212 | 195 | 195 | 223 | 223 | 207 | 207 |
| S6 | 101 | 101 | 177 | 177 | 203 | 203 | 157 | 157 | 187 | 187 | 196 | 196 | 229 | 229 | 188 | 188 | 173 | 173 | 194 | 194 | 212 | 212 | 195 | 195 | 223 | 223 | 207 | 207 |
| S7 | 101 | 101 | 177 | 177 | 203 | 203 | 157 | 157 | 187 | 187 | 196 | 196 | 229 | 229 | 188 | 188 | 173 | 173 | 194 | 194 | 212 | 212 | 195 | 195 | 223 | 223 | 207 | 207 |
| S8 | 101 | 101 | 177 | 177 | 203 | 203 | 157 | 157 | 187 | 187 | 196 | 196 | 229 | 229 | 188 | 188 | 173 | 173 | 194 | 194 | 212 | 212 | 219 | 219 | 223 | 223 | 207 | 207 |
| S9 | 101 | 101 | 177 | 177 | 203 | 203 | 157 | 157 | 187 | 187 | 196 | 196 | 229 | 229 | 188 | 188 | 173 | 173 | 194 | 194 | 212 | 212 | 195 | 195 | 223 | 223 | 188 | 207 |
| S10 | 101 | 101 | 177 | 177 | 203 | 203 | 157 | 157 | 187 | 187 | 196 | 196 | 229 | 229 | 188 | 188 | 173 | 173 | 194 | 194 | 212 | 212 | 195 | 195 | 223 | 223 | 207 | 207 |
| S11 | 101 | 101 | 177 | 177 | 203 | 203 | 157 | 157 | 187 | 187 | 196 | 196 | 229 | 229 | 188 | 188 | 173 | 173 | 194 | 194 | 212 | 212 | 195 | 195 | 223 | 223 | 207 | 207 |
| S12 | 101 | 101 | 177 | 177 | 203 | 203 | 157 | 157 | 187 | 187 | 196 | 196 | 229 | 229 | 188 | 188 | 173 | 173 | 194 | 194 | 212 | 212 | 207 | 219 | 223 | 223 | 188 | 207 |
| S13 | 101 | 101 | 177 | 177 | 203 | 203 | 157 | 157 | 187 | 187 | 196 | 196 | 229 | 229 | 188 | 188 | 173 | 173 | 194 | 194 | 212 | 212 | 207 | 219 | 223 | 223 | 207 | 207 |
| S14 | 101 | 101 | 177 | 177 | 203 | 203 | 157 | 157 | 187 | 187 | 196 | 196 | 229 | 229 | 188 | 188 | 173 | 173 | 194 | 194 | 212 | 212 | 195 | 195 | 223 | 223 | 207 | 207 |
| S15 | 101 | 101 | 177 | 177 | 203 | 203 | 157 | 157 | 187 | 187 | 196 | 196 | 229 | 229 | 188 | 188 | 173 | 173 | 194 | 194 | 212 | 212 | 195 | 195 | 223 | 223 | 207 | 207 |
| S16 | 101 | 101 | 177 | 177 | 203 | 203 | 157 | 157 | 187 | 187 | 196 | 196 | 229 | 229 | 188 | 188 | 173 | 173 | 194 | 194 | 212 | 212 | 195 | 195 | 223 | 223 | 207 | 207 |
| S17 | 101 | 101 | 177 | 177 | 203 | 203 | 157 | 157 | 187 | 187 | 196 | 196 | 229 | 229 | 188 | 188 | 173 | 173 | 194 | 194 | 212 | 212 | 195 | 195 | 223 | 223 | 207 | 207 |
| S18 | 101 | 101 | 177 | 177 | 203 | 203 | 157 | 157 | 187 | 187 | 196 | 196 | 229 | 229 | 188 | 188 | 173 | 173 | 194 | 194 | 212 | 212 | 195 | 195 | 223 | 223 | 207 | 207 |
| S19 | 101 | 101 | 177 | 177 | 203 | 203 | 157 | 157 | 187 | 187 | 196 | 196 | 229 | 229 | 188 | 188 | 173 | 173 | 194 | 194 | 212 | 212 | 195 | 195 | 223 | 223 | 207 | 207 |
| S20 | 101 | 101 | 177 | 177 | 203 | 203 | 157 | 157 | 187 | 187 | 196 | 196 | 229 | 229 | 188 | 188 | 173 | 173 | 194 | 194 | 212 | 212 | 195 | 195 | 223 | 223 | 207 | 207 |
| S21 | 101 | 101 | 177 | 177 | 203 | 203 | 157 | 157 | 187 | 187 | 196 | 196 | 229 | 229 | 188 | 188 | 173 | 173 | 194 | 194 | 212 | 212 | 195 | 195 | 223 | 223 | 207 | 207 |
| SA1B | 101 | 101 | 177 | 177 | 203 | 203 | 157 | 157 | 187 | 187 | 196 | 196 | 229 | 229 | 188 | 188 | 173 | 173 | 194 | 194 | 212 | 212 | 195 | 195 | 223 | 223 | 207 | 207 |
| SA1C | 101 | 101 | 177 | 177 | 203 | 203 | 157 | 157 | 187 | 187 | 196 | 196 | 229 | 229 | 188 | 188 | 173 | 173 | 194 | 194 | 212 | 212 | 195 | 219 | 223 | 223 | 207 | 207 |
| SA1D | 101 | 101 | 177 | 177 | 203 | 203 | 157 | 157 | 187 | 187 | 196 | 196 | 229 | 229 | 188 | 188 | 173 | 173 | 194 | 194 | 212 | 212 | 195 | 195 | 223 | 223 | 207 | 207 |
| SA1E | 101 | 101 | 177 | 177 | 203 | 203 | 157 | 157 | 187 | 187 | 196 | 196 | 229 | 229 | 188 | 188 | 173 | 173 | 194 | 194 | 212 | 212 | 207 | 207 | 223 | 223 | 207 | 207 |
| SA1F | 101 | 101 | 177 | 177 | 203 | 203 | 157 | 157 | 187 | 187 | 196 | 196 | 229 | 229 | 188 | 188 | 173 | 173 | 194 | 194 | 212 | 212 | 207 | 219 | 223 | 223 | 207 | 207 |
| SA1G | 101 | 101 | 177 | 177 | 203 | 203 | 157 | 157 | 187 | 187 | 196 | 196 | 229 | 229 | 188 | 188 | 173 | 173 | 194 | 194 | 212 | 212 | 195 | 195 | 223 | 223 | 188 | 207 |
| SA1H | 101 | 101 | 177 | 177 | 203 | 203 | 157 | 157 | 187 | 187 | 196 | 196 | 229 | 229 | 188 | 188 | 173 | 173 | 194 | 194 | 212 | 212 | 195 | 219 | 223 | 223 | 207 | 207 |
| SA1I | 101 | 101 | 177 | 177 | 203 | 203 | 157 | 157 | 187 | 187 | 196 | 196 | 229 | 229 | 188 | 188 | 173 | 173 | 194 | 194 | 212 | 212 | 195 | 195 | 223 | 223 | 207 | 207 |
| S7B | 101 | 101 | 177 | 177 | 203 | 203 | 157 | 157 | 187 | 187 | 196 | 196 | 229 | 229 | 188 | 188 | 173 | 173 | 194 | 194 | 212 | 212 | 195 | 195 | 223 | 223 | 188 | 188 |
| S7C | 101 | 101 | 177 | 177 | 203 | 203 | 157 | 157 | 187 | 187 | 196 | 196 | 229 | 229 | 188 | 188 | 173 | 173 | 194 | 194 | 212 | 212 | 219 | 219 | 223 | 223 | 207 | 207 |
| S7D | 101 | 101 | 177 | 177 | 203 | 203 | 157 | 157 | 187 | 187 | 196 | 196 | 229 | 229 | 188 | 188 | 173 | 173 | 194 | 194 | 212 | 212 | 219 | 219 | 223 | 223 | 207 | 207 |
| S7E | 101 | 101 | 177 | 177 | 203 | 203 | 157 | 157 | 187 | 187 | 196 | 196 | 229 | 229 | 188 | 188 | 173 | 173 | 194 | 194 | 212 | 212 | 195 | 195 | 223 | 223 | 207 | 207 |
| S7F | 101 | 101 | 177 | 177 | 203 | 203 | 157 | 157 | 187 | 187 | 196 | 196 | 229 | 229 | 188 | 188 | 173 | 173 | 194 | 194 | 212 | 212 | 195 | 195 | 223 | 223 | 207 | 207 |
| S7G | 101 | 101 | 177 | 177 | 203 | 203 | 157 | 157 | 187 | 187 | 196 | 196 | 229 | 229 | 188 | 188 | 173 | 173 | 194 | 194 | 212 | 212 | 195 | 195 | 223 | 223 | 207 | 207 |
| S7H | 101 | 101 | 177 | 177 | 203 | 203 | 157 | 157 | 187 | 187 | 196 | 196 | 229 | 229 | 188 | 188 | 173 | 173 | 194 | 194 | 212 | 212 | 195 | 195 | 223 | 223 | 207 | 207 |
| S7I | 101 | 101 | 177 | 177 | 203 | 203 | 157 | 157 | 187 | 187 | 196 | 196 | 229 | 229 | 188 | 188 | 173 | 173 | 194 | 194 | 212 | 212 | 195 | 195 | 223 | 223 | 207 | 207 |
| S7J | 101 | 101 | 177 | 177 | 203 | 203 | 157 | 157 | 187 | 187 | 196 | 196 | 229 | 229 | 188 | 188 | 173 | 173 | 194 | 194 | 212 | 212 | 219 | 219 | 223 | 223 | 207 | 207 |
| S7K | 101 | 101 | 177 | 177 | 203 | 203 | 157 | 157 | 187 | 187 | 196 | 196 | 229 | 229 | 188 | 188 | 173 | 173 | 194 | 194 | 212 | 212 | 195 | 195 | 223 | 223 | 207 | 207 |
| S7L | 101 | 101 | 177 | 177 | 203 | 203 | 157 | 157 | 187 | 187 | 196 | 196 | 229 | 229 | 188 | 188 | 173 | 173 | 194 | 194 | 212 | 212 | 195 | 195 | 223 | 223 | 207 | 207 |
| S7M | 101 | 101 | 177 | 177 | 203 | 203 | 157 | 157 | 187 | 187 | 196 | 196 | 229 | 229 | 188 | 188 | 173 | 173 | 194 | 194 | 212 | 212 | 195 | 195 | 223 | 223 | 207 | 207 |
| S7N | 101 | 101 | 177 | 177 | 203 | 203 | 157 | 157 | 187 | 187 | 196 | 196 | 229 | 229 | 188 | 188 | 173 | 173 | 194 | 194 | 212 | 212 | 219 | 219 | 223 | 223 | 207 | 207 |
| S8B | 101 | 101 | 177 | 177 | 203 | 203 | 157 | 157 | 187 | 187 | 196 | 196 | 229 | 229 | 188 | 188 | 173 | 173 | 194 | 194 | 212 | 212 | 195 | 195 | 223 | 223 | 207 | 207 |
| S8C | 101 | 101 | 177 | 177 | 203 | 203 | 157 | 157 | 187 | 187 | 196 | 196 | 229 | 229 | 188 | 188 | 173 | 173 | 194 | 194 | 212 | 212 | 195 | 195 | 223 | 223 | 207 | 207 |
| S8D | 101 | 101 | 177 | 177 | 203 | 203 | 157 | 157 | 187 | 187 | 196 | 196 | 229 | 229 | 188 | 188 | 173 | 173 | 194 | 194 | 212 | 212 | 195 | 195 | 223 | 223 | 207 | 207 |
| S8E | 101 | 101 | 177 | 177 | 203 | 203 | 157 | 157 | 187 | 187 | 196 | 196 | 229 | 229 | 188 | 188 | 173 | 173 | 194 | 194 | 212 | 212 | 195 | 195 | 223 | 223 | 207 | 207 |
| S8F | 101 | 101 | 177 | 177 | 203 | 203 | 157 | 157 | 187 | 187 | 196 | 196 | 229 | 229 | 188 | 188 | 173 | 173 | 194 | 194 | 212 | 212 | 219 | 219 | 223 | 223 | 207 | 207 |
| S8G | 101 | 101 | 177 | 177 | 203 | 203 | 157 | 157 | 187 | 187 | 196 | 196 | 229 | 229 | 188 | 188 | 173 | 173 | 194 | 194 | 212 | 212 | 195 | 195 | 223 | 223 | 207 | 207 |
| S8H | 101 | 101 | 177 | 177 | 203 | 203 | 157 | 157 | 187 | 187 | 196 | 196 | 229 | 229 | 188 | 188 | 173 | 173 | 194 | 194 | 212 | 212 | 195 | 195 | 223 | 223 | 207 | 207 |
| S8I | 101 | 101 | 177 | 177 | 203 | 203 | 157 | 157 | 187 | 187 | 196 | 196 | 229 | 229 | 188 | 188 | 173 | 173 | 194 | 194 | 212 | 212 | 219 | 219 | 223 | 223 | 207 | 207 |
| S8J | 101 | 101 | 177 | 177 | 203 | 203 | 157 | 157 | 187 | 187 | 196 | 196 | 229 | 229 | 188 | 188 | 173 | 173 | 194 | 194 | 212 | 212 | 195 | 195 | 223 | 223 | 207 | 207 |
| T1 | 101 | 101 | 177 | 177 | 203 | 203 | 157 | 157 | 187 | 187 | 196 | 196 | 229 | 229 | 188 | 188 | 173 | 173 | 194 | 194 | 212 | 212 | 207 | 219 | 223 | 223 | 207 | 207 |
| T2 | 101 | 101 | 177 | 177 | 203 | 203 | 157 | 157 | 187 | 187 | 196 | 196 | 229 | 229 | 188 | 188 | 173 | 173 | 194 | 194 | 212 | 212 | 195 | 219 | 223 | 223 | 188 | 207 |
| T3 | 101 | 101 | 177 | 177 | 203 | 203 | 157 | 157 | 187 | 187 | 196 | 196 | 229 | 229 | 188 | 188 | 173 | 173 | 194 | 194 | 212 | 212 | 195 | 207 | 223 | 223 | 188 | 207 |
| T4 | 101 | 101 | 177 | 177 | 203 | 203 | 157 | 157 | 187 | 187 | 196 | 196 | 229 | 229 | 188 | 188 | 173 | 173 | 194 | 194 | 212 | 212 | 195 | 219 | 223 | 223 | 188 | 207 |
| T6 | 101 | 101 | 177 | 177 | 203 | 203 | 157 | 157 | 187 | 187 | 194 | 194 | 229 | 229 | 188 | 188 | 173 | 173 | 194 | 194 | 212 | 212 | 195 | 195 | 223 | 223 | 188 | 188 |
| T7 | 101 | 101 | 177 | 185 | 203 | 203 | 157 | 157 | 187 | 187 | 196 | 196 | 229 | 229 | 188 | 188 | 173 | 173 | 194 | 194 | 212 | 212 | 195 | 219 | 223 | 223 | 188 | 207 |
| T9 | 101 | 101 | 177 | 177 | 203 | 203 | 157 | 157 | 187 | 187 | 196 | 196 | 229 | 229 | 188 | 188 | 173 | 173 | 194 | 194 | 212 | 212 | 195 | 195 | 223 | 223 | 207 | 207 |
| T11 | 101 | 101 | 177 | 177 | 203 | 203 | 157 | 157 | 187 | 187 | 194 | 196 | 229 | 229 | 188 | 188 | 173 | 173 | 194 | 194 | 212 | 212 | 195 | 219 | 223 | 223 | 188 | 207 |
| T12 | 101 | 101 | 177 | 177 | 203 | 203 | 157 | 157 | 187 | 187 | 196 | 196 | 229 | 229 | 188 | 188 | 173 | 173 | 194 | 194 | 212 | 212 | 195 | 219 | 223 | 223 | 207 | 207 |
| T13 | 101 | 101 | 177 | 177 | 203 | 203 | 157 | 157 | 187 | 187 | 196 | 196 | 229 | 229 | 188 | 188 | 173 | 173 | 194 | 194 | 212 | 212 | 219 | 219 | 223 | 223 | 207 | 207 |
| T14 | 101 | 101 | 177 | 177 | 203 | 203 | 157 | 157 | 187 | 187 | 196 | 196 | 229 | 229 | 188 | 188 | 173 | 173 | 194 | 194 | 212 | 212 | 207 | 219 | 223 | 223 | 207 | 207 |
| T16 | 101 | 101 | 177 | 177 | 203 | 203 | 157 | 157 | 187 | 187 | 196 | 196 | 229 | 229 | 188 | 188 | 173 | 173 | 194 | 194 | 212 | 212 | 195 | 195 | 223 | 223 | 207 | 207 |
| T17 | 101 | 101 | 177 | 177 | 203 | 203 | 157 | 157 | 187 | 187 | 196 | 196 | 229 | 229 | 188 | 188 | 173 | 173 | 194 | 194 | 212 | 212 | 195 | 219 | 223 | 223 | 207 | 207 |
| T18 | 101 | 101 | 177 | 177 | 203 | 203 | 157 | 157 | 187 | 187 | 196 | 196 | 229 | 229 | 188 | 188 | 173 | 173 | 194 | 194 | 212 | 212 | 195 | 219 | 223 | 223 | 207 | 207 |
| T19 | 101 | 101 | 177 | 177 | 203 | 203 | 157 | 157 | 187 | 187 | 196 | 196 | 229 | 229 | 188 | 188 | 173 | 173 | 194 | 194 | 212 | 212 | 195 | 195 | 223 | 223 | 188 | 207 |
| T20 | 101 | 101 | 177 | 177 | 203 | 203 | 157 | 157 | 187 | 187 | 196 | 196 | 229 | 229 | 188 | 188 | 173 | 173 | 194 | 194 | 212 | 212 | 207 | 207 | 223 | 223 | 188 | 207 |
| T21 | 101 | 101 | 177 | 177 | 203 | 203 | 157 | 157 | 187 | 187 | 196 | 196 | 229 | 229 | 188 | 188 | 173 | 173 | 194 | 194 | 212 | 212 | 195 | 207 | 223 | 223 | 207 | 207 |
| T22 | 101 | 101 | 177 | 177 | 203 | 203 | 157 | 157 | 187 | 187 | 196 | 196 | 229 | 229 | 188 | 188 | 173 | 173 | 194 | 194 | 212 | 212 | 219 | 219 | 223 | 223 | 207 | 207 |
| T23 | 101 | 101 | 177 | 177 | 203 | 203 | 157 | 157 | 187 | 187 | 196 | 196 | 229 | 229 | 188 | 188 | 173 | 173 | 194 | 194 | 212 | 212 | 195 | 195 | 223 | 223 | 207 | 207 |
| T24 | 101 | 101 | 177 | 177 | 203 | 203 | 157 | 157 | 187 | 187 | 194 | 194 | 229 | 229 | 188 | 188 | 173 | 173 | 194 | 194 | 212 | 212 | 195 | 195 | 223 | 223 | 207 | 207 |
| T25 | 101 | 101 | 177 | 177 | 203 | 203 | 157 | 157 | 187 | 187 | 194 | 196 | 229 | 229 | 188 | 188 | 173 | 173 | 194 | 194 | 212 | 212 | 195 | 219 | 223 | 223 | 207 | 207 |
| T26 | 101 | 101 | 177 | 177 | 203 | 203 | 157 | 157 | 187 | 187 | 196 | 196 | 229 | 229 | 188 | 188 | 173 | 173 | 194 | 194 | 212 | 212 | 195 | 195 | 223 | 223 | 207 | 207 |
| T27 | 101 | 101 | 177 | 177 | 203 | 203 | 157 | 157 | 187 | 187 | 196 | 196 | 229 | 229 | 188 | 188 | 173 | 173 | 194 | 194 | 212 | 212 | 219 | 219 | 223 | 223 | 207 | 207 |
| T28 | 101 | 101 | 177 | 177 | 203 | 203 | 157 | 157 | 187 | 187 | 196 | 196 | 229 | 229 | 188 | 188 | 173 | 173 | 194 | 194 | 212 | 212 | 195 | 195 | 223 | 223 | 188 | 207 |
| T29 | 101 | 101 | 177 | 177 | 203 | 203 | 157 | 157 | 187 | 187 | 194 | 196 | 229 | 229 | 188 | 188 | 173 | 173 | 194 | 194 | 212 | 212 | 195 | 195 | 223 | 223 | 207 | 207 |
| T30 | 101 | 101 | 177 | 177 | 203 | 203 | 157 | 157 | 187 | 187 | 194 | 196 | 229 | 229 | 188 | 188 | 173 | 173 | 194 | 194 | 212 | 212 | 195 | 195 | 223 | 223 | 207 | 207 |
| T31 | 101 | 101 | 177 | 177 | 203 | 203 | 157 | 157 | 187 | 187 | 196 | 196 | 229 | 229 | 188 | 188 | 173 | 173 | 194 | 194 | 212 | 212 | 219 | 219 | 223 | 223 | 207 | 207 |
| T32 | 101 | 101 | 177 | 177 | 203 | 203 | 157 | 157 | 187 | 187 | 196 | 196 | 229 | 229 | 188 | 188 | 173 | 173 | 194 | 194 | 212 | 212 | 207 | 219 | 223 | 223 | 207 | 207 |
| T33 | 101 | 101 | 177 | 177 | 203 | 203 | 157 | 157 | 187 | 187 | 196 | 196 | 229 | 229 | 188 | 188 | 173 | 173 | 194 | 194 | 212 | 212 | 207 | 207 | 223 | 223 | 207 | 207 |
| T34 | 101 | 101 | 177 | 177 | 203 | 203 | 157 | 157 | 187 | 187 | 196 | 196 | 229 | 229 | 188 | 188 | 173 | 173 | 194 | 194 | 212 | 212 | 207 | 207 | 223 | 223 | 207 | 207 |
| T35 | 101 | 101 | 177 | 177 | 203 | 203 | 157 | 157 | 187 | 187 | 196 | 196 | 229 | 229 | 188 | 188 | 173 | 173 | 194 | 194 | 212 | 212 | 195 | 195 | 223 | 223 | 207 | 207 |
| T36 | 101 | 101 | 177 | 177 | 203 | 203 | 157 | 157 | 187 | 187 | 196 | 196 | 229 | 229 | 188 | 188 | 173 | 173 | 194 | 194 | 212 | 212 | 195 | 195 | 223 | 223 | 207 | 207 |
| T37 | 101 | 101 | 177 | 177 | 203 | 203 | 157 | 157 | 187 | 187 | 196 | 196 | 229 | 229 | 188 | 188 | 173 | 173 | 194 | 194 | 212 | 212 | 195 | 195 | 223 | 223 | 188 | 207 |
| T38 | 101 | 101 | 177 | 177 | 203 | 203 | 157 | 157 | 187 | 187 | 196 | 196 | 229 | 229 | 188 | 188 | 173 | 173 | 194 | 194 | 212 | 212 | 195 | 195 | 223 | 223 | 207 | 207 |
| T39 | 101 | 101 | 177 | 177 | 203 | 203 | 157 | 157 | 187 | 187 | 196 | 196 | 229 | 229 | 188 | 188 | 173 | 173 | 194 | 194 | 212 | 212 | 195 | 207 | 223 | 223 | 207 | 207 |
| T45 | 101 | 101 | 177 | 177 | 203 | 203 | 157 | 157 | 187 | 187 | 196 | 196 | 229 | 229 | 188 | 188 | 173 | 173 | 194 | 194 | 212 | 212 | 195 | 195 | 223 | 223 | 188 | 207 |
| T46 | 101 | 101 | 185 | 185 | 203 | 203 | 157 | 157 | 187 | 187 | 194 | 194 | 229 | 229 | 188 | 188 | 173 | 173 | 194 | 194 | 212 | 212 | 195 | 195 | 223 | 223 | 188 | 188 |
| T47 | 101 | 101 | 185 | 185 | 203 | 203 | 157 | 157 | 187 | 187 | 194 | 196 | 229 | 229 | 188 | 188 | 173 | 173 | 194 | 194 | 212 | 212 | 195 | 195 | 223 | 223 | 188 | 207 |
| T48 | 101 | 101 | 177 | 185 | 203 | 203 | 157 | 157 | 187 | 187 | 196 | 196 | 229 | 229 | 188 | 188 | 173 | 173 | 194 | 194 | 212 | 212 | 195 | 195 | 223 | 223 | 188 | 207 |
| T49 | 101 | 101 | 177 | 177 | 203 | 203 | 157 | 157 | 187 | 187 | 194 | 194 | 229 | 229 | 188 | 188 | 173 | 173 | 194 | 194 | 212 | 212 | 195 | 195 | 223 | 223 | 188 | 207 |
| T50 | 101 | 101 | 177 | 177 | 203 | 203 | 157 | 157 | 187 | 187 | 196 | 196 | 229 | 229 | 188 | 188 | 173 | 173 | 194 | 194 | 212 | 212 | 195 | 195 | 223 | 223 | 207 | 207 |
| T51 | 101 | 101 | 177 | 177 | 203 | 203 | 157 | 157 | 187 | 187 | 196 | 196 | 229 | 229 | 188 | 188 | 173 | 173 | 194 | 194 | 212 | 212 | 195 | 207 | 223 | 223 | 188 | 207 |
| T52 | 101 | 101 | 177 | 177 | 203 | 203 | 157 | 157 | 187 | 187 | 196 | 196 | 229 | 229 | 188 | 188 | 173 | 173 | 194 | 194 | 212 | 212 | 195 | 219 | 223 | 223 | 207 | 207 |
| T53 | 101 | 101 | 177 | 177 | 203 | 203 | 157 | 157 | 187 | 187 | 196 | 196 | 229 | 229 | 188 | 188 | 173 | 173 | 194 | 194 | 212 | 212 | 195 | 219 | 223 | 223 | 188 | 207 |
| T54 | 101 | 101 | 177 | 177 | 203 | 203 | 157 | 157 | 187 | 187 | 196 | 196 | 229 | 229 | 188 | 188 | 173 | 173 | 194 | 194 | 212 | 212 | 195 | 195 | 223 | 223 | 207 | 207 |
| T55 | 101 | 101 | 177 | 177 | 203 | 203 | 157 | 157 | 187 | 187 | 196 | 196 | 229 | 229 | 188 | 188 | 173 | 173 | 194 | 194 | 212 | 212 | 195 | 195 | 223 | 223 | 207 | 207 |
| T56 | 101 | 101 | 177 | 177 | 203 | 203 | 157 | 157 | 187 | 187 | 196 | 196 | 229 | 229 | 188 | 188 | 173 | 173 | 194 | 194 | 212 | 212 | 195 | 195 | 223 | 223 | 188 | 207 |
| T57 | 101 | 101 | 177 | 177 | 203 | 203 | 157 | 157 | 187 | 187 | 196 | 196 | 229 | 229 | 188 | 188 | 173 | 173 | 194 | 194 | 212 | 212 | 195 | 219 | 223 | 223 | 207 | 207 |
| T58 | 101 | 101 | 177 | 177 | 203 | 203 | 157 | 157 | 187 | 187 | 196 | 196 | 229 | 229 | 188 | 188 | 173 | 173 | 194 | 194 | 212 | 212 | 195 | 195 | 223 | 223 | 207 | 207 |
| T59 | 101 | 101 | 177 | 177 | 203 | 203 | 157 | 157 | 187 | 187 | 196 | 196 | 229 | 229 | 188 | 188 | 173 | 173 | 194 | 194 | 212 | 212 | 195 | 195 | 223 | 223 | 188 | 207 |
| T60 | 101 | 101 | 177 | 177 | 203 | 203 | 157 | 157 | 187 | 187 | 194 | 196 | 229 | 229 | 188 | 188 | 173 | 173 | 194 | 194 | 212 | 212 | 195 | 219 | 223 | 223 | 188 | 207 |
| T61 | 101 | 101 | 177 | 177 | 203 | 203 | 157 | 157 | 187 | 187 | 196 | 196 | 229 | 229 | 188 | 188 | 173 | 173 | 194 | 194 | 212 | 212 | 195 | 195 | 223 | 223 | 207 | 207 |
| T62 | 101 | 101 | 177 | 177 | 203 | 203 | 157 | 157 | 187 | 187 | 194 | 196 | 229 | 229 | 188 | 188 | 173 | 173 | 194 | 194 | 212 | 212 | 195 | 207 | 223 | 223 | 188 | 207 |
| T63 | 101 | 101 | 177 | 177 | 203 | 203 | 157 | 157 | 187 | 187 | 196 | 196 | 229 | 229 | 188 | 188 | 173 | 173 | 194 | 194 | 212 | 212 | 195 | 207 | 223 | 223 | 188 | 207 |
| T64 | 101 | 101 | 177 | 177 | 203 | 203 | 157 | 157 | 187 | 187 | 196 | 196 | 229 | 229 | 188 | 188 | 173 | 173 | 194 | 194 | 212 | 212 | 195 | 207 | 223 | 223 | 188 | 207 |
| T65 | 101 | 101 | 177 | 177 | 203 | 203 | 157 | 157 | 187 | 187 | 194 | 196 | 229 | 229 | 188 | 188 | 173 | 173 | 194 | 194 | 212 | 212 | 195 | 195 | 223 | 223 | 207 | 207 |
| T66 | 101 | 101 | 177 | 185 | 203 | 203 | 157 | 157 | 187 | 187 | 194 | 196 | 229 | 229 | 188 | 188 | 173 | 173 | 194 | 194 | 212 | 212 | 195 | 219 | 223 | 223 | 188 | 207 |
| T67 | 101 | 101 | 177 | 185 | 203 | 203 | 157 | 157 | 187 | 187 | 194 | 196 | 229 | 229 | 188 | 188 | 173 | 173 | 194 | 194 | 212 | 212 | 219 | 219 | 223 | 223 | 188 | 207 |
| T68 | 101 | 101 | 177 | 177 | 203 | 203 | 157 | 157 | 187 | 187 | 196 | 196 | 229 | 229 | 188 | 188 | 173 | 173 | 194 | 194 | 212 | 212 | 219 | 219 | 223 | 223 | 188 | 188 |
| T69 | 101 | 101 | 177 | 177 | 203 | 203 | 157 | 157 | 187 | 187 | 196 | 196 | 229 | 229 | 188 | 188 | 173 | 173 | 194 | 194 | 212 | 212 | 219 | 219 | 223 | 223 | 188 | 188 |
| T70 | 101 | 101 | 177 | 177 | 203 | 203 | 157 | 157 | 187 | 187 | 196 | 196 | 229 | 229 | 188 | 188 | 173 | 173 | 194 | 194 | 212 | 212 | 195 | 195 | 223 | 223 | 188 | 188 |
| T71 | 101 | 101 | 177 | 185 | 203 | 203 | 157 | 157 | 187 | 187 | 194 | 196 | 229 | 229 | 188 | 188 | 173 | 173 | 194 | 194 | 212 | 212 | 219 | 219 | 223 | 223 | 188 | 207 |
| T72 | 101 | 101 | 177 | 185 | 203 | 203 | 157 | 157 | 187 | 187 | 196 | 196 | 229 | 229 | 188 | 188 | 173 | 173 | 194 | 194 | 212 | 212 | 195 | 219 | 223 | 223 | 188 | 207 |
| T73 | 101 | 101 | 177 | 177 | 203 | 203 | 157 | 157 | 187 | 187 | -1 | -1 | 229 | 229 | 188 | 188 | 173 | 173 | 194 | 194 | 212 | 212 | 195 | 195 | 223 | 223 | 188 | 207 |
| T74 | 101 | 101 | 177 | 177 | 203 | 203 | 157 | 157 | 187 | 187 | 196 | 196 | 229 | 229 | 188 | 188 | 173 | 173 | 194 | 194 | 212 | 212 | 219 | 219 | 223 | 223 | 207 | 207 |
| T75 | 101 | 101 | 177 | 177 | 203 | 203 | 157 | 157 | 187 | 187 | 196 | 196 | 229 | 229 | 188 | 188 | 173 | 173 | 194 | 194 | 212 | 212 | 219 | 219 | 223 | 223 | 188 | 188 |
| T76 | 101 | 101 | 177 | 177 | 203 | 203 | 157 | 157 | 187 | 187 | 196 | 196 | 229 | 229 | 188 | 188 | 173 | 173 | 194 | 194 | 212 | 212 | 207 | 207 | 223 | 223 | 188 | 188 |
| T77 | 101 | 101 | 177 | 177 | 203 | 203 | 157 | 157 | 187 | 187 | 196 | 196 | 229 | 229 | 188 | 188 | 173 | 173 | 194 | 194 | 212 | 212 | 195 | 195 | 223 | 223 | 207 | 207 |
| T79 | 101 | 101 | 177 | 177 | 203 | 203 | 157 | 157 | 187 | 187 | 194 | 194 | 229 | 229 | 188 | 188 | 173 | 173 | 194 | 194 | 212 | 212 | 219 | 219 | 223 | 223 | -1 | -1 |
| V1 | 101 | 101 | 177 | 177 | 203 | 203 | 157 | 157 | 187 | 187 | 196 | 196 | 229 | 229 | 188 | 188 | 173 | 173 | 194 | 194 | 212 | 212 | 195 | 195 | 223 | 223 | 207 | 207 |
| V2 | 101 | 101 | 177 | 177 | 203 | 203 | 157 | 157 | 187 | 187 | 196 | 196 | 229 | 229 | 188 | 188 | 173 | 173 | 194 | 194 | 212 | 212 | 195 | 195 | 223 | 223 | 188 | 207 |
| V11 | 101 | 101 | 177 | 177 | 203 | 203 | 157 | 157 | 187 | 187 | 196 | 196 | 229 | 229 | 188 | 188 | 173 | 173 | 194 | 194 | 212 | 212 | 195 | 195 | 223 | 223 | 207 | 207 |
| V20 | 101 | 101 | 177 | 177 | 203 | 203 | 157 | 157 | 187 | 187 | 196 | 196 | 229 | 229 | 188 | 188 | 173 | 173 | 194 | 194 | 212 | 212 | 195 | 207 | 223 | 223 | 188 | 207 |
| V64 | 101 | 101 | 177 | 177 | 203 | 203 | 157 | 157 | 187 | 187 | 196 | 196 | 229 | 229 | 188 | 188 | 173 | 173 | 194 | 194 | 212 | 212 | 207 | 207 | 223 | 223 | 207 | 207 |
| V86 | 101 | 101 | 177 | 177 | 203 | 203 | 157 | 157 | 187 | 187 | 196 | 196 | 229 | 229 | 188 | 188 | 173 | 173 | 194 | 194 | 212 | 212 | 195 | 195 | 223 | 223 | 188 | 207 |
| V120 | 101 | 101 | 177 | 177 | 203 | 203 | 157 | 157 | 187 | 187 | 196 | 196 | 229 | 229 | 188 | 188 | 173 | 173 | 194 | 194 | 212 | 212 | 195 | 195 | 223 | 223 | 207 | 207 |
| V198 | 101 | 101 | 177 | 177 | 203 | 203 | 157 | 157 | 187 | 187 | 196 | 196 | 229 | 229 | 188 | 188 | 173 | 173 | 194 | 194 | 212 | 212 | 195 | 195 | 223 | 223 | 207 | 207 |
| V201 | 101 | 101 | 177 | 177 | 203 | 203 | 157 | 157 | 187 | 187 | 196 | 196 | 229 | 229 | 188 | 188 | 173 | 173 | 194 | 194 | 212 | 212 | 207 | 207 | 223 | 223 | 207 | 207 |
| V205 | 101 | 101 | 177 | 177 | 203 | 203 | 157 | 157 | 187 | 187 | 194 | 194 | 229 | 229 | 188 | 188 | 173 | 173 | 194 | 194 | 212 | 212 | 207 | 207 | 223 | 223 | 207 | 207 |
| V206 | 101 | 101 | 177 | 177 | 203 | 203 | 157 | 157 | 187 | 187 | 196 | 196 | 229 | 229 | 188 | 188 | 173 | 173 | 194 | 194 | 212 | 212 | 195 | 195 | 223 | 223 | 207 | 207 |
| V216 | 101 | 101 | 177 | 177 | 203 | 203 | 157 | 157 | 187 | 187 | 194 | 194 | 229 | 229 | 188 | 188 | 173 | 173 | 194 | 194 | 212 | 212 | 195 | 195 | 223 | 223 | 188 | 188 |
| V226 | 101 | 101 | 177 | 177 | 203 | 203 | 157 | 157 | 187 | 187 | 196 | 196 | 229 | 229 | 188 | 188 | 173 | 173 | 194 | 194 | 212 | 212 | 195 | 195 | 223 | 223 | 188 | 188 |
| V227 | 101 | 101 | 177 | 177 | 203 | 203 | 157 | 157 | 187 | 187 | 196 | 196 | 229 | 229 | 188 | 188 | 173 | 173 | 194 | 194 | 212 | 212 | 195 | 195 | 223 | 223 | 207 | 207 |
| V232 | 101 | 101 | 177 | 177 | 203 | 203 | 157 | 157 | 187 | 187 | 194 | 194 | 229 | 229 | 188 | 188 | 173 | 173 | 194 | 194 | 212 | 212 | 207 | 207 | 223 | 223 | 207 | 207 |
| V237 | 101 | 101 | 177 | 177 | 203 | 203 | 157 | 157 | 187 | 187 | 196 | 196 | 229 | 229 | 188 | 188 | 173 | 173 | 194 | 194 | 212 | 212 | 207 | 207 | 223 | 223 | 207 | 207 |
| V243 | 101 | 101 | 177 | 177 | 203 | 203 | 157 | 157 | 187 | 187 | 196 | 196 | 229 | 229 | 188 | 188 | 173 | 173 | 194 | 194 | 212 | 212 | 207 | 207 | 223 | 223 | 207 | 207 |
| V244 | 101 | 101 | 177 | 177 | 203 | 203 | 157 | 157 | 187 | 187 | 196 | 196 | 229 | 229 | 188 | 188 | 173 | 173 | 194 | 194 | 212 | 212 | 207 | 207 | 223 | 223 | 207 | 207 |
| V246 | 101 | 101 | 177 | 177 | 203 | 203 | 157 | 157 | 187 | 187 | 196 | 196 | 229 | 229 | 188 | 188 | 173 | 173 | 194 | 194 | 212 | 212 | 207 | 207 | 223 | 223 | 207 | 207 |
| V248 | 101 | 101 | 177 | 177 | 203 | 203 | 157 | 157 | 187 | 187 | 194 | 196 | 229 | 229 | 188 | 188 | 173 | 173 | 194 | 194 | 212 | 212 | 195 | 195 | 223 | 223 | 207 | 207 |
| V249 | 101 | 101 | 177 | 177 | 203 | 203 | 157 | 157 | 187 | 187 | 196 | 196 | 229 | 229 | 188 | 188 | 173 | 173 | 194 | 194 | 212 | 212 | 195 | 195 | 223 | 223 | 207 | 207 |
| V253 | 101 | 101 | 177 | 177 | 203 | 203 | 157 | 157 | 187 | 187 | 196 | 196 | 229 | 229 | 188 | 188 | 173 | 173 | 194 | 194 | 212 | 212 | 195 | 195 | 223 | 223 | 207 | 207 |
| V348 | 101 | 101 | 177 | 177 | 203 | 203 | 157 | 157 | 187 | 187 | 196 | 196 | 229 | 229 | 188 | 188 | 173 | 173 | 194 | 194 | 212 | 212 | 195 | 195 | 223 | 223 | 207 | 207 |
| V349 | 101 | 101 | 177 | 177 | 203 | 203 | 157 | 157 | 187 | 187 | 196 | 196 | 229 | 229 | 188 | 188 | 173 | 173 | 194 | 194 | 212 | 212 | 195 | 207 | 223 | 223 | 207 | 207 |
| V350 | 101 | 101 | 177 | 177 | 203 | 203 | 157 | 157 | 187 | 187 | 194 | 196 | 229 | 229 | 188 | 188 | 173 | 173 | 194 | 194 | 212 | 212 | 195 | 195 | 223 | 223 | 188 | 207 |
| V352 | 101 | 101 | 177 | 177 | 203 | 203 | 157 | 157 | 187 | 187 | 196 | 196 | 229 | 229 | 188 | 188 | 173 | 173 | 194 | 194 | 212 | 212 | 195 | 195 | 223 | 223 | 188 | 188 |
| V353 | 101 | 101 | 177 | 177 | 203 | 203 | 157 | 157 | 187 | 187 | 196 | 196 | 229 | 229 | 188 | 188 | 173 | 173 | 194 | 194 | 212 | 212 | 195 | 207 | 223 | 223 | 188 | 207 |
| V355 | 101 | 101 | 177 | 177 | 203 | 203 | 157 | 157 | 187 | 187 | 196 | 196 | 229 | 229 | 188 | 188 | 173 | 173 | 194 | 194 | 212 | 212 | 195 | 207 | 223 | 223 | 207 | 207 |
| V356 | 101 | 101 | 177 | 177 | 203 | 203 | 157 | 157 | 187 | 187 | 196 | 196 | 229 | 229 | 188 | 188 | 173 | 173 | 194 | 194 | 212 | 212 | 195 | 219 | 223 | 223 | 207 | 207 |
| V357 | 101 | 101 | 177 | 177 | 203 | 203 | 157 | 157 | 187 | 187 | 196 | 196 | 229 | 229 | 188 | 188 | 173 | 173 | 194 | 194 | 212 | 212 | 195 | 195 | 223 | 223 | 188 | 207 |
| V358 | 101 | 101 | 177 | 177 | 203 | 203 | 157 | 157 | 187 | 187 | 196 | 196 | 229 | 229 | 188 | 188 | 173 | 173 | 194 | 194 | 212 | 212 | 195 | 195 | 223 | 223 | 207 | 207 |
| V359 | 101 | 101 | 177 | 177 | 203 | 203 | 157 | 157 | 187 | 187 | 194 | 196 | 229 | 229 | 188 | 188 | 173 | 173 | 194 | 194 | 212 | 212 | 195 | 219 | 223 | 223 | 188 | 207 |
| V360 | 101 | 101 | 177 | 177 | 203 | 203 | 157 | 157 | 187 | 187 | 194 | 196 | 229 | 229 | 188 | 188 | 173 | 173 | 194 | 194 | 212 | 212 | 195 | 195 | 223 | 223 | 207 | 207 |
| V361 | 101 | 101 | 177 | 177 | 203 | 203 | 157 | 157 | 187 | 187 | 194 | 196 | 229 | 229 | 188 | 188 | 173 | 173 | 194 | 194 | 212 | 212 | 195 | 207 | 223 | 223 | 207 | 207 |
| V362 | 101 | 101 | 177 | 177 | 203 | 203 | 157 | 157 | 187 | 187 | 196 | 196 | 229 | 229 | 188 | 188 | 173 | 173 | 194 | 194 | 212 | 212 | 195 | 207 | 223 | 223 | 207 | 207 |
| V363 | 101 | 101 | 177 | 177 | 203 | 203 | 157 | 157 | 187 | 187 | 196 | 196 | 229 | 229 | 188 | 188 | 173 | 173 | 192 | 194 | 212 | 212 | 195 | 195 | 223 | 223 | 207 | 207 |
| V364 | 101 | 101 | 177 | 177 | 203 | 203 | 157 | 157 | 187 | 187 | 196 | 196 | 229 | 229 | 188 | 188 | 173 | 173 | 192 | 194 | 212 | 212 | 195 | 195 | 223 | 223 | -1 | -1 |
| V365 | 101 | 101 | 177 | 177 | 203 | 203 | 157 | 157 | 187 | 187 | 196 | 196 | 229 | 229 | 188 | 188 | 173 | 173 | 194 | 194 | 212 | 212 | 195 | 195 | 223 | 223 | 188 | 188 |
| V366 | 101 | 101 | 177 | 177 | 203 | 203 | 157 | 157 | 187 | 187 | 196 | 196 | 229 | 229 | 188 | 188 | 173 | 173 | 194 | 194 | 212 | 212 | 195 | 219 | 223 | 223 | 207 | 207 |
| V367 | 101 | 101 | 177 | 177 | 203 | 203 | 157 | 157 | 187 | 187 | 196 | 196 | 229 | 229 | 188 | 188 | 173 | 173 | 192 | 194 | 212 | 212 | 195 | 219 | 223 | 223 | 207 | 207 |
| V368 | 101 | 101 | 177 | 177 | 203 | 203 | 157 | 157 | 187 | 187 | 196 | 196 | 229 | 229 | 188 | 188 | 173 | 173 | 192 | 194 | 212 | 212 | 195 | 207 | 223 | 223 | 188 | 207 |
| V375 | 101 | 101 | 177 | 177 | 203 | 203 | 157 | 157 | 187 | 187 | 196 | 196 | 229 | 229 | 188 | 188 | 173 | 173 | 192 | 194 | 212 | 212 | 207 | 207 | 223 | 223 | 188 | 188 |
| V376 | 101 | 101 | 177 | 177 | 203 | 203 | 157 | 157 | 187 | 187 | 196 | 196 | 229 | 229 | 188 | 188 | 173 | 173 | 194 | 194 | 212 | 212 | 207 | 207 | 223 | 223 | 207 | 207 |
| V377 | 101 | 101 | 177 | 177 | 203 | 203 | 157 | 157 | 187 | 187 | 196 | 196 | 229 | 229 | 188 | 188 | 173 | 173 | 192 | 194 | 212 | 212 | 195 | 207 | 223 | 223 | 207 | 207 |
| V378 | 101 | 101 | 177 | 177 | 203 | 203 | 157 | 157 | 187 | 187 | 194 | 196 | 229 | 229 | 188 | 188 | 173 | 173 | 192 | 194 | 212 | 212 | 219 | 195 | 223 | 223 | 207 | 207 |
| V379 | 101 | 101 | 177 | 177 | 203 | 203 | 157 | 157 | 187 | 187 | 196 | 196 | 229 | 229 | 188 | 188 | 173 | 173 | 194 | 194 | 212 | 212 | 219 | 219 | 223 | 223 | 207 | 207 |
| V380 | 101 | 101 | 177 | 177 | 203 | 203 | 157 | 157 | 187 | 187 | 196 | 196 | 229 | 229 | 188 | 188 | 173 | 173 | 194 | 194 | 212 | 212 | 195 | 195 | 223 | 223 | 207 | 207 |
| V381 | 101 | 101 | 177 | 177 | 203 | 203 | 157 | 157 | 187 | 187 | 194 | 196 | 229 | 229 | 188 | 188 | 173 | 173 | 192 | 194 | 212 | 212 | 195 | 195 | 223 | 223 | 207 | 207 |
| V382 | 101 | 101 | 177 | 177 | 203 | 203 | 157 | 157 | 187 | 187 | 196 | 196 | 229 | 229 | 188 | 188 | 173 | 173 | 194 | 194 | 212 | 212 | 195 | 207 | 223 | 223 | 188 | 188 |
| V383 | 101 | 101 | 177 | 177 | 203 | 203 | 157 | 157 | 187 | 187 | 196 | 196 | 229 | 229 | 188 | 188 | 173 | 173 | 194 | 194 | 212 | 212 | 195 | 207 | 223 | 223 | 207 | 207 |
| V384 | 101 | 101 | 177 | 177 | 203 | 203 | 157 | 157 | 187 | 187 | 196 | 196 | 229 | 229 | 188 | 188 | 173 | 173 | 194 | 194 | 212 | 212 | 195 | 207 | 223 | 223 | 207 | 207 |
| V385 | 101 | 101 | 177 | 177 | 203 | 203 | 157 | 157 | 187 | 187 | 196 | 196 | 229 | 229 | 188 | 188 | 173 | 173 | 194 | 194 | 212 | 212 | 195 | 207 | 223 | 223 | 207 | 207 |
| V386 | 101 | 101 | 177 | 177 | 203 | 203 | 157 | 157 | 187 | 187 | 196 | 196 | 229 | 229 | 188 | 188 | 173 | 173 | 194 | 194 | 212 | 212 | 195 | 195 | 223 | 223 | 188 | 207 |
| V387 | 101 | 101 | 177 | 177 | 203 | 203 | 157 | 157 | 187 | 187 | 196 | 196 | 229 | 229 | 188 | 188 | 173 | 173 | 194 | 194 | 212 | 212 | 195 | 195 | 223 | 223 | 207 | 207 |
| V388 | 101 | 101 | 177 | 177 | 203 | 203 | 157 | 157 | 187 | 187 | 196 | 196 | 229 | 229 | 188 | 188 | 173 | 173 | 194 | 194 | 212 | 212 | 207 | 207 | 223 | 223 | 207 | 207 |
| V393 | 101 | 101 | 177 | 177 | 203 | 203 | 157 | 157 | 187 | 187 | 196 | 196 | 229 | 229 | 188 | 188 | 173 | 173 | 194 | 194 | 212 | 212 | 207 | 207 | 223 | 223 | 207 | 207 |
| V394 | 101 | 101 | 177 | 177 | 203 | 203 | 157 | 157 | 187 | 187 | 196 | 196 | 229 | 229 | 188 | 188 | 173 | 173 | 194 | 194 | 212 | 212 | 195 | 195 | 223 | 223 | 207 | 207 |
| V396 | 101 | 101 | 177 | 177 | 203 | 203 | 157 | 157 | 187 | 187 | 196 | 196 | 229 | 229 | 188 | 188 | 173 | 173 | 194 | 194 | 212 | 212 | 207 | 207 | 223 | 223 | 188 | 207 |
| V397 | 101 | 101 | 177 | 185 | 203 | 203 | 157 | 157 | 187 | 187 | 194 | 194 | 229 | 229 | 188 | 188 | 173 | 173 | 194 | 194 | 212 | 212 | 207 | 207 | 223 | 223 | 207 | 207 |
| V403 | 101 | 101 | 177 | 177 | 203 | 203 | 157 | 157 | 187 | 187 | 196 | 196 | 229 | 229 | 188 | 188 | 173 | 173 | 194 | 194 | 212 | 212 | 207 | 219 | 223 | 223 | 188 | 207 |
| V404 | 101 | 101 | 177 | 177 | 203 | 203 | 157 | 157 | 187 | 187 | 196 | 196 | 229 | 229 | 188 | 188 | 173 | 173 | 194 | 194 | 212 | 212 | 195 | 207 | 223 | 223 | 207 | 207 |
| V405 | 101 | 101 | 177 | 177 | 203 | 203 | 157 | 157 | 187 | 187 | 196 | 196 | 229 | 229 | 188 | 188 | 173 | 173 | 194 | 194 | 212 | 212 | 205 | 207 | 223 | 223 | 207 | 207 |
| V406 | 101 | 101 | 177 | 177 | 203 | 203 | 157 | 157 | 187 | 187 | 194 | 196 | 229 | 229 | 188 | 188 | 173 | 173 | 194 | 194 | 212 | 212 | 195 | 195 | 223 | 223 | 207 | 207 |
| V407 | 101 | 101 | 177 | 177 | 203 | 203 | 157 | 157 | 187 | 187 | 194 | 196 | 229 | 229 | 188 | 188 | 173 | 173 | 194 | 194 | 212 | 212 | 195 | 195 | 223 | 223 | 207 | 207 |
| V408 | 101 | 101 | 177 | 177 | 203 | 203 | 157 | 157 | 187 | 187 | 194 | 196 | 229 | 229 | 188 | 188 | 173 | 173 | 194 | 194 | 212 | 212 | 195 | 207 | 223 | 223 | 207 | 207 |
| V410 | 101 | 101 | 177 | 177 | 203 | 203 | 157 | 157 | 187 | 187 | 196 | 196 | 229 | 229 | 188 | 188 | 173 | 173 | 194 | 194 | 212 | 212 | 195 | 207 | 223 | 223 | 207 | 207 |
| V414 | 101 | 101 | 177 | 177 | 203 | 203 | 157 | 157 | 187 | 187 | 196 | 196 | 229 | 229 | 188 | 188 | 173 | 173 | 194 | 194 | 212 | 212 | 195 | 207 | 223 | 223 | 188 | 207 |
| V417 | 101 | 101 | 177 | 177 | 203 | 203 | 157 | 157 | 187 | 187 | 196 | 196 | 229 | 229 | 188 | 188 | 173 | 173 | 194 | 194 | 212 | 212 | 195 | 207 | 223 | 223 | 207 | 207 |
| V420 | 101 | 101 | 177 | 177 | 203 | 203 | 157 | 157 | 187 | 187 | 196 | 196 | 229 | 229 | 188 | 188 | 173 | 173 | 194 | 194 | 212 | 212 | 195 | 195 | 223 | 223 | 207 | 207 |
| V421 | 101 | 101 | 177 | 177 | 203 | 203 | 157 | 157 | 187 | 187 | 196 | 196 | 229 | 229 | 188 | 188 | 173 | 173 | 194 | 194 | 212 | 212 | 195 | 195 | 223 | 223 | 207 | 207 |
| V422 | 101 | 101 | 177 | 177 | 203 | 203 | 157 | 157 | 187 | 187 | 196 | 196 | 229 | 229 | 188 | 188 | 173 | 173 | 194 | 194 | 212 | 212 | 195 | 195 | 223 | 223 | 188 | 207 |
| V425 | 101 | 101 | 177 | 177 | 203 | 203 | 157 | 157 | 187 | 187 | 196 | 196 | 229 | 229 | 188 | 188 | 173 | 173 | 194 | 194 | 212 | 212 | 195 | 207 | 223 | 223 | 188 | 207 |
| V428 | 101 | 101 | 177 | 177 | 203 | 203 | 157 | 157 | 187 | 187 | 196 | 196 | 229 | 229 | 188 | 188 | 173 | 173 | 194 | 194 | 212 | 212 | 195 | 207 | 223 | 223 | 188 | 207 |
| V432 | 101 | 101 | 177 | 177 | 203 | 203 | 157 | 157 | 187 | 187 | 196 | 196 | 229 | 229 | 188 | 188 | 173 | 173 | 194 | 194 | 212 | 212 | 195 | 195 | 223 | 223 | 207 | 207 |
| V433 | 101 | 101 | 177 | 177 | 203 | 203 | 157 | 157 | 187 | 187 | 196 | 196 | 229 | 229 | 188 | 188 | 173 | 173 | 194 | 194 | 212 | 212 | 195 | 195 | 223 | 223 | 207 | 207 |
| V434 | 101 | 101 | 177 | 177 | 203 | 203 | 157 | 157 | 187 | 187 | 196 | 196 | 229 | 229 | 188 | 188 | 173 | 173 | 194 | 194 | 212 | 212 | 195 | 195 | 223 | 223 | 207 | 207 |
| V435 | 101 | 101 | 177 | 177 | 203 | 203 | 157 | 157 | 187 | 187 | 196 | 196 | 229 | 229 | 188 | 188 | 173 | 173 | 194 | 194 | 212 | 212 | 195 | 207 | 223 | 223 | 188 | 207 |
| V437 | 101 | 101 | 177 | 177 | 203 | 203 | 157 | 157 | 187 | 187 | 196 | 196 | 229 | 229 | 188 | 188 | 173 | 173 | 194 | 194 | 212 | 212 | 207 | 219 | 223 | 223 | 207 | 207 |
| V439 | 101 | 101 | 177 | 177 | 203 | 203 | 157 | 157 | 187 | 187 | 194 | 194 | 229 | 229 | 188 | 188 | 173 | 173 | 194 | 194 | 212 | 212 | 207 | 207 | 223 | 223 | 188 | 207 |
| V440 | 101 | 101 | 177 | 177 | 203 | 203 | 157 | 157 | 187 | 187 | 196 | 196 | 229 | 229 | 188 | 188 | 173 | 173 | 194 | 194 | 212 | 212 | 195 | 195 | 223 | 223 | 207 | 207 |
| V441 | 101 | 101 | 177 | 177 | 203 | 203 | 157 | 157 | 187 | 187 | 194 | 194 | 229 | 229 | 188 | 188 | 173 | 173 | 194 | 194 | 212 | 212 | 195 | 195 | 223 | 223 | 207 | 207 |
| V442 | 101 | 101 | 177 | 177 | 203 | 203 | 157 | 157 | 187 | 187 | 196 | 196 | 229 | 229 | 188 | 188 | 173 | 173 | 194 | 194 | 212 | 212 | 195 | 195 | 223 | 223 | 207 | 207 |
| V447 | 101 | 101 | 177 | 177 | 203 | 203 | 157 | 157 | 187 | 187 | 194 | 196 | 229 | 229 | 188 | 188 | 173 | 173 | 194 | 194 | 212 | 212 | 195 | 195 | 223 | 223 | 188 | 188 |
| V448 | 101 | 101 | 177 | 177 | 203 | 203 | 157 | 157 | 187 | 187 | 194 | 194 | 229 | 229 | 188 | 188 | 173 | 173 | 194 | 194 | 212 | 212 | 195 | 195 | 223 | 223 | 188 | 207 |
| V449 | 101 | 101 | 177 | 177 | 203 | 203 | 157 | 157 | 187 | 187 | 196 | 196 | 229 | 229 | 188 | 188 | 173 | 173 | 194 | 194 | 212 | 212 | 195 | 195 | 223 | 223 | 207 | 207 |
| V450 | 101 | 101 | 177 | 177 | 203 | 203 | 157 | 157 | 187 | 187 | 196 | 196 | 229 | 229 | 188 | 188 | 173 | 173 | 194 | 194 | 212 | 212 | 195 | 195 | 223 | 223 | 188 | 188 |
| V451 | 101 | 101 | 177 | 177 | 203 | 203 | 157 | 157 | 187 | 187 | 196 | 196 | 229 | 229 | 188 | 188 | 173 | 173 | 194 | 194 | 212 | 212 | 195 | 195 | 223 | 223 | 188 | 188 |
| V452 | 101 | 101 | 177 | 177 | 203 | 203 | 157 | 157 | 187 | 187 | 196 | 196 | 229 | 229 | 188 | 188 | 173 | 173 | 194 | 194 | 212 | 212 | 195 | 195 | 223 | 223 | 188 | 188 |
| V453 | 101 | 101 | 177 | 177 | 203 | 203 | 157 | 157 | 187 | 187 | 194 | 196 | 229 | 229 | 188 | 188 | 173 | 173 | 194 | 194 | 212 | 212 | 195 | 195 | 223 | 223 | 188 | 207 |
| V454 | 101 | 101 | 177 | 177 | 203 | 203 | 157 | 157 | 187 | 187 | 196 | 196 | 229 | 229 | 188 | 188 | 173 | 173 | 194 | 194 | 212 | 212 | 195 | 195 | 223 | 223 | 188 | 207 |
| V455 | 101 | 101 | 177 | 177 | 203 | 203 | 157 | 157 | 187 | 187 | 196 | 196 | 229 | 229 | 188 | 188 | 173 | 173 | 194 | 194 | 212 | 212 | 195 | 195 | 223 | 223 | 188 | 188 |
| V456 | 101 | 101 | 177 | 177 | 203 | 203 | 157 | 157 | 187 | 187 | 196 | 196 | 229 | 229 | 188 | 188 | 173 | 173 | 194 | 194 | 212 | 212 | 195 | 195 | 223 | 223 | 188 | 188 |
| V457 | 101 | 101 | 177 | 177 | 203 | 203 | 157 | 157 | 187 | 187 | 196 | 196 | 229 | 229 | 188 | 188 | 173 | 173 | 194 | 194 | 212 | 212 | 195 | 195 | 223 | 223 | 188 | 188 |
| V458 | 101 | 101 | 177 | 177 | 203 | 203 | 157 | 157 | 187 | 187 | 196 | 196 | 229 | 229 | 188 | 188 | 173 | 173 | 194 | 194 | 212 | 212 | 195 | 195 | 223 | 223 | 188 | 188 |
| V459 | 101 | 101 | 177 | 177 | 203 | 203 | 157 | 157 | 187 | 187 | 196 | 196 | 229 | 229 | 188 | 188 | 173 | 173 | 194 | 194 | 212 | 212 | 195 | 207 | 223 | 223 | 188 | 188 |
| V460 | 101 | 101 | 177 | 177 | 203 | 203 | 157 | 157 | 187 | 187 | 196 | 196 | 229 | 229 | 188 | 188 | 173 | 173 | 194 | 194 | 212 | 212 | 195 | 207 | 223 | 223 | 188 | 188 |
| V461 | 101 | 101 | 177 | 177 | 203 | 203 | 157 | 157 | 187 | 187 | 196 | 196 | 229 | 229 | 188 | 188 | 173 | 173 | 194 | 194 | 212 | 212 | 195 | 195 | 223 | 223 | 188 | 188 |
| V462 | 101 | 101 | 177 | 177 | 203 | 203 | 157 | 157 | 187 | 187 | 196 | 196 | 229 | 229 | 188 | 188 | 173 | 173 | 194 | 194 | 212 | 212 | 195 | 195 | 223 | 223 | 188 | 207 |
| V463 | 101 | 101 | 177 | 177 | 203 | 203 | 157 | 157 | 187 | 187 | 194 | 196 | 229 | 229 | 188 | 188 | 173 | 173 | 194 | 194 | 212 | 212 | 195 | 195 | 223 | 223 | 207 | 207 |
| V464 | 101 | 101 | 177 | 177 | 203 | 203 | 157 | 157 | 187 | 187 | 196 | 196 | 229 | 229 | 188 | 188 | 173 | 173 | 194 | 194 | 212 | 212 | 195 | 207 | 223 | 223 | 207 | 207 |
| V465 | 101 | 101 | 177 | 177 | 203 | 203 | 157 | 157 | 187 | 187 | 194 | 196 | 229 | 229 | 188 | 188 | 173 | 173 | 194 | 194 | 212 | 212 | 195 | 207 | 223 | 223 | 207 | 207 |
| V466 | 101 | 101 | 177 | 177 | 203 | 203 | 157 | 157 | 187 | 187 | 196 | 196 | 229 | 229 | 188 | 188 | 173 | 173 | 194 | 194 | 212 | 212 | 195 | 207 | 223 | 223 | 207 | 207 |
| V467 | 101 | 101 | 177 | 177 | 203 | 203 | 157 | 157 | 187 | 187 | 194 | 196 | 229 | 229 | 188 | 188 | 173 | 173 | 194 | 194 | 212 | 212 | 195 | 219 | 223 | 223 | 207 | 207 |
| V468 | 101 | 101 | 177 | 177 | 203 | 203 | 157 | 157 | 187 | 187 | 196 | 196 | 229 | 229 | 188 | 188 | 173 | 173 | 194 | 194 | 212 | 212 | 195 | 207 | 223 | 223 | 207 | 207 |
| V469 | 101 | 101 | 177 | 177 | 203 | 203 | 157 | 157 | 187 | 187 | 194 | 196 | 229 | 229 | 188 | 188 | 173 | 173 | 194 | 194 | 212 | 212 | 195 | 195 | 223 | 223 | 188 | 207 |
| V470 | 101 | 101 | 177 | 177 | 203 | 203 | 157 | 157 | 187 | 187 | 196 | 196 | 229 | 229 | 188 | 188 | 173 | 173 | 194 | 194 | 212 | 212 | 207 | 207 | 223 | 223 | 207 | 207 |
| V471 | 101 | 101 | 177 | 177 | 203 | 203 | 157 | 157 | 187 | 187 | 196 | 196 | 229 | 229 | 188 | 188 | 173 | 173 | 194 | 194 | 212 | 212 | 207 | 207 | 223 | 223 | 207 | 207 |
| V472 | 101 | 101 | 177 | 177 | 203 | 203 | 157 | 157 | 187 | 187 | 196 | 196 | 229 | 229 | 188 | 188 | 173 | 173 | 194 | 194 | 212 | 212 | 207 | 207 | 223 | 223 | 207 | 207 |
| V473 | 101 | 101 | 177 | 177 | 203 | 203 | 157 | 157 | 187 | 187 | 196 | 196 | 229 | 229 | 188 | 188 | 173 | 173 | 194 | 194 | 212 | 212 | 207 | 207 | 223 | 223 | 207 | 207 |
| V474 | 101 | 101 | 177 | 177 | 203 | 203 | 157 | 157 | 187 | 187 | 196 | 196 | 229 | 229 | 188 | 188 | 173 | 173 | 194 | 194 | 212 | 212 | 207 | 207 | 223 | 223 | 207 | 207 |
| V475 | 101 | 101 | 177 | 177 | 203 | 203 | 157 | 157 | 187 | 187 | 194 | 196 | 229 | 229 | 188 | 188 | 173 | 173 | 194 | 194 | 212 | 212 | 195 | 207 | 223 | 223 | 207 | 207 |
| V476 | 101 | 101 | 177 | 177 | 203 | 203 | 157 | 157 | 187 | 187 | 194 | 196 | 229 | 229 | 188 | 188 | 173 | 173 | 194 | 194 | 212 | 212 | 207 | 207 | 223 | 223 | 207 | 207 |
| V477 | 101 | 101 | 177 | 177 | 203 | 203 | 157 | 157 | 187 | 187 | 194 | 196 | 229 | 229 | 188 | 188 | 173 | 173 | 194 | 194 | 212 | 212 | 195 | 207 | 223 | 223 | 207 | 207 |
| V478 | 101 | 101 | 177 | 177 | 203 | 203 | 157 | 157 | 187 | 187 | 194 | 196 | 229 | 229 | 188 | 188 | 173 | 173 | 194 | 194 | 212 | 212 | 207 | 207 | 223 | 223 | 207 | 207 |
| V479 | 101 | 101 | 177 | 177 | 203 | 203 | 157 | 157 | 187 | 187 | 196 | 196 | 229 | 229 | 188 | 188 | 173 | 173 | 194 | 194 | 212 | 212 | 207 | 207 | 223 | 223 | 207 | 207 |
| V480 | 101 | 101 | 177 | 177 | 203 | 203 | 157 | 157 | 187 | 187 | 196 | 196 | 229 | 229 | 188 | 188 | 173 | 173 | 194 | 194 | 212 | 212 | 195 | 219 | 223 | 223 | 207 | 207 |
| V481 | 101 | 101 | 177 | 177 | 203 | 203 | 157 | 157 | 187 | 187 | 196 | 196 | 229 | 229 | 188 | 188 | 173 | 173 | 194 | 194 | 212 | 212 | 195 | 207 | 223 | 223 | 188 | 207 |
| V482 | 101 | 101 | 177 | 177 | 203 | 203 | 157 | 157 | 187 | 187 | 196 | 196 | 229 | 229 | 188 | 188 | 173 | 173 | 194 | 194 | 212 | 212 | 195 | 219 | 223 | 223 | 207 | 207 |
| V483 | 101 | 101 | 177 | 177 | 203 | 203 | 157 | 157 | 187 | 187 | 196 | 196 | 229 | 229 | 188 | 188 | 173 | 173 | 194 | 194 | 212 | 212 | 195 | 219 | 223 | 223 | 207 | 207 |
| V484 | 101 | 101 | 177 | 177 | 203 | 203 | 157 | 157 | 187 | 187 | 194 | 196 | 229 | 229 | 188 | 188 | 173 | 173 | 194 | 194 | 212 | 212 | 195 | 195 | 223 | 223 | 207 | 207 |
| V485 | 101 | 101 | 177 | 177 | 203 | 203 | 157 | 157 | 187 | 187 | 196 | 196 | 229 | 229 | 188 | 188 | 173 | 173 | 194 | 194 | 212 | 212 | 195 | 195 | 223 | 223 | 207 | 207 |
| V486 | 101 | 101 | 177 | 177 | 203 | 203 | 157 | 157 | 187 | 187 | 196 | 196 | 229 | 229 | 188 | 188 | 173 | 173 | 194 | 194 | 212 | 212 | 195 | 195 | 223 | 223 | 188 | 207 |
| V487 | 101 | 101 | 177 | 177 | 203 | 203 | 157 | 157 | 187 | 187 | 196 | 196 | 229 | 229 | 188 | 188 | 173 | 173 | 194 | 194 | 212 | 212 | 195 | 195 | 223 | 223 | 188 | 207 |
| V488 | 101 | 101 | 177 | 177 | 203 | 203 | 157 | 157 | 187 | 187 | 196 | 196 | 229 | 229 | 188 | 188 | 173 | 173 | 194 | 194 | 212 | 212 | 195 | 195 | 223 | 223 | -1 | -1 |
| V489 | 101 | 101 | 177 | 177 | 203 | 203 | 157 | 157 | 187 | 187 | 196 | 196 | 229 | 229 | 188 | 188 | 173 | 173 | 194 | 194 | 212 | 212 | 195 | 195 | 223 | 223 | 188 | 207 |
| V490 | 101 | 101 | 177 | 177 | 203 | 203 | 157 | 157 | 187 | 187 | 196 | 196 | 229 | 229 | 188 | 188 | 173 | 173 | 194 | 194 | 212 | 212 | 195 | 195 | 223 | 223 | 188 | 188 |
| V491 | 101 | 101 | 177 | 177 | 203 | 203 | 157 | 157 | 187 | 187 | 196 | 196 | 229 | 229 | 188 | 188 | 173 | 173 | 194 | 194 | 212 | 212 | 195 | 195 | 223 | 223 | 188 | 188 |
| V492 | 101 | 101 | 177 | 177 | 203 | 203 | 157 | 157 | 187 | 187 | 196 | 196 | 229 | 229 | 188 | 188 | 173 | 173 | 194 | 194 | 212 | 212 | 195 | 195 | 223 | 223 | 188 | 207 |
| V493 | 101 | 101 | 177 | 177 | 203 | 203 | 157 | 157 | 187 | 187 | 196 | 196 | 229 | 229 | 188 | 188 | 173 | 173 | 194 | 194 | 212 | 212 | 195 | 195 | 223 | 223 | 188 | 207 |
| V494 | 101 | 101 | 177 | 177 | 203 | 203 | 157 | 157 | 187 | 187 | 196 | 196 | 229 | 229 | 188 | 188 | 173 | 173 | 194 | 194 | 212 | 212 | 195 | 219 | 223 | 223 | 207 | 207 |
| V495 | 101 | 101 | 177 | 177 | 203 | 203 | 157 | 157 | 187 | 187 | 196 | 196 | 229 | 229 | 188 | 188 | 173 | 173 | 194 | 194 | 212 | 212 | 195 | 219 | 223 | 223 | 207 | 207 |
| V496 | 101 | 101 | 177 | 177 | 203 | 203 | 157 | 157 | 187 | 187 | 196 | 196 | 229 | 229 | 188 | 188 | 173 | 173 | 194 | 194 | 212 | 212 | 195 | 207 | 223 | 223 | 188 | 207 |
| V497 | 101 | 101 | 177 | 177 | 203 | 203 | 157 | 157 | 187 | 187 | 196 | 196 | 229 | 229 | 188 | 188 | 173 | 173 | 194 | 194 | 212 | 212 | 195 | 195 | 223 | 223 | 188 | 207 |
| NZS1 | 101 | 101 | 177 | 177 | 203 | 203 | 157 | 157 | 187 | 187 | 194 | 196 | 229 | 229 | 188 | 188 | 173 | 173 | 194 | 194 | 212 | 212 | 195 | 207 | 223 | 223 | 188 | 188 |
| NZS2 | 101 | 101 | 177 | 177 | 203 | 203 | 157 | 157 | 187 | 187 | 194 | 196 | 229 | 229 | 188 | 188 | 173 | 173 | 194 | 194 | 212 | 212 | 195 | 195 | 223 | 223 | 207 | 207 |
| NZS3a | 101 | 101 | 177 | 185 | 203 | 203 | 157 | 157 | 187 | 187 | 196 | 196 | 229 | 229 | 188 | 188 | 173 | 173 | 194 | 194 | 212 | 212 | 195 | 195 | 223 | 223 | 188 | 207 |
| NZS3b | 101 | 101 | 177 | 185 | 203 | 203 | 157 | 157 | 187 | 187 | 194 | 196 | 229 | 229 | 188 | 188 | 173 | 173 | 194 | 194 | 212 | 212 | 195 | 195 | 223 | 223 | 188 | 207 |
| NZS3c | 101 | 101 | 177 | 185 | 203 | 203 | 157 | 157 | 187 | 187 | 196 | 196 | 229 | 229 | 188 | 188 | 173 | 173 | 194 | 194 | 212 | 212 | 207 | 207 | 223 | 223 | 188 | 207 |
| NZS3d | 101 | 101 | 177 | 185 | 203 | 203 | 157 | 157 | 187 | 187 | 194 | 196 | 229 | 229 | 188 | 188 | 173 | 173 | 194 | 194 | 212 | 212 | 195 | 219 | 223 | 223 | 207 | 207 |
| NZS3e | 101 | 101 | 177 | 177 | 203 | 203 | 157 | 157 | 187 | 187 | 196 | 196 | 229 | 229 | 188 | 188 | 173 | 173 | 194 | 194 | 212 | 212 | 207 | 207 | 223 | 223 | 188 | 188 |
| NZS4a | 101 | 101 | 177 | 185 | 203 | 203 | 157 | 157 | 187 | 187 | 194 | 196 | 229 | 229 | 188 | 188 | 173 | 173 | 194 | 194 | 212 | 212 | 195 | 207 | 223 | 223 | 207 | 207 |
| NZS4b | 101 | 101 | 177 | 177 | 203 | 206 | 157 | 157 | 187 | 187 | 196 | 196 | 229 | 229 | 188 | 188 | 173 | 173 | 194 | 194 | 212 | 212 | 195 | 195 | 223 | 223 | 188 | 188 |
| NZS4c | 101 | 101 | 177 | 185 | 203 | 203 | 157 | 157 | 187 | 187 | 194 | 196 | 229 | 229 | 188 | 188 | 173 | 173 | 194 | 194 | 212 | 212 | 195 | 195 | 223 | 223 | 207 | 207 |
| NZS4d | 101 | 101 | 177 | 185 | 203 | 203 | 157 | 157 | 187 | 187 | 196 | 196 | 229 | 229 | 188 | 188 | 173 | 173 | 194 | 194 | 212 | 212 | 195 | 195 | 223 | 223 | 188 | 207 |
| NZS4e | 101 | 101 | 185 | 185 | 203 | 203 | 157 | 157 | 187 | 187 | 196 | 196 | 229 | 229 | 188 | 188 | 173 | 173 | 194 | 194 | 212 | 212 | 219 | 219 | 223 | 223 | 207 | 207 |
| NZS4f | 101 | 101 | 185 | 185 | 203 | 203 | 157 | 157 | 187 | 187 | 196 | 196 | 229 | 229 | 188 | 188 | 173 | 173 | 194 | 194 | 212 | 212 | 195 | 195 | 223 | 223 | 207 | 207 |
| NZS4g | 101 | 101 | 177 | 177 | 203 | 203 | 157 | 157 | 187 | 187 | 194 | 194 | 229 | 229 | 188 | 188 | 173 | 173 | 194 | 194 | 212 | 212 | 195 | 195 | 223 | 223 | 188 | 188 |
| NZS4h | 101 | 101 | 177 | 177 | 203 | 203 | 157 | 157 | 187 | 187 | 196 | 196 | 229 | 229 | 188 | 188 | 173 | 173 | 194 | 194 | 212 | 212 | 195 | 195 | 223 | 223 | 188 | 188 |
| NZS6 | 101 | 101 | 177 | 185 | 203 | 203 | 157 | 157 | 187 | 187 | 196 | 196 | 229 | 229 | 188 | 188 | 173 | 173 | 194 | 194 | 212 | 212 | 195 | 195 | 223 | 223 | 188 | 207 |
| NZS7 | 101 | 101 | 177 | 185 | 203 | 203 | 157 | 157 | 187 | 187 | 196 | 196 | 229 | 229 | 188 | 188 | 173 | 173 | 194 | 194 | 212 | 212 | 195 | 195 | 223 | 223 | 188 | 188 |
| NZS12a | 101 | 101 | 177 | 185 | 203 | 203 | 157 | 157 | 187 | 187 | 196 | 196 | 229 | 229 | -1 | -1 | 173 | 173 | 194 | 194 | 212 | 212 | 195 | 207 | 223 | 223 | 188 | 188 |
| NZS12b | 101 | 101 | 177 | 185 | 203 | 203 | 157 | 157 | 187 | 187 | 196 | 196 | 229 | 229 | 188 | 188 | 173 | 173 | 194 | 194 | 212 | 212 | 195 | 207 | 223 | 223 | 207 | 207 |
| NZS12c | 101 | 101 | 177 | 177 | 203 | 203 | 157 | 157 | 187 | 187 | 196 | 196 | 229 | 229 | 188 | 188 | 173 | 173 | 194 | 194 | 212 | 212 | 207 | 207 | 223 | 223 | 188 | 188 |
| NZS12d | 101 | 101 | 177 | 177 | 203 | 203 | 157 | 157 | 187 | 187 | 196 | 196 | 229 | 229 | 188 | 188 | 173 | 173 | 194 | 194 | 212 | 212 | 195 | 195 | 223 | 223 | 207 | 207 |
| NZS13a | 101 | 101 | 177 | 177 | 203 | 203 | 157 | 157 | 187 | 187 | 194 | 194 | 229 | 229 | 188 | 188 | 173 | 173 | 194 | 194 | 212 | 212 | 195 | 195 | 223 | 223 | 207 | 207 |
| NZS13b | 101 | 101 | 185 | 185 | 203 | 203 | 157 | 157 | 187 | 187 | 196 | 196 | 229 | 229 | 188 | 188 | 173 | 173 | 194 | 194 | 212 | 212 | 207 | 207 | 223 | 223 | 207 | 207 |
| NZS13c | 101 | 101 | 185 | 185 | 203 | 203 | 157 | 157 | 187 | 187 | 196 | 196 | 229 | 229 | 188 | 188 | 173 | 173 | 194 | 194 | 212 | 212 | 195 | 195 | 223 | 223 | 207 | 207 |
| NZS14a | 101 | 101 | 177 | 185 | 203 | 203 | 157 | 157 | 187 | 187 | 196 | 196 | 229 | 229 | 188 | 188 | 173 | 173 | 194 | 194 | 212 | 212 | 195 | 195 | 223 | 223 | 188 | 207 |
| NZS14b | 101 | 101 | 177 | 177 | 203 | 203 | 157 | 157 | 187 | 187 | 196 | 196 | 229 | 229 | 188 | 188 | 173 | 173 | 194 | 194 | 212 | 212 | 195 | 219 | 223 | 223 | 207 | 207 |
| NZS14c | 101 | 101 | 177 | 185 | 203 | 203 | 157 | 157 | 187 | 187 | 196 | 196 | 229 | 229 | 188 | 188 | 173 | 173 | 194 | 194 | 212 | 212 | 195 | 219 | 223 | 223 | 188 | 207 |
| NZS14d | 101 | 101 | 177 | 185 | 203 | 203 | 157 | 157 | 187 | 187 | 196 | 196 | 229 | 229 | 188 | 188 | 173 | 173 | 194 | 194 | 212 | 212 | 195 | 219 | 223 | 223 | 207 | 207 |
| NZS14e | 101 | 101 | 177 | 177 | 203 | 203 | 157 | 157 | 187 | 187 | 196 | 196 | 229 | 229 | 188 | 188 | 173 | 173 | 194 | 194 | 212 | 212 | 195 | 195 | 223 | 223 | 188 | 207 |
| NZS14f | 101 | 101 | 177 | 177 | 203 | 203 | 157 | 157 | 187 | 187 | 196 | 196 | 229 | 229 | 188 | 188 | 173 | 173 | 194 | 194 | 212 | 212 | 195 | 219 | 223 | 223 | 207 | 207 |
| NZS14g | 101 | 101 | 177 | 185 | 203 | 203 | 157 | 157 | 187 | 187 | 196 | 196 | 229 | 229 | 188 | 188 | 173 | 173 | 194 | 194 | 212 | 212 | 195 | 195 | 223 | 223 | 207 | 207 |
| NZS14h | 101 | 101 | 177 | 177 | 203 | 203 | 157 | 157 | 187 | 187 | 196 | 196 | 229 | 229 | 188 | 188 | 173 | 173 | 194 | 194 | 212 | 212 | 195 | 219 | 223 | 223 | 207 | 207 |
| NZS12E | 101 | 101 | 177 | 177 | 203 | 203 | 157 | 157 | 187 | 187 | 196 | 196 | 229 | 229 | 188 | 188 | 173 | 173 | 194 | 194 | 212 | 212 | 195 | 195 | 223 | 223 | 188 | 188 |
| NZS13D | 101 | 101 | 177 | 177 | 203 | 203 | 157 | 157 | 187 | 187 | 196 | 196 | 229 | 229 | 188 | 188 | 173 | 173 | 194 | 194 | 212 | 212 | 195 | 195 | 223 | 223 | 207 | 207 |
| NZS13E | 101 | 101 | 177 | 177 | 203 | 203 | 157 | 157 | 187 | 187 | 196 | 196 | 229 | 229 | 188 | 188 | 173 | 173 | 194 | 194 | 212 | 212 | 195 | 195 | 223 | 223 | 207 | 207 |
| NZS13F | 101 | 101 | 185 | 185 | 203 | 203 | 157 | 157 | 187 | 187 | 196 | 196 | 229 | 229 | 188 | 188 | 173 | 173 | 194 | 194 | 212 | 212 | 195 | 195 | 223 | 223 | 207 | 207 |
| NZS13G | 101 | 101 | 177 | 177 | 203 | 203 | 157 | 157 | 187 | 187 | 196 | 196 | 229 | 229 | 188 | 188 | 173 | 173 | 194 | 194 | 212 | 212 | 207 | 207 | 223 | 223 | 207 | 207 |
| NZS13H | 101 | 101 | 185 | 185 | 203 | 203 | 157 | 157 | 187 | 187 | 196 | 196 | 229 | 229 | 188 | 188 | 173 | 173 | 194 | 194 | 212 | 212 | 219 | 219 | 223 | 223 | 207 | 207 |
| NZS13I | 101 | 101 | 177 | 177 | 203 | 203 | 157 | 157 | 187 | 187 | 196 | 196 | 229 | 229 | 188 | 188 | 173 | 173 | 194 | 194 | 212 | 212 | 195 | 195 | 223 | 223 | 207 | 207 |
| NZS13J | 101 | 101 | 177 | 177 | 203 | 203 | 157 | 157 | 187 | 187 | 196 | 196 | 229 | 229 | 188 | 188 | 173 | 173 | 194 | 194 | 212 | 212 | 195 | 195 | 223 | 223 | 188 | 188 |
| NZS13K | 101 | 101 | 177 | 177 | 203 | 203 | 157 | 157 | 187 | 187 | 196 | 196 | 229 | 229 | 188 | 188 | 173 | 173 | 194 | 194 | 212 | 212 | 195 | 195 | 223 | 223 | 188 | 188 |
| NZS13L | 101 | 101 | 177 | 177 | 206 | 206 | 157 | 157 | 187 | 187 | 194 | 194 | 229 | 229 | 188 | 188 | 173 | 173 | 194 | 194 | 212 | 212 | 219 | 219 | 223 | 223 | 207 | 207 |
| NZS13M | 101 | 101 | 177 | 177 | 203 | 203 | 157 | 157 | 187 | 187 | 196 | 196 | 229 | 229 | 188 | 188 | 173 | 173 | 194 | 194 | 212 | 212 | 195 | 195 | 223 | 223 | 207 | 207 |
| NZS13N | 101 | 101 | 185 | 185 | 203 | 203 | 157 | 157 | 187 | 187 | 196 | 196 | 229 | 229 | 188 | 188 | 173 | 173 | 194 | 194 | 212 | 212 | 195 | 195 | 223 | 223 | 207 | 207 |
| NZS13O | 101 | 101 | 177 | 177 | 203 | 203 | 157 | 157 | 187 | 187 | 196 | 196 | 229 | 229 | 188 | 188 | 173 | 173 | 194 | 194 | 212 | 212 | 195 | 195 | 223 | 223 | 207 | 207 |
| Z1 | 101 | 101 | 177 | 177 | 203 | 203 | 157 | 157 | 187 | 187 | 196 | 196 | 229 | 229 | 188 | 188 | 173 | 173 | 194 | 194 | 212 | 212 | 207 | 207 | 223 | 223 | 207 | 207 |
| Z2 | 101 | 101 | 177 | 185 | 203 | 203 | 157 | 157 | 187 | 187 | 196 | 196 | 229 | 229 | 188 | 188 | 173 | 173 | 194 | 194 | 212 | 212 | 207 | 207 | 223 | 223 | 188 | 207 |
| Z3 | 101 | 101 | 177 | 177 | 203 | 203 | 157 | 157 | 187 | 187 | 194 | 196 | 229 | 229 | 188 | 188 | 173 | 173 | 194 | 194 | 212 | 212 | 195 | 219 | 223 | 223 | 188 | 207 |
| Z4 | 101 | 101 | 177 | 185 | 203 | 203 | 157 | 157 | 187 | 187 | 196 | 196 | 229 | 229 | 188 | 188 | -1 | -1 | 194 | 194 | 212 | 212 | 207 | 207 | 223 | 223 | 207 | 207 |
| Z5 | 101 | 101 | 177 | 177 | 203 | 203 | 157 | 157 | 187 | 187 | 196 | 196 | 229 | 229 | 188 | 188 | 173 | 173 | 194 | 194 | 212 | 212 | 207 | 207 | 223 | 223 | 188 | 207 |
| Z6 | 101 | 101 | 177 | 185 | 203 | 203 | 157 | 157 | 187 | 187 | 196 | 196 | 229 | 229 | 188 | 188 | 173 | 173 | 194 | 194 | 212 | 212 | 195 | 207 | 223 | 223 | 207 | 207 |
| Z7 | 101 | 101 | 177 | 185 | 203 | 203 | 157 | 157 | 187 | 187 | 194 | 196 | 229 | 229 | 188 | 188 | 173 | 173 | 194 | 194 | 212 | 212 | 195 | 207 | 223 | 223 | 207 | 207 |
| Z8 | 101 | 101 | 177 | 177 | 203 | 203 | 157 | 157 | 187 | 187 | 196 | 196 | 229 | 229 | 188 | 188 | 173 | 173 | 194 | 194 | 212 | 212 | 195 | 207 | 223 | 223 | 207 | 207 |
| Z9 | 101 | 101 | 177 | 177 | 203 | 203 | 157 | 157 | 187 | 187 | 196 | 196 | 229 | 229 | 188 | 188 | 173 | 173 | 194 | 194 | 212 | 212 | 195 | 219 | 223 | 223 | 188 | 207 |
| Z10 | 101 | 101 | 177 | 177 | 203 | 203 | 157 | 157 | 187 | 187 | 196 | 196 | 229 | 229 | 188 | 188 | 173 | 173 | 194 | 194 | 212 | 212 | 207 | 207 | 223 | 223 | 188 | 188 |
| Z11 | 101 | 101 | 185 | 185 | 203 | 203 | 157 | 157 | 187 | 187 | 196 | 196 | 229 | 229 | 188 | 188 | 173 | 173 | 194 | 194 | 212 | 212 | 195 | 219 | 223 | 223 | 207 | 207 |
| Z12 | 101 | 101 | 177 | 177 | 203 | 203 | 157 | 157 | 187 | 187 | 196 | 196 | 229 | 229 | 188 | 188 | 173 | 173 | 194 | 194 | 212 | 212 | 207 | 207 | 223 | 223 | 207 | 207 |
| Z13 | 101 | 101 | 177 | 185 | 203 | 203 | 157 | 157 | 187 | 187 | 196 | 196 | 229 | 229 | 188 | 188 | 173 | 173 | 194 | 194 | 212 | 212 | 195 | 207 | 223 | 223 | 207 | 207 |
| Z14 | 101 | 101 | 177 | 177 | 203 | 203 | 157 | 157 | 187 | 187 | 196 | 196 | 229 | 229 | 188 | 188 | 173 | 173 | 194 | 194 | 212 | 212 | 207 | 207 | 223 | 223 | 207 | 207 |
| Z15 | 101 | 101 | 177 | 177 | 203 | 203 | 157 | 157 | 187 | 187 | 196 | 196 | 229 | 229 | 188 | 188 | 173 | 173 | 194 | 194 | 212 | 212 | 195 | 207 | 223 | 223 | 207 | 207 |
| Z16 | 101 | 101 | 177 | 177 | 203 | 203 | 157 | 157 | 187 | 187 | 194 | 196 | 229 | 229 | 188 | 188 | 173 | 173 | 194 | 194 | 212 | 212 | 195 | 219 | 223 | 223 | 188 | 207 |
| NO2B | 101 | 101 | 177 | 177 | 203 | 203 | 157 | 157 | 187 | 187 | 196 | 196 | 229 | 229 | 188 | 188 | 173 | 173 | 194 | 194 | 212 | 212 | 195 | 195 | 223 | 223 | 188 | 207 |
| NO2C | 101 | 101 | 177 | 177 | 203 | 203 | 157 | 157 | 187 | 187 | 196 | 196 | 229 | 229 | 188 | 188 | 173 | 173 | 194 | 194 | 212 | 212 | 207 | 207 | 223 | 223 | 207 | 207 |
| NO2D | 101 | 101 | 177 | 177 | 203 | 203 | 157 | 157 | 187 | 187 | 196 | 196 | 229 | 229 | 188 | 188 | 173 | 173 | 194 | 194 | 212 | 212 | 195 | 195 | 223 | 223 | 207 | 207 |
| NO3B | 101 | 101 | 177 | 177 | 203 | 203 | 157 | 157 | 187 | 187 | 196 | 196 | 229 | 229 | 188 | 188 | 173 | 173 | 194 | 194 | 212 | 212 | 195 | 219 | 223 | 223 | 207 | 207 |
| NO3C | 101 | 101 | 177 | 177 | 203 | 203 | 157 | 157 | 187 | 187 | 196 | 196 | 229 | 229 | 188 | 188 | 173 | 173 | 194 | 194 | 212 | 212 | 195 | 219 | 223 | 223 | 207 | 207 |
| FCNI021495 | 101 | 101 | 177 | 177 | 203 | 203 | 157 | 157 | 187 | 187 | 196 | 196 | 229 | 229 | 188 | 188 | 173 | 173 | 194 | 194 | 212 | 212 | 195 | 207 | 223 | 223 | 188 | 188 |
| FCNI021496 | 101 | 101 | 177 | 177 | 203 | 203 | 157 | 157 | 187 | 187 | 194 | 196 | 229 | 229 | 188 | 188 | 173 | 173 | 194 | 194 | 212 | 212 | 195 | 195 | 223 | 223 | 188 | 188 |
| FCNI021504 | 101 | 101 | -1 | -1 | 203 | 203 | 157 | 157 | 187 | 187 | -1 | -1 | 229 | 229 | 188 | 188 | 173 | 173 | 194 | 194 | 212 | 212 | 195 | 207 | 223 | 223 | 188 | 188 |
| FCNI021505 | 95 | 95 | 185 | 185 | 203 | 203 | -1 | -1 | 187 | 187 | -1 | -1 | 229 | 229 | 188 | 188 | 173 | 173 | 194 | 194 | -1 | -1 | 195 | 195 | 223 | 223 | 188 | 207 |
| FCNI021481 | 101 | 101 | 177 | 177 | 203 | 203 | 157 | 157 | 187 | 187 | 194 | 196 | 229 | 229 | 188 | 188 | 173 | 173 | 194 | 194 | 212 | 212 | 207 | 207 | 223 | 223 | 188 | 207 |
| FCNI021486 | 101 | 101 | 177 | 177 | 203 | 203 | 157 | 157 | 187 | 187 | 196 | 196 | 229 | 229 | 188 | 188 | 173 | 173 | 194 | 194 | 212 | 212 | 207 | 207 | 223 | 223 | 188 | 207 |
| SADH2 | 101 | 101 | 177 | 177 | 203 | 203 | 157 | 157 | 187 | 187 | 196 | 196 | 229 | 229 | 188 | 188 | 173 | 173 | 194 | 194 | 212 | 212 | 195 | 195 | 223 | 223 | 188 | 188 |
| SADHH1 | 101 | 101 | 185 | 185 | 203 | 203 | 157 | 157 | 185 | 187 | -1 | -1 | 229 | 229 | 188 | 188 | 173 | 173 | 194 | 194 | 212 | 212 | 205 | 205 | 223 | 223 | 188 | 188 |
| TTOMH10 | 101 | 101 | 177 | 185 | 203 | 203 | 157 | 157 | 187 | 187 | 194 | 196 | 229 | 229 | 188 | 188 | 173 | 173 | 194 | 194 | 212 | 212 | 195 | 219 | 223 | 223 | 188 | 207 |
| TKLTH11 | 101 | 101 | 177 | 177 | 203 | 203 | 157 | 157 | 187 | 187 | 196 | 196 | 229 | 229 | 188 | 188 | 173 | 173 | 194 | 194 | 212 | 212 | 195 | 195 | 223 | 223 | 188 | 207 |
| TH12 | 101 | 101 | 177 | 185 | 203 | 203 | 157 | 157 | 187 | 187 | 194 | 196 | 229 | 229 | 188 | 188 | 173 | 173 | 194 | 194 | 212 | 212 | 195 | 219 | 223 | 223 | 188 | 207 |
| TKLTH14 | 101 | 101 | 177 | 177 | 203 | 203 | 157 | 157 | 187 | 187 | 194 | 194 | 229 | 229 | 188 | 188 | 173 | 173 | 194 | 194 | 212 | 212 | 195 | 195 | 223 | 223 | 188 | 207 |
| TADBH15 | 101 | 101 | 185 | 185 | -1 | -1 | 157 | 157 | 187 | 187 | 194 | 196 | 229 | 229 | 188 | 188 | -1 | -1 | 194 | 194 | 212 | 212 | 195 | 195 | 223 | 223 | 188 | 188 |
| TLMWH1 | 95 | 95 | 177 | 177 | 203 | 203 | -1 | -1 | -1 | -1 | 192 | 192 | -1 | -1 | 188 | 188 | 173 | 173 | 194 | 194 | 212 | 212 | 207 | 207 | 223 | 223 | 188 | 188 |
| TLMWH2 | 95 | 95 | 177 | 177 | 203 | 203 | -1 | -1 | 183 | 187 | 196 | 196 | -1 | -1 | 188 | 188 | 173 | 173 | 194 | 194 | 212 | 212 | -1 | -1 | 223 | 223 | 176 | 176 |
| TLMWH3 | 95 | 101 | 177 | 177 | 203 | 203 | 157 | 157 | 187 | 187 | 196 | 196 | 229 | 229 | 188 | 188 | 173 | 173 | 194 | 194 | 209 | 209 | 186 | 207 | -1 | -1 | 188 | 207 |
| THMNH13 | 101 | 101 | 185 | 185 | 203 | 203 | 157 | 157 | 187 | 187 | 194 | 196 | -1 | -1 | 188 | 188 | 173 | 173 | 194 | 194 | 212 | 212 | 195 | 219 | 223 | 223 | 188 | 207 |
| T44H16 | 95 | 101 | 177 | 177 | 203 | 203 | -1 | -1 | 187 | 187 | 196 | 196 | 229 | 229 | 188 | 188 | 173 | 173 | 194 | 194 | 209 | 212 | 195 | 195 | 223 | 223 | 196 | 196 |
| T43H17 | 101 | 101 | 177 | 177 | -1 | -1 | -1 | -1 | 185 | 187 | -1 | -1 | 229 | 229 | 188 | 188 | 173 | 173 | 194 | 194 | 212 | 212 | -1 | -1 | 223 | 223 | 188 | 188 |
| TH6 | 101 | 101 | 177 | 185 | 200 | 203 | 155 | 157 | 185 | 187 | 194 | 196 | 229 | 229 | 188 | 188 | 173 | 173 | 194 | 194 | 212 | 212 | 195 | 219 | 223 | 223 | 188 | 207 |
| TADBH4 | 101 | 101 | 177 | 177 | 203 | 203 | 157 | 157 | 185 | 187 | 194 | 194 | 229 | 229 | 188 | 188 | 173 | 173 | 194 | 194 | 212 | 212 | 219 | 219 | 223 | 223 | 188 | 188 |
| TPJW | 101 | 101 | 185 | 185 | 200 | 203 | 157 | 157 | 185 | 187 | 196 | 196 | 229 | 229 | 188 | 188 | 173 | 173 | 194 | 194 | 212 | 212 | 195 | 219 | 223 | 223 | 188 | 188 |
| TBHH7 | 101 | 101 | 181 | 181 | 203 | 203 | 157 | 157 | -1 | -1 | 194 | 194 | 229 | 229 | -1 | -1 | 173 | 173 | 194 | 194 | 212 | 212 | 193 | 193 | 223 | 223 | 188 | 188 |
| TKHH7 | 101 | 101 | 177 | 177 | 203 | 203 | 157 | 157 | 185 | 187 | 196 | 196 | 229 | 229 | 188 | 188 | -1 | -1 | 194 | 194 | -1 | -1 | 195 | 195 | 219 | 223 | 188 | 188 |
| TH9 | 101 | 101 | 177 | 177 | 203 | 203 | -1 | -1 | 185 | 187 | 194 | 194 | -1 | -1 | -1 | -1 | 173 | 173 | 194 | 194 | 212 | 212 | 195 | 195 | 223 | 223 | 188 | 188 |
| NZ1H1 | 101 | 101 | 185 | 185 | 203 | 203 | 157 | 157 | 187 | 187 | -1 | -1 | 229 | 229 | 188 | 188 | -1 | -1 | 194 | 194 | -1 | -1 | 195 | 195 | 223 | 223 | 188 | 207 |
| NZLH2 | 95 | 101 | 185 | 185 | 203 | 203 | -1 | -1 | -1 | -1 | 196 | 196 | 229 | 229 | 186 | 188 | -1 | -1 | 194 | 194 | 212 | 212 | 195 | 195 | 223 | 223 | 188 | 207 |
| NZLH3 | 95 | 95 | 177 | 177 | 203 | 203 | -1 | -1 | 187 | 187 | -1 | -1 | 229 | 229 | 188 | 188 | 173 | 173 | 194 | 194 | 212 | 212 | 195 | 195 | 223 | 223 | 188 | 207 |
| NZLH4 | 93 | 93 | 185 | 185 | 203 | 203 | -1 | -1 | 187 | 187 | 196 | 196 | 229 | 229 | 188 | 188 | 173 | 173 | 194 | 194 | 212 | 212 | 195 | 195 | 219 | 223 | 188 | 207 |
| NZLH5 | 95 | 95 | 177 | 185 | 203 | 203 | 157 | 157 | 187 | 187 | -1 | -1 | 229 | 229 | 188 | 188 | -1 | -1 | 194 | 194 | -1 | -1 | 195 | 195 | 223 | 223 | 188 | 188 |
| NZLH6 | 95 | 101 | 177 | 177 | 203 | 203 | 157 | 157 | 187 | 187 | -1 | -1 | 229 | 229 | 188 | 188 | 173 | 173 | 194 | 194 | 212 | 212 | 195 | 195 | 223 | 223 | 188 | 188 |
| NZLH7 | 95 | 95 | 177 | 177 | 203 | 203 | 157 | 157 | 187 | 187 | 194 | 194 | 229 | 229 | 188 | 188 | 173 | 173 | 194 | 194 | -1 | -1 | -1 | -1 | 219 | 223 | 188 | 188 |
| NZLH8 | 101 | 101 | 177 | 177 | 203 | 203 | 157 | 157 | 185 | 187 | 194 | 194 | 229 | 229 | 188 | 188 | 173 | 173 | 194 | 194 | -1 | -1 | -1 | -1 | 223 | 223 | 188 | 207 |
| NZLH9 | 101 | 101 | 177 | 177 | 203 | 203 | -1 | -1 | 185 | 187 | -1 | -1 | 229 | 229 | 188 | 188 | 173 | 173 | 194 | 194 | 212 | 212 | 195 | 195 | 223 | 223 | 188 | 207 |
| NZLH11 | 95 | 101 | 177 | 177 | 203 | 203 | 157 | 157 | 185 | 185 | 196 | 196 | 229 | 229 | 196 | 196 | 173 | 173 | 194 | 194 | 212 | 212 | -1 | -1 | 223 | 223 | 188 | 188 |
| NZLH10 | 101 | 101 | 177 | 177 | 203 | 203 | 157 | 157 | 179 | 195 | 194 | 196 | 229 | 229 | -1 | -1 | 173 | 173 | 194 | 194 | 209 | 209 | 195 | 195 | 223 | 223 | 188 | 188 |
| NZLH12 | 101 | 101 | 177 | 185 | 203 | 203 | 157 | 157 | 185 | 187 | 194 | 196 | 229 | 229 | 188 | 188 | 175 | 175 | 194 | 194 | 209 | 209 | -1 | -1 | 223 | 223 | 188 | 207 |
| NZLH13 | 101 | 101 | 177 | 177 | 203 | 203 | 157 | 157 | 185 | 187 | 196 | 196 | 229 | 229 | 188 | 188 | 173 | 173 | 194 | 194 | -1 | -1 | 219 | 219 | 223 | 223 | 188 | 188 |
| NZLH14 | 101 | 101 | 173 | 177 | 203 | 203 | -1 | -1 | 185 | 187 | 196 | 196 | 229 | 229 | 186 | 188 | 173 | 173 | 194 | 194 | 212 | 212 | 195 | 215 | 223 | 223 | 188 | 188 |
| NZLH15 | 101 | 101 | 177 | 177 | -1 | -1 | 157 | 157 | 179 | 179 | 194 | 196 | 229 | 229 | -1 | -1 | 173 | 173 | 194 | 194 | 212 | 212 | 193 | 195 | 223 | 223 | 188 | 188 |
| NZLH16 | 101 | 101 | 177 | 177 | 203 | 203 | 157 | 157 | 185 | 187 | 196 | 196 | 229 | 229 | 188 | 188 | 173 | 173 | 194 | 194 | -1 | -1 | 207 | 207 | 223 | 223 | 188 | 188 |
