## Supplementary Table S5 for "The influence of biosecurity on the diversity and management of a 120-year old joint insect-fungus pest invasion"

Supplementary Table S5. Alleles scored at each locus for strains of *A. areolatum*. ‘-1’ I indicate alleles that could not be scored and ‘Na’ fungal isolates that could not be deposited into the CMW collection as these have been isolated from the mycangia of ethanol-preserved female *S. noctilio*. The 11 microsatellite markers in this study were designed by (Mlonyeni et al. 2018). The forward primers of these loci were fluorescently labeled using FAM, NED, PET and VIC, dyes for filter set G5 (Applied biosystem). The fragments were analyzed on an ABI PRISM TM 3500xI analyser analyzer (Applied Biosystems), at the DNA sequencing facility, University of Pretoria. This collection included samples from Australia and New Newland (Samples IDs: **New South Wales**, N72 -N115; **Queensland**, Q1 – Q12; **South Australia**, S2 – S21; **Victoria**, V5 – V496; **The** **Australian commercial *Amylostereum* isolates**, Ecogrow RM1-Subbed 2013-KamFung and **Global isolates from CMW collection** , NZ01 – AT03).

| Sample ID | CMW number | Aa3-4 | | Aa3-5 | | Aa3-7 | | Aa3-8 | | Aa3-9 | | Aa3-14 | | Aa3-23 | | Aa3-24 | | Aa3-25 | | Aa4-3 | | Aa4-5 | |
| --- | --- | --- | --- | --- | --- | --- | --- | --- | --- | --- | --- | --- | --- | --- | --- | --- | --- | --- | --- | --- | --- | --- | --- |
| N72 | CMW49995 | 226 | 229 | 200 | 210 | 221 | 230 | 189 | 195 | 248 | 251 | 222 | 225 | 243 | 255 | 205 | 225 | 222 | 237 | 206 | 260 | 174 | 182 |
| N73 | CMW49996 | 226 | 229 | 200 | 210 | 221 | 230 | -1 | -1 | 248 | 251 | 222 | 225 | -1 | -1 | 205 | 225 | 222 | 237 | 206 | 260 | -1 | -1 |
| N74 | CMW53306 | 226 | 229 | 200 | 210 | 221 | 230 | 189 | 195 | 248 | 251 | 222 | 225 | 243 | 255 | 205 | 225 | 222 | 237 | 206 | 260 | 174 | 182 |
| N75 | CMW49997 | -1 | -1 | 200 | 210 | 221 | 230 | 189 | 195 | 248 | 251 | 222 | 225 | 243 | 243 | 205 | 225 | 222 | 237 | 206 | 260 | 174 | 182 |
| N76 | CMW49998 | -1 | -1 | -1 | -1 | 221 | 230 | 189 | 195 | 248 | 251 | 222 | 225 | -1 | -1 | 205 | 225 | 222 | 237 | 206 | 260 | 174 | 182 |
| N80 | CMW50245 | 226 | 229 | 200 | 210 | 221 | 230 | -1 | -1 | 248 | 251 | 222 | 225 | 243 | 255 | 205 | 225 | 222 | 237 | -1 | -1 | 174 | 182 |
| N95 | CMW53576 | -1 | -1 | 200 | 210 | 221 | 230 | 189 | 195 | 248 | 251 | 222 | 225 | 243 | 243 | 205 | 225 | 222 | 237 | 206 | 260 | 174 | 182 |
| N96 | CMW53577 | 226 | 229 | 200 | 210 | 221 | 230 | 189 | 195 | 248 | 251 | 222 | 225 | 243 | 243 | 205 | 225 | 222 | 237 | 206 | 260 | 174 | 182 |
| N97 | CMW53578 | 226 | 229 | 200 | 210 | 221 | 230 | 189 | 195 | 248 | 251 | 222 | 225 | 243 | 243 | 205 | 225 | 222 | 237 | 206 | 260 | 174 | 182 |
| N98 | CMW53495 | 226 | 229 | 200 | 210 | 221 | 230 | 189 | 195 | 248 | 251 | 222 | 225 | 243 | 243 | 205 | 225 | 222 | 237 | 206 | 260 | 174 | 182 |
| N99 | CMW53579 | 226 | 229 | 200 | 210 | 221 | 230 | 189 | 195 | 248 | 251 | 222 | 225 | 243 | 243 | 205 | 225 | 222 | 237 | 206 | 260 | 174 | 182 |
| N100 | CMW53496 | 226 | 229 | 200 | 210 | 221 | 230 | 189 | 195 | 248 | 251 | -1 | -1 | 243 | 243 | 205 | 225 | -1 | -1 | 206 | 260 | 174 | 182 |
| N101 | CMW53497 | 226 | 229 | 200 | 210 | 221 | 230 | 189 | 195 | 248 | 251 | -1 | -1 | 243 | 243 | 205 | 225 | -1 | -1 | 206 | 260 | 174 | 182 |
| N102 | CMW53580 | 226 | 229 | 200 | 210 | 221 | 230 | 189 | 195 | 248 | 251 | 222 | 225 | 243 | 243 | 205 | 225 | 222 | 237 | 206 | 260 | 174 | 182 |
| N103 | CMW53498 | 226 | 229 | 200 | 210 | 221 | 230 | 189 | 195 | 248 | 251 | 222 | 225 | 243 | 243 | 205 | 225 | 222 | 237 | 206 | 260 | 174 | 182 |
| N104 | CMW53499 | 226 | 229 | 200 | 210 | 221 | 230 | 189 | 195 | 248 | 251 | 222 | 225 | 243 | 243 | 205 | 225 | 222 | 237 | 206 | 260 | 174 | 182 |
| N105 | CMW53500 | 226 | 229 | 200 | 210 | 221 | 230 | 189 | 195 | 248 | 251 | 222 | 225 | 243 | 243 | 205 | 225 | 222 | 237 | 206 | 260 | 174 | 182 |
| N107 | CMW53329 | 226 | 229 | 200 | 210 | 221 | 230 | 189 | 195 | 248 | 251 | 222 | 225 | 243 | 243 | 205 | 225 | 222 | 237 | 206 | 260 | 174 | 182 |
| N108 | CMW53330 | 226 | 229 | 200 | 210 | 221 | 230 | 189 | 195 | 248 | 251 | 222 | 225 | 243 | 243 | 205 | 225 | 222 | 237 | 206 | 260 | 174 | 182 |
| N109 | CMW53501 | 226 | 229 | 200 | 210 | 221 | 230 | 189 | 195 | 248 | 251 | 222 | 225 | 243 | 243 | 205 | 225 | 222 | 237 | 206 | 260 | 174 | 182 |
| N110 | CMW53502 | 226 | 229 | 200 | 210 | 221 | 230 | 189 | 195 | 248 | 251 | 222 | 225 | 243 | 243 | 205 | 225 | 222 | 237 | 206 | 260 | 174 | 182 |
| N111 | CMW53331 | 226 | 229 | 200 | 210 | 221 | 230 | 189 | 195 | 248 | 251 | 222 | 225 | 243 | 243 | 205 | 225 | 222 | 237 | 206 | 260 | 174 | 182 |
| N112 | CMW53581 | 226 | 229 | 200 | 210 | 221 | 230 | 189 | 195 | 248 | 251 | 222 | 225 | 243 | 243 | 205 | 225 | 222 | 237 | 206 | 260 | 174 | 182 |
| N114 | CMW53333 | 226 | 229 | 200 | 210 | 221 | 230 | 189 | 195 | 248 | 251 | 222 | 225 | 243 | 243 | 205 | 225 | 222 | 237 | 206 | 260 | 174 | 182 |
| N115 | CMW53334 | 226 | 229 | 200 | 210 | 221 | 230 | 189 | 195 | 248 | 251 | 222 | 225 | 243 | 243 | 205 | 225 | 222 | 237 | 206 | 260 | 174 | 182 |
| Q1 | CMW49666 | -1 | -1 | 200 | 210 | 221 | 230 | 189 | 195 | 248 | 251 | 222 | 225 | 243 | 243 | 205 | 225 | 222 | 237 | 206 | 260 | 174 | 182 |
| Q2 | CMW49667 | 226 | 229 | 200 | 210 | 221 | 230 | 189 | 195 | 248 | 251 | 222 | 225 | 243 | 243 | 205 | 225 | 222 | 237 | 206 | 260 | 174 | 182 |
| Q3 | CMW49668 | 226 | 229 | 200 | 210 | 221 | 230 | 189 | 195 | 248 | 251 | 222 | 225 | 243 | 243 | 205 | 225 | 222 | 237 | -1 | -1 | 174 | 182 |
| Q6 | CMW53582 | 226 | 229 | 200 | 210 | 221 | 230 | 189 | 195 | 248 | 251 | 222 | 225 | 243 | 243 | 205 | 225 | 222 | 237 | 206 | 260 | 174 | 182 |
| Q7 | CMW53583 | 226 | 229 | 200 | 210 | 221 | 230 | 189 | 195 | 248 | 251 | 222 | 225 | 243 | 243 | 205 | 225 | 222 | 237 | 206 | 260 | 174 | 182 |
| Q8 | CMW53584 | -1 | -1 | 200 | 210 | 221 | 230 | 189 | 195 | 248 | 251 | 222 | 225 | 243 | 243 | 205 | 225 | 222 | 237 | 206 | 260 | 174 | 182 |
| Q9 | CMW53585 | 226 | 229 | 200 | 210 | -1 | -1 | 189 | 195 | 248 | 251 | 222 | 225 | 243 | 243 | 205 | 225 | 222 | 237 | 206 | 260 | 174 | 182 |
| Q10 | CMW53337 | 226 | 229 | 200 | 210 | 221 | 230 | 189 | 195 | 248 | 251 | 222 | 225 | 243 | 243 | 205 | 225 | 222 | 237 | 206 | 260 | 174 | 182 |
| Q11 | CMW53338 | 226 | 229 | 200 | 210 | 221 | 230 | 189 | 195 | 248 | 251 | 222 | 225 | 243 | 243 | 205 | 225 | 222 | 237 | 206 | 260 | 174 | 182 |
| Q12 | CMW53339 | 226 | 229 | 200 | 210 | 221 | 230 | 189 | 195 | 248 | 251 | 222 | 225 | 243 | 243 | 205 | 225 | 222 | 237 | 206 | 260 | 174 | 182 |
| S2 | CMW50006 | 226 | 229 | 200 | 210 | 221 | 230 | 189 | 195 | 248 | 251 | 222 | 225 | 243 | 243 | 205 | 225 | 222 | 237 | 206 | 260 | 174 | 182 |
| S12 | CMW53586 | 226 | 229 | 200 | 210 | 221 | 230 | 189 | 195 | 248 | 251 | 222 | 225 | 243 | 243 | 205 | 225 | 222 | 237 | 206 | 260 | 174 | 182 |
| S13 | CMW53503 | 226 | 229 | 200 | 210 | 221 | 230 | 189 | 195 | 248 | 251 | 222 | 225 | 243 | 243 | 205 | 225 | 222 | 237 | 206 | 260 | 174 | 182 |
| S14 | CMW53504 | 226 | 229 | 200 | 210 | 221 | 230 | 189 | 195 | 248 | 251 | 222 | 225 | 243 | 243 | 205 | 225 | 222 | 237 | 206 | 260 | 174 | 182 |
| S15 | CMW53505 | 226 | 229 | 200 | 210 | 221 | 230 | 189 | 195 | 248 | 251 | 222 | 225 | 243 | 243 | 205 | 225 | 222 | 237 | 206 | 260 | 174 | 182 |
| S16 | CMW53587 | 226 | 229 | 200 | 210 | 221 | 230 | 189 | 195 | 248 | 251 | 222 | 225 | 243 | 243 | -1 | -1 | 222 | 237 | 206 | 260 | 174 | 182 |
| S17 | CMW53506 | 226 | 229 | 200 | 210 | 221 | 230 | 189 | 195 | 248 | 251 | -1 | -1 | 243 | 243 | -1 | -1 | -1 | -1 | 206 | 260 | 174 | 182 |
| S18 | CMW53588 | 226 | 229 | 200 | 210 | 221 | 230 | 189 | 195 | 248 | 251 | 222 | 225 | 243 | 243 | 205 | 225 | 222 | 237 | 206 | 260 | 174 | 182 |
| S20 | CMW53507 | 226 | 229 | 200 | 210 | 221 | 230 | 189 | 195 | 248 | 251 | 222 | 225 | 243 | 243 | 205 | 225 | 222 | 237 | 206 | 260 | 174 | 182 |
| S21 | CMW53336 | 226 | 229 | 200 | 210 | 221 | 230 | 189 | 195 | 248 | 251 | 222 | 225 | 243 | 243 | 205 | 225 | 222 | 237 | 206 | 260 | 174 | 182 |
| V5 | CMW53293 | -1 | -1 | 200 | 210 | 221 | 230 | 189 | 195 | 248 | 251 | 222 | 225 | 243 | 243 | 205 | 225 | 222 | 237 | 206 | 260 | 174 | 182 |
| V10 | CMW49657 | 226 | 229 | 200 | 210 | 221 | 230 | 189 | 195 | 248 | 251 | 222 | 225 | 243 | 243 | 205 | 225 | 222 | 237 | 206 | 260 | 174 | 182 |
| V11 | CMW49658 | 226 | 229 | 200 | 210 | 221 | 230 | 189 | 195 | 248 | 251 | 222 | 225 | -1 | -1 | 205 | 225 | 222 | 237 | -1 | -1 | 174 | 182 |
| V12 | CMW49659 | 226 | 229 | 200 | 210 | 221 | 230 | 189 | 195 | 248 | 251 | 222 | 225 | 243 | 243 | 205 | 225 | 222 | 237 | 206 | 260 | 174 | 182 |
| V15 | CMW49660 | 226 | 229 | 200 | 210 | 221 | 230 | 189 | 195 | 248 | 251 | 222 | 225 | -1 | -1 | 205 | 225 | 222 | 237 | -1 | -1 | -1 | -1 |
| V40 | CMW49661 | 226 | 229 | 200 | 210 | 221 | 230 | 189 | 195 | 248 | 251 | 222 | 225 | 243 | 243 | 205 | 225 | 222 | 237 | 206 | 260 | 174 | 182 |
| V100 | CMW49663 | 226 | 229 | 200 | 210 | 221 | 230 | 189 | 195 | 248 | 251 | -1 | -1 | 243 | 243 | 205 | 225 | 222 | 237 | 206 | 260 | 174 | 182 |
| V106 | CMW49792 | 226 | 229 | 200 | 210 | 221 | 230 | 189 | 195 | 248 | 251 | -1 | -1 | 243 | 243 | 205 | 225 | 222 | 237 | 206 | 260 | -1 | -1 |
| V111 | CMW49664 | 226 | 229 | 200 | 210 | 221 | 230 | 189 | 195 | 248 | 251 | 222 | 225 | -1 | -1 | 205 | 225 | 222 | 237 | -1 | -1 | -1 | -1 |
| V351 | CMW50008 | 226 | 229 | 200 | 210 | 221 | 230 | 189 | 195 | 248 | 251 | 222 | 225 | 243 | 243 | 205 | 225 | 222 | 237 | 206 | 260 | 174 | 182 |
| V353 | CMW50009 | 226 | 229 | 200 | 210 | -1 | -1 | 189 | 195 | 248 | 251 | 222 | 225 | 243 | 243 | 205 | 225 | 222 | 237 | 206 | 260 | 174 | 182 |
| V354 | CMW50002 | -1 | -1 | -1 | -1 | 221 | 230 | 189 | 195 | 248 | 251 | 222 | 225 | -1 | -1 | 205 | 225 | 222 | 237 | 206 | 260 | 174 | 182 |
| V357 | CMW53307 | 226 | 229 | 200 | 210 | 221 | 230 | -1 | -1 | -1 | -1 | 222 | 225 | 243 | 243 | 205 | 225 | 222 | 237 | -1 | -1 | 174 | 182 |
| V358 | CMW53296 | 226 | 229 | 200 | 210 | 221 | 230 | 189 | 195 | 248 | 251 | 222 | 225 | 243 | 243 | 205 | 225 | 222 | 237 | 206 | 260 | 174 | 182 |
| V361 | CMW53311 | 226 | 229 | 200 | 210 | 221 | 230 | 189 | 195 | 248 | 251 | 222 | 225 | 243 | 243 | 205 | 225 | 222 | 237 | 206 | 260 | 174 | 182 |
| V372 | CMW53297 | 226 | 229 | 200 | 210 | 221 | 230 | 189 | 195 | 248 | 251 | 222 | 225 | 243 | 243 | 205 | 225 | 222 | 237 | 206 | 260 | 174 | 182 |
| V377 | CMW53308 | 226 | 229 | 200 | 210 | 221 | 230 | -1 | -1 | 248 | 251 | 222 | 225 | 243 | 243 | -1 | -1 | 222 | 237 | -1 | -1 | 174 | 182 |
| V378 | CMW53309 | 226 | 229 | 200 | 210 | 221 | 230 | -1 | -1 | 248 | 251 | 222 | 225 | 243 | 255 | 205 | 225 | 222 | 237 | 206 | 260 | 174 | 182 |
| V385 | CMW53315 | -1 | -1 | 200 | 210 | 221 | 230 | -1 | -1 | 248 | 251 | 222 | 225 | -1 | -1 | 205 | 225 | 222 | 237 | -1 | -1 | 174 | 182 |
| V396 | CMW53314 | 226 | 229 | 200 | 210 | 221 | 230 | -1 | -1 | 248 | 251 | 222 | 225 | 243 | 255 | 205 | 225 | 222 | 237 | -1 | -1 | -1 | -1 |
| V404 | CMW53590 | 226 | 229 | 200 | 210 | 221 | 230 | 189 | 195 | 248 | 251 | 222 | 225 | 243 | 243 | 205 | 225 | 222 | 237 | 206 | 260 | 174 | 182 |
| V406 | CMW53591 | 226 | 229 | 200 | 210 | 221 | 230 | 189 | 195 | 248 | 251 | 222 | 225 | 243 | 243 | 205 | 225 | 222 | 237 | 206 | 260 | 174 | 182 |
| V414 | CMW53592 | 226 | 229 | 200 | 210 | -1 | -1 | 189 | 195 | 248 | 251 | 222 | 225 | 243 | 243 | 205 | 225 | 222 | 237 | 206 | 260 | 174 | 182 |
| V420 | CMW53593 | 226 | 229 | 200 | 210 | 221 | 230 | 189 | 195 | 248 | 251 | 222 | 225 | 243 | 243 | 205 | 225 | 222 | 237 | 206 | 260 | 174 | 182 |
| V421 | CMW53594 | 226 | 229 | 200 | 210 | 221 | 230 | 189 | 195 | 248 | 251 | 222 | 225 | 243 | 243 | 205 | 225 | 222 | 237 | 206 | 260 | 174 | 182 |
| V425 | CMW53312 | 226 | 229 | 200 | 210 | 221 | 230 | 189 | 195 | -1 | -1 | -1 | -1 | -1 | -1 | -1 | -1 | 222 | 237 | 206 | 260 | 174 | 182 |
| V436 | CMW53300 | 226 | 229 | 200 | 210 | 221 | 230 | 189 | 195 | 248 | 251 | 222 | 225 | 243 | 243 | 205 | 225 | 222 | 237 | 206 | 260 | 174 | 182 |
| V437 | CMW53301 | 226 | 229 | 200 | 210 | 221 | 230 | 189 | 195 | 248 | 251 | 222 | 225 | 243 | 243 | 205 | 225 | 222 | 237 | 206 | 260 | 174 | 182 |
| V438 | CMW53302 | 226 | 229 | 200 | 210 | 221 | 230 | 189 | 195 | 248 | 251 | 222 | 225 | 243 | 255 | 205 | 225 | 222 | 237 | 206 | 260 | 174 | 182 |
| V439 | CMW53303 | 226 | 229 | 200 | 210 | 221 | 230 | 189 | 195 | 248 | 251 | 222 | 225 | 243 | 243 | 205 | 225 | 222 | 237 | 206 | 260 | 174 | 182 |
| V445 | CMW53305 | 226 | 229 | 200 | 210 | 221 | 230 | 189 | 195 | 248 | 251 | 222 | 225 | -1 | -1 | 205 | 225 | 222 | 237 | -1 | -1 | -1 | -1 |
| V448 | CMW53371 | 226 | 229 | 200 | 210 | 221 | 230 | 189 | 195 | 248 | 251 | 222 | 225 | 243 | 243 | 205 | 225 | 222 | 237 | 206 | 260 | 174 | 182 |
| V449 | CMW53508 | 226 | 229 | 200 | 210 | 221 | 230 | 189 | 195 | 248 | 251 | 222 | 225 | 243 | 243 | 205 | 225 | 222 | 237 | 206 | 260 | 174 | 182 |
| V451 | CMW53596 | 226 | 229 | 200 | 210 | -1 | -1 | 189 | 195 | 248 | 251 | 222 | 225 | 243 | 243 | 205 | 225 | 222 | 237 | 206 | 260 | 174 | 182 |
| V455 | CMW53510 | 226 | 229 | 200 | 210 | -1 | -1 | 189 | 195 | 248 | 251 | 222 | 225 | 243 | 243 | 205 | 225 | 222 | 237 | 206 | 260 | 174 | 182 |
| V456 | CMW53511 | 226 | 229 | 200 | 210 | 221 | 230 | 189 | 195 | 248 | 251 | 222 | 225 | 243 | 243 | 205 | 225 | 222 | 237 | 206 | 260 | 174 | 182 |
| V458 | CMW53513 | 226 | 229 | 200 | 210 | 221 | 230 | 189 | 195 | 248 | 251 | 222 | 225 | 243 | 243 | 205 | 225 | 222 | 237 | 206 | 260 | 174 | 182 |
| V459 | CMW53598 | 226 | 229 | 200 | 210 | 221 | 230 | 189 | 195 | 248 | 251 | 222 | 225 | 243 | 243 | 205 | 225 | 222 | 237 | 206 | 260 | -1 | -1 |
| V460 | CMW53599 | 226 | 229 | 200 | 210 | 221 | 230 | 189 | 195 | 248 | 251 | 222 | 225 | 243 | 243 | 205 | 225 | 222 | 237 | 206 | 260 | 174 | 182 |
| V461 | CMW53514 | 226 | 229 | 200 | 210 | 221 | 230 | 189 | 195 | 248 | 251 | 222 | 225 | 243 | 243 | 205 | 225 | 222 | 237 | 206 | 260 | 174 | 182 |
| V462 | CMW53515 | 226 | 229 | 200 | 210 | 221 | 230 | 189 | 195 | 248 | 251 | 222 | 225 | 243 | 243 | 205 | 225 | 222 | 237 | 206 | 260 | 174 | 182 |
| V463 | CMW53516 | 226 | 229 | 200 | 210 | 221 | 230 | 189 | 195 | 248 | 251 | 222 | 225 | 243 | 243 | 205 | 225 | 222 | 237 | 206 | 260 | 174 | 182 |
| V464 | CMW53600 | 226 | 229 | 200 | 210 | 221 | 230 | 189 | 195 | 248 | 251 | 222 | 225 | 243 | 243 | 205 | 225 | 222 | 237 | 206 | 260 | 174 | 182 |
| V465 | CMW53517 | 226 | 229 | 200 | 210 | 221 | 230 | 189 | 195 | 248 | 251 | 222 | 225 | 243 | 243 | 205 | 225 | 222 | 237 | 206 | 260 | -1 | -1 |
| V466 | CMW53518 | 226 | 229 | 200 | 210 | 221 | 230 | 189 | 195 | 248 | 251 | 222 | 225 | 243 | 243 | 205 | 225 | 222 | 237 | 206 | 260 | 174 | 182 |
| V467 | CMW53519 | 226 | 229 | 200 | 210 | 221 | 230 | 189 | 195 | 248 | 251 | 222 | 225 | 243 | 243 | 205 | 225 | 222 | 237 | 206 | 260 | 174 | 182 |
| V468 | CMW53520 | 226 | 229 | 200 | 210 | 221 | 230 | 189 | 195 | 248 | 251 | 222 | 225 | 243 | 243 | 205 | 225 | 222 | 237 | 206 | 260 | 174 | 182 |
| V469 | CMW53521 | 226 | 229 | 200 | 210 | 221 | 230 | 189 | 195 | 248 | 251 | 222 | 225 | 243 | 243 | 205 | 225 | 222 | 237 | 206 | 260 | 174 | 182 |
| V470 | CMW53522 | 226 | 229 | 200 | 210 | 221 | 230 | 189 | 195 | 248 | 251 | 222 | 225 | 243 | 243 | 205 | 225 | 222 | 237 | 206 | 260 | 174 | 182 |
| V471 | CMW53523 | 226 | 229 | 200 | 210 | 221 | 230 | 189 | 195 | 248 | 251 | 222 | 225 | 243 | 243 | 205 | 225 | 222 | 237 | 206 | 260 | 174 | 182 |
| V472 | CMW53601 | 226 | 229 | 200 | 210 | 221 | 230 | 189 | 195 | 248 | 251 | 222 | 225 | 243 | 243 | 205 | 225 | 222 | 237 | 206 | 260 | 174 | 182 |
| V474 | CMW53602 | 226 | 229 | 200 | 210 | 221 | 230 | 189 | 195 | 248 | 251 | 222 | 225 | 243 | 243 | 205 | 225 | 222 | 237 | 206 | 260 | 174 | 182 |
| V475 | CMW53603 | 226 | 229 | 200 | 210 | 221 | 230 | 189 | 195 | 248 | 251 | 222 | 225 | 243 | 243 | 205 | 225 | 222 | 237 | 206 | 260 | 174 | 182 |
| V476 | CMW53525 | 226 | 229 | 200 | 210 | 221 | 230 | 189 | 195 | 248 | 251 | 222 | 225 | 243 | 255 | 205 | 225 | 222 | 237 | 206 | 260 | 174 | 182 |
| V477 | CMW53604 | 226 | 229 | 200 | 210 | 221 | 230 | 189 | 195 | 248 | 251 | 222 | 225 | 243 | 243 | 205 | 225 | 222 | 237 | 206 | 260 | 174 | 182 |
| V478 | CMW53526 | 226 | 229 | 200 | 210 | 221 | 230 | 189 | 195 | 248 | 251 | 222 | 225 | 243 | 243 | 205 | 225 | 222 | 237 | 206 | 260 | 174 | 182 |
| V479 | CMW53605 | 226 | 229 | 200 | 210 | 221 | 230 | 189 | 195 | 248 | 251 | 222 | 225 | 243 | 243 | 205 | 225 | 222 | 237 | 206 | 260 | 174 | 182 |
| V481 | CMW53607 | 226 | 229 | 200 | 210 | 221 | 230 | 189 | 195 | 248 | 251 | 222 | 225 | 243 | 243 | 205 | 225 | 222 | 237 | 206 | 260 | 174 | 182 |
| V482 | CMW53527 | 226 | 229 | 200 | 210 | 221 | 230 | 189 | 195 | 248 | 251 | 222 | 225 | 243 | 243 | 205 | 225 | 222 | 237 | 206 | 260 | 174 | 182 |
| V483 | CMW53528 | 226 | 229 | 200 | 210 | 221 | 230 | 189 | 195 | 248 | 251 | 222 | 225 | 243 | 243 | 205 | 225 | 222 | 237 | 206 | 260 | 174 | 182 |
| V484 | CMW53529 | 226 | 229 | 200 | 210 | 221 | 230 | 189 | 195 | 248 | 251 | 222 | 225 | 243 | 243 | 205 | 225 | 222 | 237 | 206 | 260 | 174 | 182 |
| V485 | CMW53608 | 226 | 229 | 200 | 210 | 221 | 230 | 189 | 195 | 248 | 251 | 222 | 225 | 243 | 243 | 205 | 225 | 222 | 237 | 206 | 260 | 174 | 182 |
| V486 | CMW53609 | 226 | 229 | 200 | 210 | 221 | 230 | 189 | 195 | 248 | 251 | 222 | 225 | 243 | 243 | 205 | 225 | 222 | 237 | 206 | 260 | 174 | 182 |
| V487 | CMW53530 | 226 | 229 | 200 | 210 | 221 | 230 | 189 | 195 | 248 | 251 | 222 | 225 | 243 | 243 | 205 | 225 | 222 | 237 | 206 | 260 | 174 | 182 |
| V488 | CMW53610 | 226 | 229 | 200 | 210 | 221 | 230 | 189 | 195 | 248 | 251 | 222 | 225 | 243 | 243 | 205 | 225 | 222 | 237 | 206 | 260 | 174 | 182 |
| V489 | CMW53531 | 226 | 229 | 200 | 210 | 221 | 230 | 189 | 195 | 248 | 251 | 222 | 225 | 243 | 243 | 205 | 225 | 222 | 237 | 206 | 260 | 174 | 182 |
| V490 | CMW53532 | 226 | 229 | 200 | 210 | 221 | 230 | 189 | 195 | 248 | 251 | 222 | 225 | 243 | 243 | 205 | 225 | 222 | 237 | 206 | 260 | 174 | 182 |
| V491 | CMW53533 | 226 | 229 | 200 | 210 | 221 | 230 | 189 | 195 | 248 | 251 | 222 | 225 | 243 | 243 | 205 | 225 | 222 | 237 | 206 | 260 | -1 | -1 |
| V492 | CMW53611 | 226 | 229 | 200 | 210 | 221 | 230 | 189 | 195 | 248 | 251 | 222 | 225 | 243 | 243 | 205 | 225 | 222 | 237 | 206 | 260 | 174 | 182 |
| V493 | CMW53612 | 226 | 229 | 200 | 210 | 221 | 230 | 189 | 195 | 248 | 251 | 222 | 225 | 243 | 243 | 205 | 225 | 222 | 237 | 206 | 260 | 174 | 182 |
| V494 | CMW53534 | 226 | 229 | 200 | 210 | 221 | 230 | 189 | 195 | 248 | 251 | 222 | 225 | 243 | 243 | 205 | 225 | 222 | 237 | 206 | 260 | 174 | 182 |
| V495 | CMW53613 | 226 | 229 | 200 | 210 | 221 | 230 | 189 | 195 | 248 | 251 | 222 | 225 | 243 | 243 | 205 | 225 | 222 | 237 | 206 | 260 | 174 | 182 |
| V496 | CMW53614 | 226 | 229 | 200 | 210 | 221 | 230 | 189 | 195 | 248 | 251 | 222 | 225 | 243 | 243 | 205 | 225 | 222 | 237 | 206 | 260 | 174 | 182 |
| EcogrowRM1 | CMW53288 | 226 | 229 | 200 | 210 | 221 | 230 | 189 | 195 | 248 | 251 | 222 | 225 | 243 | 243 | 205 | 225 | 222 | 237 | 206 | 260 | 174 | 182 |
| Subbed2013 | CMW53289 | 226 | 229 | 200 | 210 | 221 | 230 | 189 | 195 | 248 | 251 | 222 | 225 | 243 | 243 | 205 | 225 | 222 | 237 | 206 | 260 | 174 | 182 |
| KamFung | CMW49355 | 226 | 229 | 200 | 210 | 221 | 230 | 189 | 195 | 248 | 251 | 222 | 225 | 243 | 243 | 205 | 225 | 222 | 237 | 206 | 260 | 174 | 182 |
| NZ1 | Na | 226 | 229 | 200 | 210 | 221 | 230 | 189 | 195 | 248 | 251 | 222 | 225 | -1 | -1 | 205 | 225 | 222 | 237 | 206 | 260 | -1 | -1 |
| NZ2 | Na | 226 | 229 | 200 | 210 | -1 | -1 | 189 | 195 | 248 | 251 | 222 | 225 | 243 | 243 | 205 | 225 | 222 | 237 | 206 | 260 | 174 | 182 |
| NZ6 | Na | 226 | 229 | 200 | 210 | 221 | 230 | 189 | 195 | 248 | 251 | 222 | 225 | 243 | 243 | 205 | 225 | 222 | 237 | 206 | 260 | -1 | -1 |
| NZ7 | Na | 226 | 229 | 200 | 210 | 221 | 230 | 189 | 195 | 248 | 251 | 222 | 225 | 243 | 243 | 205 | 225 | 222 | 237 | 206 | 260 | 174 | 182 |
| NZs3a | Na | 226 | 229 | 200 | 210 | 221 | 230 | 189 | 195 | 248 | 248 | 225 | 225 | -1 | -1 | 205 | 225 | 222 | 237 | 206 | 260 | 174 | 182 |
| NZs3b | Na | 226 | 229 | 200 | 210 | 221 | 230 | 189 | 195 | 248 | 251 | 222 | 225 | 243 | 243 | 205 | 225 | 222 | 237 | 206 | 260 | 174 | 182 |
| NZ3c | Na | 226 | 229 | 200 | 210 | 221 | 230 | 189 | 195 | 248 | 251 | 222 | 225 | 243 | 243 | 205 | 225 | 222 | 237 | 206 | 260 | 174 | 182 |
| NZ3d | Na | 226 | 229 | 200 | 210 | -1 | -1 | 189 | 195 | 248 | 251 | 222 | 225 | -1 | -1 | 205 | 225 | 222 | 237 | -1 | -1 | 174 | 182 |
| NZ3e | Na | 226 | 229 | 200 | 210 | -1 | -1 | 189 | 195 | 248 | 251 | 222 | 225 | 243 | 243 | 205 | 225 | 222 | 237 | -1 | -1 | 174 | 182 |
| NZ4a | Na | 226 | 229 | 200 | 210 | 221 | 230 | 189 | 195 | 248 | 251 | 222 | 225 | 243 | 243 | 205 | 225 | 222 | 237 | 206 | 260 | 174 | 182 |
| NZ4c | Na | 226 | 229 | 200 | 210 | 221 | 230 | 189 | 195 | 248 | 251 | 222 | 225 | 243 | 243 | 205 | 225 | 222 | 237 | 206 | 260 | 174 | 182 |
| NZ4d | Na | 226 | 229 | 200 | 210 | 221 | 230 | 189 | 195 | 248 | 251 | 222 | 225 | 243 | 243 | 205 | 225 | 222 | 237 | 206 | 260 | 174 | 182 |
| NZS9 | CMW53317 | 226 | 229 | 200 | 210 | 221 | 221 | 189 | 195 | 248 | 251 | 222 | 225 | -1 | -1 | 205 | 225 | -1 | -1 | 206 | 260 | 174 | 182 |
| NZ9C | CMW53320 | -1 | -1 | 200 | 210 | 221 | 230 | 189 | 195 | 248 | 251 | 222 | 225 | 243 | 255 | -1 | -1 | 222 | 237 | 206 | 260 | 174 | 182 |
| NZ9E | CMW53322 | -1 | -1 | 210 | 210 | 221 | 230 | 189 | 195 | 248 | 251 | 222 | 225 | 243 | 255 | -1 | -1 | 222 | 237 | 206 | 260 | 174 | 182 |
| NZ9H | CMW53325 | -1 | -1 | 200 | 210 | 221 | 230 | -1 | -1 | 248 | 251 | 222 | 225 | -1 | -1 | 205 | 225 | 222 | 237 | 206 | 260 | 174 | 182 |
| NZ12a | Na | 226 | 229 | 200 | 210 | -1 | -1 | 189 | 195 | 248 | 251 | 222 | 225 | 243 | 243 | 205 | 225 | 222 | 237 | -1 | -1 | 174 | 182 |
| NZ12b | Na | 226 | 229 | 200 | 210 | 221 | 230 | 189 | 195 | 248 | 251 | 222 | 225 | 243 | 243 | 206 | 226 | 222 | 237 | 206 | 260 | 174 | 182 |
| NZ14a | Na | 226 | 229 | 200 | 210 | 221 | 230 | 189 | 195 | 248 | 251 | 222 | 225 | -1 | -1 | 205 | 225 | 222 | 237 | 206 | 260 | -1 | -1 |
| NZ14c | Na | 226 | 229 | 200 | 210 | 221 | 230 | 189 | 195 | 248 | 251 | 222 | 225 | 243 | 243 | 205 | 225 | 222 | 237 | 206 | 260 | 174 | 182 |
| NZ14d | Na | 226 | 229 | 200 | 210 | -1 | -1 | 189 | 195 | 248 | 251 | 222 | 225 | -1 | -1 | 205 | 225 | 222 | 237 | 206 | 260 | 174 | 182 |
| NZ14e | Na | 226 | 229 | 200 | 210 | 221 | 230 | 189 | 195 | 248 | 251 | 222 | 225 | 243 | 255 | 205 | 225 | 222 | 237 | 206 | 260 | 174 | 182 |
| NZ14f | Na | 226 | 229 | 200 | 210 | 221 | 230 | 189 | 195 | 248 | 251 | 222 | 225 | 243 | 243 | 206 | 226 | 222 | 237 | 206 | 260 | 174 | 182 |
| NZ14g | Na | 226 | 229 | 200 | 210 | 221 | 230 | 189 | 195 | 248 | 251 | 222 | 225 | 243 | 243 | 206 | 226 | 222 | 237 | 206 | 260 | 174 | 182 |
| NZ14h | Na | 226 | 229 | 200 | 210 | 221 | 230 | 189 | 195 | 248 | 251 | 225 | 225 | 243 | 243 | 205 | 225 | 222 | 237 | -1 | -1 | 174 | 182 |
| NZ01 | CMW37117 | -1 | -1 | 200 | 210 | 221 | 221 | 189 | 195 | 248 | 251 | 222 | 225 | -1 | -1 | 205 | 225 | -1 | -1 | 206 | 260 | 174 | 182 |
| AUS01 | CMW4644 | -1 | -1 | 197 | 210 | 221 | 221 | 189 | 189 | 248 | 251 | 222 | 222 | -1 | -1 | 205 | 205 | -1 | -1 | 206 | 206 | 174 | 186 |
| SA01 | CMW13659 | 226 | 229 | 200 | 210 | 221 | 221 | 189 | 195 | 248 | 251 | 222 | 225 | 253 | 253 | 205 | 333 | -1 | -1 | 206 | 260 | 174 | 186 |
| CN01 | CMW37118 | 226 | 226 | 200 | 210 | 221 | 221 | 189 | 195 | 251 | 251 | 225 | 225 | 246 | 246 | 205 | 225 | -1 | -1 | 206 | 260 | 174 | 182 |
| CN02 | CMW37119 | 226 | 226 | 203 | 203 | -1 | -1 | 189 | 195 | 251 | 251 | 222 | 222 | 246 | 246 | 208 | 208 | 222 | 237 | 206 | 206 | -1 | -1 |
| US04 | CMW42617 | -1 | -1 | 203 | 203 | 221 | 221 | 173 | 177 | -1 | -1 | 222 | 222 | 246 | 246 | 208 | 208 | 222 | 237 | 206 | 206 | -1 | -1 |
| US01 | CMW42624 | 226 | 229 | 203 | 203 | 221 | 221 | 173 | 177 | 251 | 251 | 222 | 222 | 246 | 246 | 208 | 208 | -1 | -1 | 206 | 206 | -1 | -1 |
| US02 | CMW42626 | 226 | 229 | 200 | 210 | 221 | 221 | 189 | 195 | 248 | 251 | 222 | 222 | 246 | 246 | 205 | 225 | 222 | 237 | 206 | 264 | 174 | 186 |
| CZ01 | CMW27368 | 223 | 226 | 197 | 210 | 221 | 224 | 189 | 189 | 248 | 248 | 222 | 222 | 240 | 240 | 205 | 205 | 228 | 237 | 206 | 206 | 186 | 186 |
| CZ02 | CMW27369 | 223 | 226 | 197 | 210 | 221 | 224 | 186 | 186 | 248 | 248 | 222 | 222 | -1 | -1 | 205 | 205 | -1 | -1 | 206 | 206 | 186 | 186 |
| CZ03 | CMW27371 | 223 | 226 | 197 | 210 | 221 | 224 | 189 | 189 | 248 | 248 | 222 | 222 | 240 | 240 | 205 | 205 | 228 | 237 | 206 | 206 | 186 | 186 |
| CZ04 | CMW27374 | 223 | 226 | 197 | 210 | 221 | 224 | 189 | 189 | 248 | 248 | 222 | 222 | 240 | 240 | 205 | 205 | 228 | 237 | 206 | 206 | 186 | 186 |
| ESP01 | CMW40566 | -1 | -1 | 200 | 210 | 221 | 221 | 189 | 195 | 248 | 251 | 222 | 225 | 243 | 253 | 205 | 225 | 228 | 237 | 206 | 260 | 174 | 186 |
| ESP02 | CMW40567 | 226 | 229 | 200 | 210 | 221 | 221 | 189 | 195 | 248 | 251 | 222 | 225 | 243 | 253 | 205 | 225 | 228 | 237 | 206 | 260 | 174 | 186 |
| ESP03 | CMW46037 | 226 | 229 | 200 | 210 | 221 | 221 | 189 | 195 | 248 | 251 | 222 | 225 | 243 | 243 | 205 | 225 | -1 | -1 | 206 | 264 | 174 | 186 |
| ESP04 | CMW40568 | 226 | 229 | 200 | 210 | 221 | 221 | 189 | 195 | 248 | 251 | 222 | 225 | -1 | -1 | 205 | 225 | 228 | 237 | 206 | 264 | 174 | 186 |
| ESP05 | CMW40570 | 226 | 229 | 200 | 210 | 221 | 221 | 189 | 195 | 248 | 251 | 222 | 225 | -1 | -1 | 205 | 225 | 222 | 237 | 206 | 260 | 174 | 186 |
| ESP06 | CMW46039 | 226 | 229 | 200 | 210 | 221 | 221 | 189 | 195 | 248 | 251 | 222 | 225 | 246 | 246 | 205 | 225 | -1 | -1 | 206 | 264 | 174 | 186 |
| ESP07 | CMW42630 | -1 | -1 | 200 | 210 | 221 | 221 | 189 | 195 | 248 | 251 | 222 | 225 | 246 | 246 | 205 | 225 | 222 | 237 | 206 | 260 | 174 | 186 |
| ESP08 | CMW42632 | -1 | -1 | 200 | 210 | 221 | 221 | 189 | 195 | 248 | 251 | 222 | 225 | -1 | -1 | 205 | 225 | -1 | -1 | 206 | 264 | 174 | 186 |
| AT01 | CMW16850 | 217 | 223 | 188 | 188 | 221 | 221 | 189 | 189 | 248 | 248 | 219 | 219 | 237 | 237 | 205 | 225 | -1 | -1 | 226 | 226 | 174 | 174 |
| AT02 | CMW16854 | -1 | -1 | 188 | 188 | 221 | 221 | 186 | 186 | 248 | 248 | 219 | 219 | 237 | 237 | 205 | 225 | 222 | 237 | 226 | 226 | 170 | 170 |
| AT03 | CMW16859 | 217 | 223 | 188 | 188 | 221 | 221 | 186 | 186 | 248 | 248 | -1 | -1 | 237 | 237 | 205 | 225 | 222 | 237 | 202 | 202 | 170 | 170 |

*CMW= fungal culture collection at Forestry and Agricultural Biotechnology institute (FABI), University of Pretoria
