## Supplementary Table S6 for "The influence of biosecurity on the diversity and management of a 120-year old joint insect-fungus pest invasion"

Supplementary Table S6. The woodwasp *S. noctilio* genetic diversity based on allelic diversity.

|  | Population | N^a^ | Allele^b^ | 1-D^c^ | Hexp^d^ | Evenness^e^ |
| --- | --- | --- | --- | --- | --- | --- |
| All populations | Argentina | 82 | 4.667 | 0.324 | 0.327 | 0.578 |
|  | Australia | 440 | 2.167 | 0.064 | 0.065 | 0.427 |
|  | Chile | 30 | 3.417 | 0.355 | 0.362 | 0.692 |
|  | Europe-Pop1 | 34 | 4.750 | 0.443 | 0.451 | 0.635 |
|  | New Zealand | 86 | 2.667 | 0.140 | 0.141 | 0.468 |
|  | North America | 54 | 3.670 | 0.400 | 0.410 | 0.820 |
|  | South Africa | 100 | 2.917 | 0.224 | 0.225 | 0.554 |
|  | Europe-Pop2 (Switzerland) | 101 | 2.833 | 0.185 | 0.186 | 0.514 |
|  | Uruguay | 11 | 2.330 | 0.320 | 0.330 | 0.830 |
|  | Total | 938 | 7.667 | 0.281 | 0.281 | 0.461 |
| Australasian  sub-populations | New South Wales | 94 | 1.500 | 0.082 | 0.083 | 0.549 |
|  | Queensland | 53 | 1.357 | 0.062 | 0.063 | 0.594 |
|  | South Australia | 51 | 1.214 | 0.034 | 0.035 | 0.565 |
|  | Tasmania | 69 | 1.500 | 0.100 | 0.100 | 0.680 |
|  | Victoria | 133 | 1.571 | 0.085 | 0.086 | 0.573 |
|  | New Zealand | 61 | 1.571 | 0.123 | 0.124 | 0.615 |
|  | Total | 461 | 2.071 | 0.091 | 0.091 | 0.451 |

^a^ N: Number of insect specimen per collection site

^b^ Allele: Mean number of observed alleles per loci

^c^ 1-D: Mean number of Simpson index

^d^ Hexp: Mean number of Nei's 1978 gene diversity

^e^ Evenness: distribution of alleles abundance per population
