## Supplementary Table S7 for "The influence of biosecurity on the diversity and management of a 120-year old joint insect-fungus pest invasion"

Supplementary Table S7. *Amylostereum areolatum* genetic diversity based on allelic diversity.

| Population | N^a^ | Allele^b^ | 1-D^c^ | Hexp^d^ | Evenness^e^ |
| --- | --- | --- | --- | --- | --- |
| Australia | 84 | 2.000 | 0.459 | 0.462 | 0.946 |
| Austria | 3 | 1.820 | 0.330 | 0.390 | 0.940 |
| Czech Republic | 4 | 1.640 | 0.260 | 0.300 | 0.930 |
| New Zealand | 14 | 2.450 | 0.510 | 0.530 | 0.930 |
| North America | 6 | 3.180 | 0.520 | 0.560 | 0.770 |
| South Africa | 55 | 2.550 | 0.420 | 0.430 | 0.880 |
| Spain | 8 | 2.450 | 0.520 | 0.550 | 0.980 |
| Mass-rearing strains | 5 | 2.550 | 0.540 | 0.600 | 0.910 |
| Total | 179 | 5.640 | 0.580 | 0.580 | 0.820 |

^a^ N: Number of fungal isolates per collection site

^b^ Allele: Mean number of observed alleles per loci

^c^ 1-D: Mean number of Simpson index

^d^ Hexp: Mean number of Nei's 1978 gene diversity

^e^ Evenness: distribution of alleles abundance per population
