## Supplementary Table S8 for "The influence of biosecurity on the diversity and management of a 120-year old joint insect-fungus pest invasion"

Supplementary Table S8. Summary of population differentiation between *Sirex noctilio* populations from Australia and New Zealand. Genetic differentiation (ϕP_T_) and gene flow (Nm) between pairs of *Sirex noctilio* populations represented below and above the diagonal, respectively. Note [F_ST_ (ϕP_T_) 0 means complete sharing of genetic material and F_ST_ (ϕP_T_) 1 means no sharing].

|  | New South Wales | Queensland | | Tasmania | South Australia | Victoria | New Zealand |
| --- | --- | --- | --- | --- | --- | --- | --- |
| New South Wales | - | | 1.603 | 5.926 | 1.576 | 13.388 | 3.701 |
| Queensland | 0.135 | | - | 1.083 | 0.517 | 1.744 | 1.310 |
| Tasmania | 0.040 | | 0.188 | - | 2.525 | 8.089 | 6.866 |
| South Australia | 0.137 | | 0.326 | 0.090 | - | 2.468 | 1.529 |
| Victoria | 0.018 | | 0.125 | 0.030 | 0.092 | - | 4.185 |
| New Zealand | 0.063 | | 0.160 | 0.035 | 0.141 | 0.056 | - |
