## Supplementary figures and images for "The influence of biosecurity on the diversity and management of a 120-year old joint insect-fungus pest invasion"

### Supplementary Fig S1

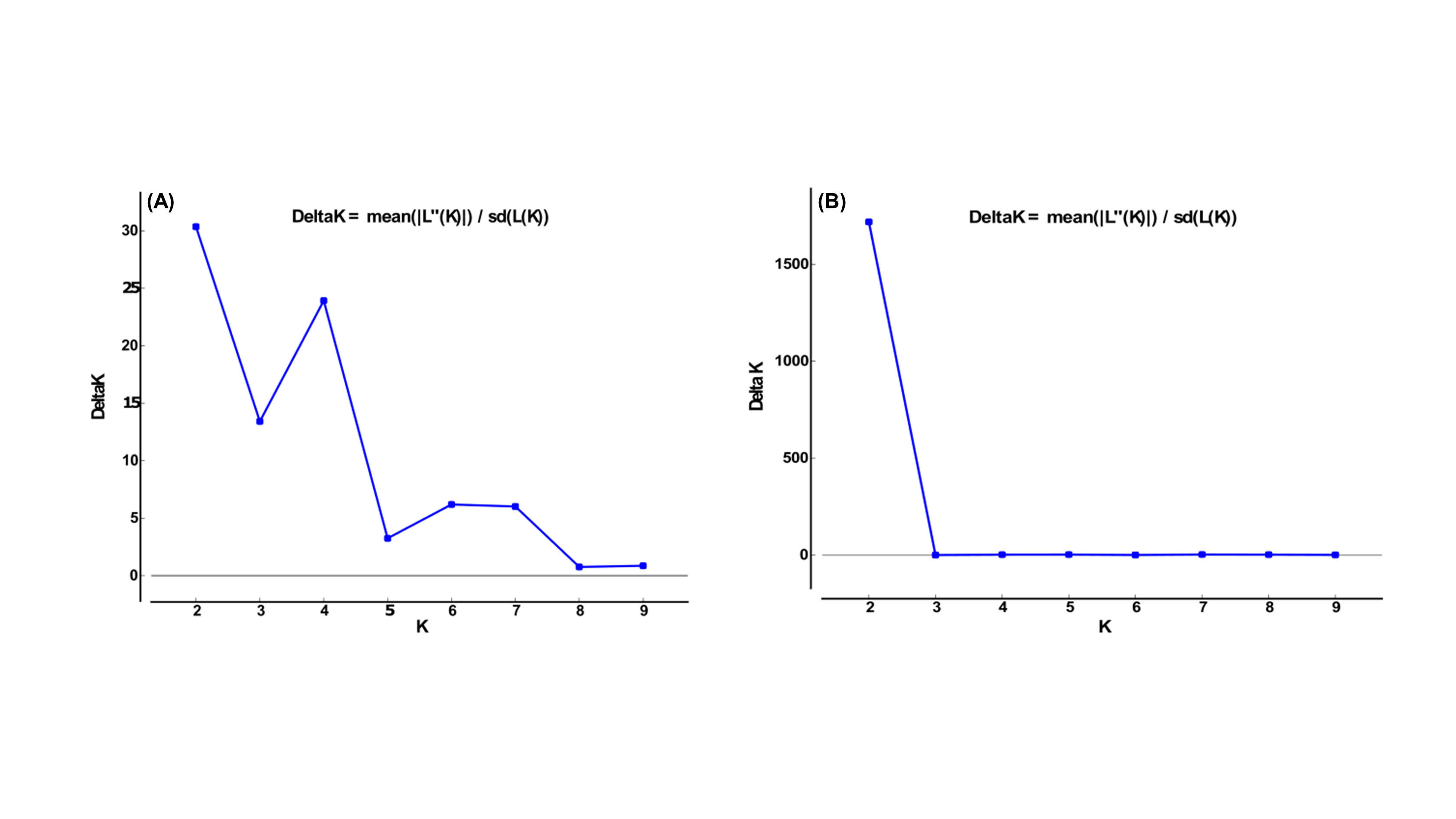

### Supplementary Fig S2

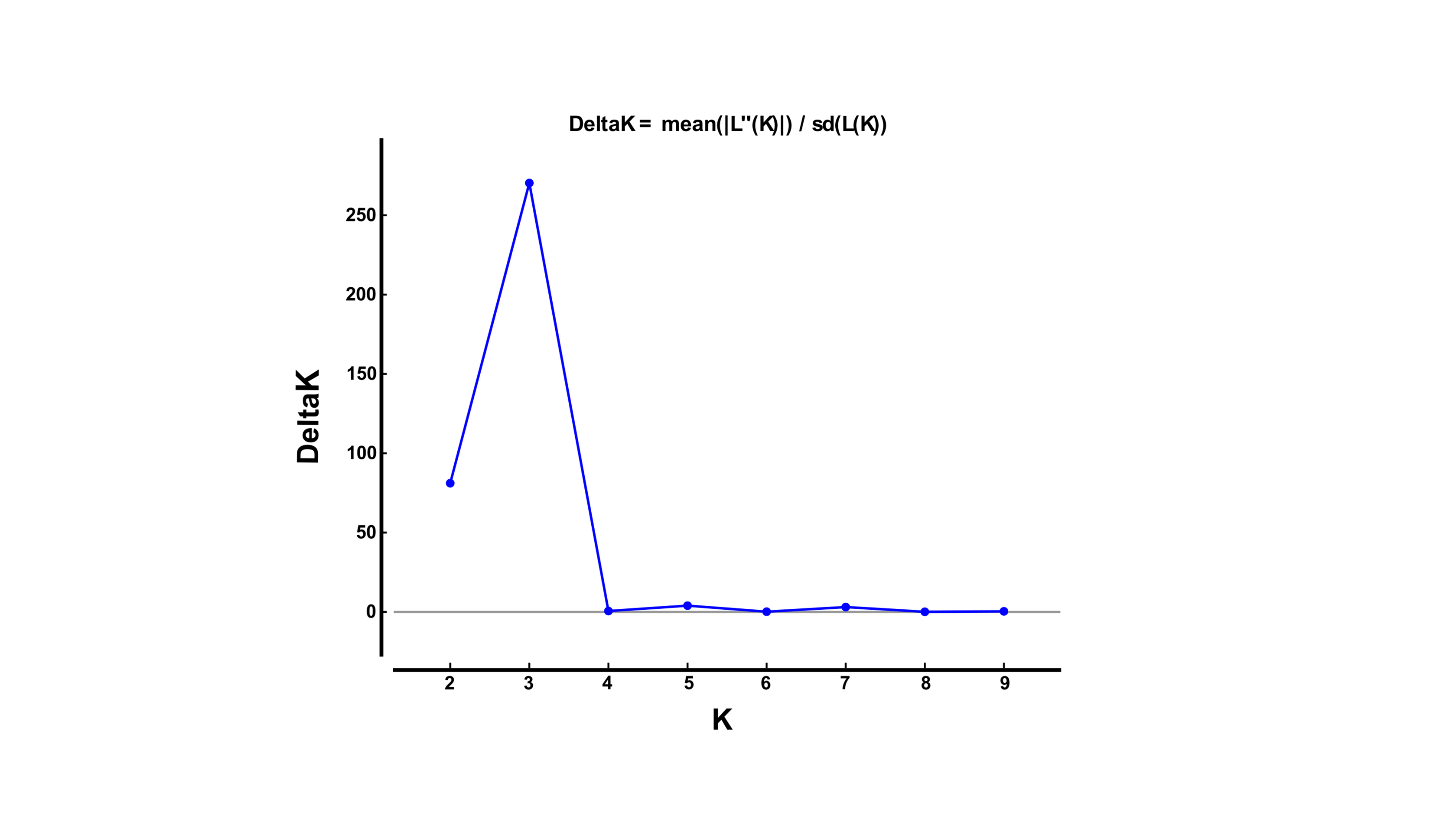

### Supplementary Fig S3

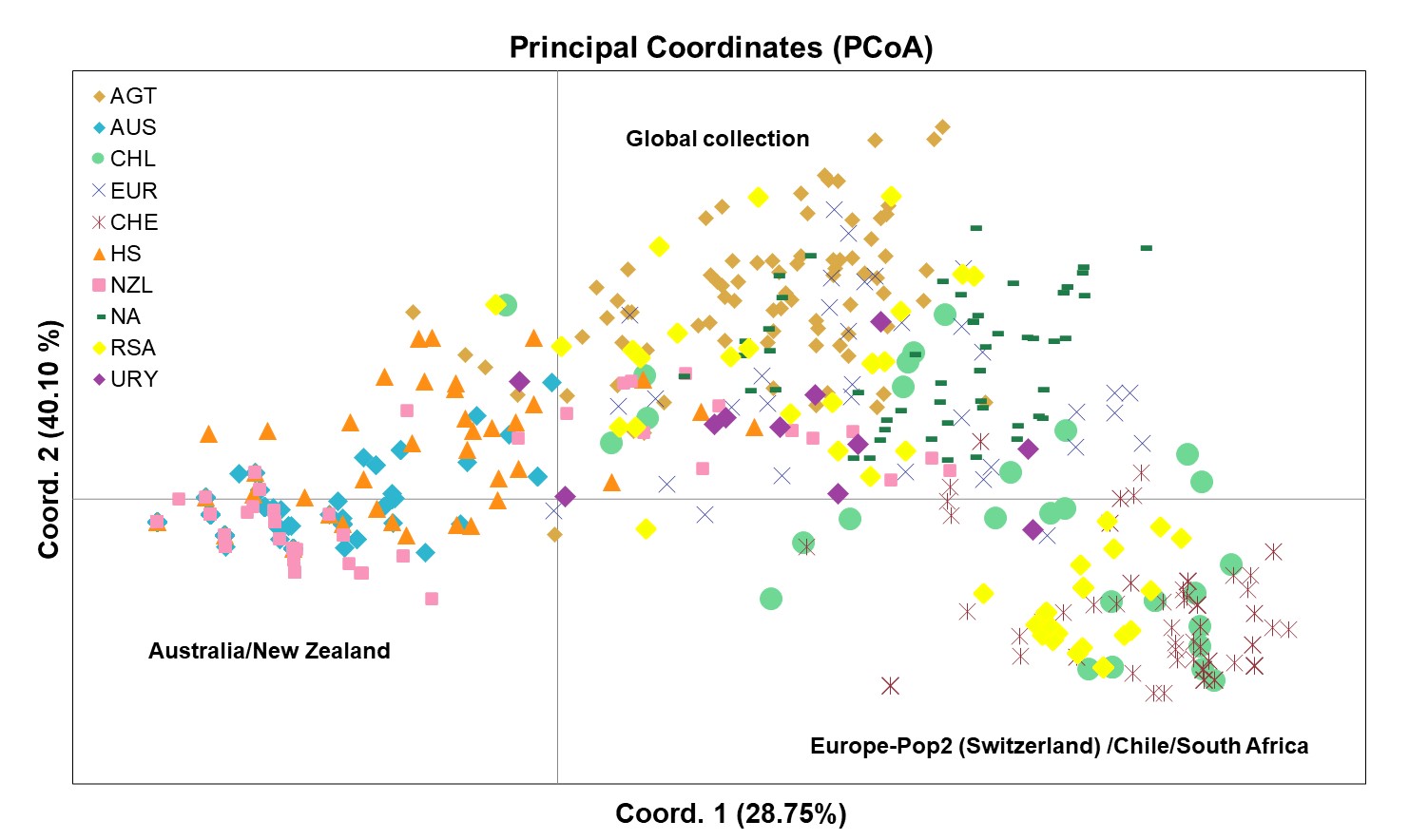

### Supplementary Fig S4

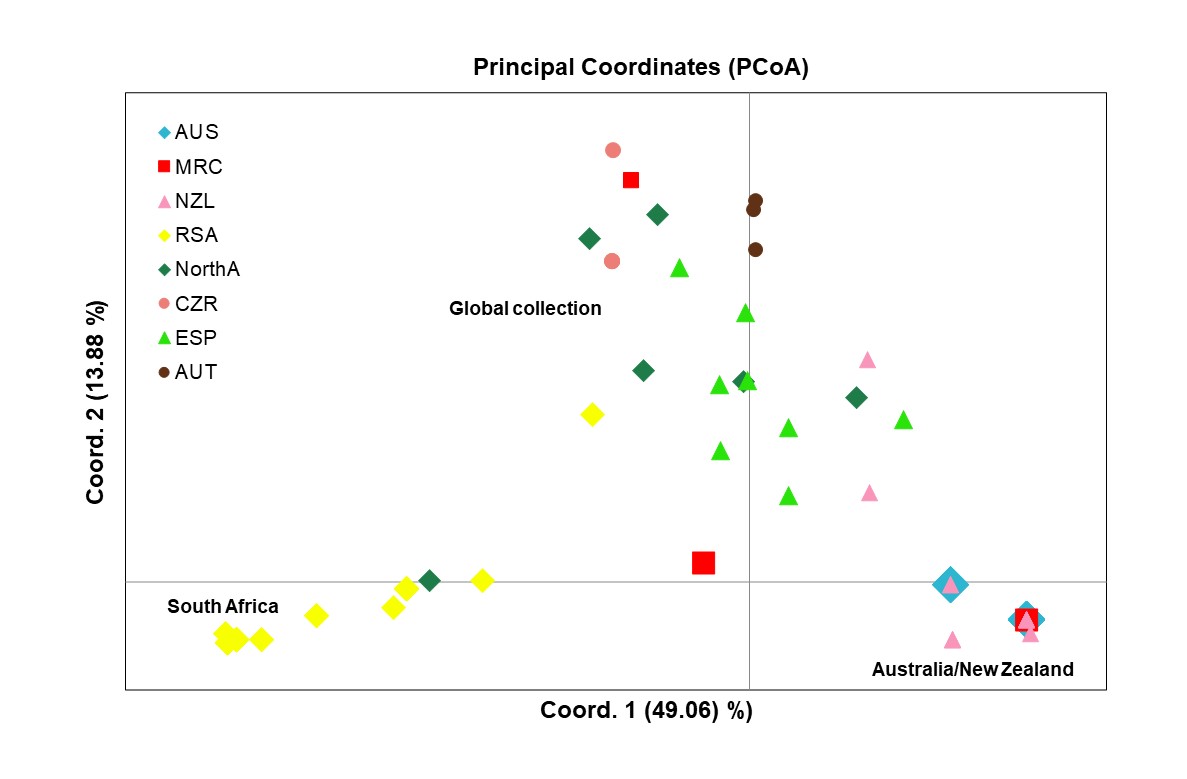
